# Genome-wide association studies in Samoans give insight into the genetic architecture of fasting serum lipid levels

**DOI:** 10.1101/411546

**Authors:** Jenna C. Carlson, Daniel E. Weeks, Nicola L. Hawley, Guangyun Sun, Hong Cheng, Take Naseri, Muagututi‘a Sefuiva Reupena, Ranjan Deka, Stephen T. McGarvey, Ryan L. Minster

**Author notes:** DEW, RD, STM, RLM are Joint Senior Authors.

## Abstract

The current understanding of the genetic architecture of lipids has largely come from genome-wide association studies. To date, few studies have examined the genetic architecture of lipids in Polynesians, and none have in Samoans, whose unique population history, including many population bottlenecks, may provide insight into the biological foundations of variation in lipid levels. Here we performed a genome-wide association study of four fasting serum lipid levels: total cholesterol (TC), high-density lipoprotein (HDL), low-density lipoprotein (LDL), and triglycerides (TG) in a sample of 2,849 Samoans, with validation genotyping for associations in a replication cohort comprising 1,798 Samoans and American Samoans. We identified multiple genome-wide significant associations (*P* < 5 × 10^−8^) previously seen in other populations – *APOA1* with TG, *CETP* with HDL, and *APOE* with TC and LDL – and several suggestive associations (*P* < 1 × 10^−5^), including an association of variants downstream of *MGAT1* and *RAB21* with HDL. However, we observed different association signals for variants near *APOE* than what has been previously reported in non-Polynesian populations. The association with several known lipid loci combined with the newly-identified associations with variants near *MGAT1* and *RAB21* suggest that while some of the genetic architecture of lipids is shared between Samoans and other populations, part of the genetic architecture may be Polynesian-specific.

## Introduction

The Samoan Islands, comprising both the U.S. Territory of American Samoa (American Samoan) and the Independent State of Samoa (Samoa), have experienced a rise in prevalence of cardiovascular disease and other non-communicable diseases in the last 30 years partly due to economic modernization, rapid urbanization, and lifestyle changes such as increased caloric intake and sedentary behavior [1–3]. A 2010 population-based survey in Samoa, which gathered the "discovery cohort" studied further here, found that many Samoans are at elevated risk of cardiovascular disease based on known risk factors—increased total cholesterol (TC), low-density lipoprotein cholesterol (LDL), and triglycerides (TG), as well as decreased high-density lipoprotein (HDL) [4–6]—with 47% of the 2,938 adult Samoans examined in the study having elevated TC (≥ 5.2 mmol/L), 88% of men and 91% of women having elevated LDL (> 2.59 mmol/L), and 43% of women having low HDL (< 1.29 mmol/L) [1].

Our current understanding of the genetic component of serum lipid level variation has been largely due to genome-wide association studies (GWAS) [7]. The Global Lipids Genetics Consortium found strong evidence for 157 loci associated with one or more of these traits using a sample of 188,577 individuals of European, East Asian, South Asian, and African ancestry [8]. However, few GWAS of serum lipid levels have been conducted in Pacific Islanders [9, 10] and to our knowledge only one has included a small number of Polynesians [11]. Previous studies have estimated the heritability of serum lipid levels in Samoans, ranging from 16% for HDL to 23% for TG, and have identified genetic susceptibility loci via linkage analysis [12], warranting further study of the genetic architecture of serum lipid levels in Samoans. Samoans are a genetically-isolated founder population, with unique evolutionary history, making them particularly useful in genomic studies [13, 14]. Thus, genomic studies of serum lipid levels could reveal novel lipid-altering loci specific to Pacific Islander populations, as well as highlight susceptibility loci shared with global populations.

Here we report the results of a GWAS of fasting TC, LDL, HDL, and TG in up to 2,849 individuals from independent Samoa followed by replication in up to 1,798 individuals from independent Samoa and American Samoa, as part of ongoing genome-wide association studies of cardiometabolic disease and adiposity-related traits in the Samoan Islands [1]. We identified multiple genome-wide significant associations previously seen in other populations – *APOA1* with TG, *CETP* with HDL, and *APOE* with TC and LDL – and several suggestive associations, including an association between variants downstream of *MGAT1* and *RAB21* with HDL.

## Methods

### Discovery Cohort and Genotyping

The discovery cohort data are available from dbGaP (accession number: phs000914.v1.p1). The discovery cohort of 2,849 individuals is drawn from a population-based sample recruited from Samoa in 2010 (Table 1). The sample selection, data collection methods, and phenotyping, including the laboratory assays for serum lipid and lipoprotein levels, have been previously reported [1, 13]. Briefly, serum lipid levels were derived from fasting whole blood samples collected after a minimum 10-hour overnight fast. Genotyping was performed using Genome-Wide Human SNP 6.0 arrays (Affymetrix). Extensive quality control was conducted on the basis of a pipeline developed by Laurie et al [15]. Additional details for sample genotyping and genotype quality control are described in Minster et al [13]. This study was approved by the institutional review board of Brown University and the Health Research Committee of the Samoa Ministry of Health. All participants gave informed consent. Imputation was not performed in this study because prior experience with this population using extant imputation panels such as the Phase 3 1000 Genomes panel showed that the resulting imputed genotypes did not correlate well with observed genotypes [13].

**Table 1.** Demographic, anthropometric, and blood biochemistry statistics of the genotyped discovery and replication cohorts.

|  | 2010 Discovery Cohort |  | 2002–2003 Replication Sample Set |  | 1994–1995 Replication Sample Set |  | 2002–2003 Replication Sample Set |  | 1994–1995 Replication Sample Set |  |
| --- | --- | --- | --- | --- | --- | --- | --- | --- | --- | --- |
|  | Samoa, n = 2849 |  | Samoa, n = 490 |  | Samoa, n = 468 |  | American Samoa, n = 592 |  | American Samoa, n = 248 |  |
|  | Women | Men | Women | Men | Women | Men | Women | Men | Women | Men |
|  | n = 1703 | n = 1146 | n = 245 | n = 245 | n = 243 | n = 225 | n = 337 | n = 255 | n = 137 | n = 111 |
| age (years) | 44.8 (11.10) | 45.6 (11.10) | 44 (17.00) | 40.7 (16.30) | 43.3 (8.60) | 43.2 (8.90) | 43 (16.00) | 43.2 (16.60) | 43.3 (9.10) | 45.5 (10.60) |
| BMI (kg/m<sup>2</sup>) | 34.8 (6.70) | 31.3 (5.90) | 33.2 (7.70) | 28.8 (5.40) | 31.9 (5.70) | 28.8 (5.00) | 36.5 (8.40) | 33.4 (7.60) | 37.4 (6.90) | 34.9 (6.00) |
| total cholesterol (mg/dL) | 199.3 (36.10) | 200.3 (38.70) | 202.3 (35.90) | 195.5 (40.60) | 199.1 (37.20) | 197.4 (35.00) | 187.1 (38.60) | 189.6 (37.80) | 197.8 (31.20) | 194.1 (31.00) |
| HDL (mg/dL) | 46.5 (10.80) | 43.7 (11.20) | 47.1 (10.30) | 46.3 (11.20) | 43.4 (11.00) | 42.5 (11.40) | 40.9 (8.80) | 40 (8.80) | 34.8 (7.60) | 32.8 (8.30) |
| LDL (mg/dL) | 130 (32.60) | 129.6 (35.30) | 130.3 (32.80) | 128.6 (37.50) | 140.2 (34.80) | 133.5 (33.00) | 118.5 (33.70) | 118.9 (35.10) | 136.6 (32.40) | 136 (37.50) |
| triglycerides (mg/dL) | 114.9 (80.50) | 139.3 (113.10) | 110.9 (58.90) | 120.1 (91.30) | 85.6 (72.80) | 102.8 (97.20) | 130.8 (78.10) | 200.2 (206.50) | 118.9 (63.40) | 168.2 (109.00) |
| fasting glucose (mg/dL) | 107.6 (56.20) | 107.4 (54.80) | 102.2 (46.40) | 98.6 (49.80) | 90.9 (38.00) | 85.3 (18.90) | 104.2 (50.30) | 108.8 (54.70) | 107.3 (49.80) | 125.2 (61.70) |

### Replication Cohort and Genotyping

The replication cohort of 1,798 individuals contains two sample sets recruited from Samoa and American Samoa (Table 1). Although there is substantial economic variation across the two polities, with American Samoans generally having a higher standard of living, the Samoans from both territories form a single socio-cultural unit with frequent exchange of mates; genetically, they represent a single homogenous population [3, 16]. The first sample set, referred to as the 1990–95 replication sample set, contains 716 unrelated individuals derived from a longitudinal study of adiposity and cardiovascular disease risk factors among adults from American Samoa and Samoa (Table 1). Detailed descriptions of the sampling and recruitment have been reported previously [17–19]. Briefly, participants were recruited from 46 villages and worksites in American Samoa in 1990 and 9 villages in Samoa (Western Samoa, at the time of recruitment) in 1991 and followed up four years later in 1994 and 1995, respectively. All participants were, at baseline, free of self-reported history of heart disease, hypertension, or diabetes. This study was approved by the institutional review board of the Miriam Hospital, Providence, RI. All participants gave informed consent. The second sample set, referred to as the 2002–03 replication sample set, contains 1,082 individuals from American Samoa and Samoa and was drawn from an extended family-based genetic linkage analysis of cardiometabolic traits (Table 1) [12, 20–23]. Probands and relatives were unselected for obesity or related phenotypes. The recruitment process and criteria used for inclusion in this study have been described in detail previously [21, 23]. This study was approved by the institutional review board of Brown University and research ethics review committees in both Samoa and American Samoa. All participants gave informed consent. Imputation was not performed in these studies for the same rationale as the discovery cohort above, but also because genome-wide marker data was not available for the samples in these studies.

In both replication sample sets, blood samples were collected in the morning after a minimum of 10 hours fasting, from which serum lipid levels were derived using assay methods published previously [12, 17]. Genotyping of variants selected for validation in the replication cohort (described below) was performed using custom-designed TaqMan OpenArray Real-Time PCR assays (Applied Biosystems). SNPs that could not be genotyped using OpenArray assays were genotyped individually using TaqMan SNP Genotyping assays (Applied Biosystems). Eight variants could not be genotyped due to technical difficulties.

### Statistical Analyses

Prior to association analyses, residuals were generated for all four lipid traits. First, traits were transformed to normality with the Box–Cox power transformation; secondly, model selection was performed using step-wise linear regression with initial model covariates previously associated with serum lipid levels: age, age^2^, sex, log-transformed BMI, fasting glucose, smoking status, farming status (as a measure of physical activity), and interactions between age, age^2^, and sex. The final TC model adjusted for age, age^2^, sex, age × sex, and age^2^ × sex; the final LDL and TG models adjusted for age, age^2^, sex, and age^2^ × sex; the final HDL model adjusted for age and sex.

Preliminary associations were performed, and variants were selected for validation without consideration of hypolipidemic medication use, as it was not measured. However, participants did self-report use of heart disease medication. Sensitivity analysis revealed that this self-reported use of medication to treat heart disease was significantly associated with TC and LDL (results not shown); individuals reporting such medication use (n = 17) were excluded from analyses. The prioritization of variants for validation genotyping was updated using these analyses, but only after available resources were fully expended. Unfortunately, not all variants that should have been prioritized for validation genotyping were successfully genotyped. All results presented are those of the corrected analyses, removing the individuals with heart disease medication use.

Additional sensitivity analysis was performed for TG by excluding one outlying observation (i.e., TG > 4 standard deviations above mean); results did not change qualitatively, and, since the recorded value was within the range of plausible values for TG, the individual was retained for presented analyses.

Association between lipid residuals and autosomal genotypes of 659,492 SNPs with minor allele frequency (MAF) ≥ 0.05 and Hardy-Weinberg Equilibrium (HWE) test *P* value ≥ 5 × 10^−5^ was assessed using linear mixed modelling in GenABEL, including previously-derived empirical kinship estimates to adjust for subject relatedness [13, 24]. The association between X-chromosome genotypes and the lipid phenotypes were calculated in GenABEL, without adjustment using the empirical kinship estimates. Genomic inflation due to population stratification and cryptic relatedness was assessed by estimating λ_GC_ using the lower 90% of the *P* value distribution [25]. GWAS *P* values in the discovery cohort (*P*_D_) were compared to a threshold for genome-wide significance of *P*_D_ < 5 × 10^−8^ and a suggestive association threshold of *P*_D_ < 1 × 10^−5^. Statistical power to detect signals at these thresholds was calculated using the Genetic Power Calculator [26].

Gene-set enrichment analysis with MAGENTA was also performed to identify any biological pathways enriched for discovery association signals [27]. Briefly, gene scores were obtained from the most significant *P* value among SNPs located within each gene using the association results from each lipid GWAS. Genes scores were adjusted for confounding factors including gene size, number of variants, and linkage disequilibrium-related properties by using step-wise multiple linear regression. The 95th percentile of all gene scores was used as the enrichment cutoff for each trait [28]. Gene-set enrichment *P* values were obtained for highly ranked gene scores. Gene sets were obtained from Gene Ontology (April 2010), pathway information from the Ingenuity (June 2008) and KEGG (June 2010), and biological processes and molecular function from PANTHER (January 2010).

For each of the lipid traits, the INRICH program [29] was used to test for enrichment of known genes (as constructed from Teslovich et al. [30] and Willer et al [8]). INRICH tests if more known genes are contained in associated intervals than expected by chance, using permutation based on 1 million replicates to generate experiment-wide empirical *P* values. For each lipid trait, we defined the associated intervals as 100 kb intervals centered on the most significant SNP within association peaks with *P*_*D*_ < 1 × 10^−4^.

We selected 21 regions demonstrating at least suggestive association for association validation in the replication cohort. An additional 10 regions which should have selected for validation were not followed-up because their exclusion was based on preliminary analyses that included 17 participants taking heart disease medication—participants who were ultimately excluded from these studies. The variant from each locus with smallest *P* value across the four lipid scans (defined as 1 Mb windows surrounding the peak SNP) or a proxy SNP in high linkage disequilibrium with the lowest-*P* value SNP was selected as representative of the locus for replication genotyping.

Statistical association was measured in the 1990–95 and 2002–03 replication sample sets independently, and results were combined using meta-analysis (see below). Association analyses for both sample sets were performed using GenABEL [31] in R [32], using the same regression models as in the discovery cohort but additionally adjusting for polity (American Samoa or Samoa); the 2002-03 sample set was additionally adjusted using expected kinship, as derived from familial pedigree information [33].

Prior to meta-analysis, quality control was performed using EasyQC to check for strand and allele frequency consistency [34]. *P* value-based meta-analysis using sample sizes as weights was performed using METAL [35] to generate two *P* values: one for the meta-analysis of the two replication cohorts (*P*_R_) and one for the replication cohorts and discovery sample together (*P*_DR_). Resulting meta-analysis signals were evaluated based on genome-wide significance and suggestive thresholds (as described above) and by the contribution of the replication sample to the signal. Effect directions for meta-analysis results of peak SNPs were qualitatively compared to those of previously reported lead SNPs.

For ease of reference, any locus identified here with a corresponding signal within 1 mega base pairs (Mb) in a prior lipid study is referred to by the previously prescribed locus name [8, 10]; for loci not previously associated with lipid traits, the symbol of the gene nearest the peak SNP in the locus or the hyphen-separated symbols of the nearest two genes is used as the locus label.

## Results

The demographic, anthropometric, and biochemical characteristics of the 2,849 participants composing the discovery cohort for this GWAS of serum lipids levels and the 1,798 participants composing the replication cohort are presented in Table 1. A detailed description of the discovery cohort and its trends compared to the historical sample sets making up the replication cohort has been previously reported [1]. Briefly, the average age was similar for all cohorts; average BMI was higher among women compared to men and in American Samoa compared to Samoa; average BMI for men and women in Samoa was higher in more recent studies; average lipid levels are largely similar across cohorts with minor exceptions.

We assessed 659,492 unique genome-wide markers for association with 4 traits—TC, HDL, LDL and TG—in up to 2,849 Samoans in the discovery cohort. Relatedness within the discovery cohort was well-controlled using the empirical kinship coefficients; λ_GC_ ranged between 1.03 and 1.07 for the four lipid traits (Figs S2, S4, S6, and S8 in S1 Appendix). We observed 38 genome-wide suggestive or significant associations across 31 loci from the four GWAS (Fig 1; Figs S1, S3, S5, and S7 in S1 Appendix; Tables S1, S7, S10, and S13 in S1 Appendix).

**Fig 1.**
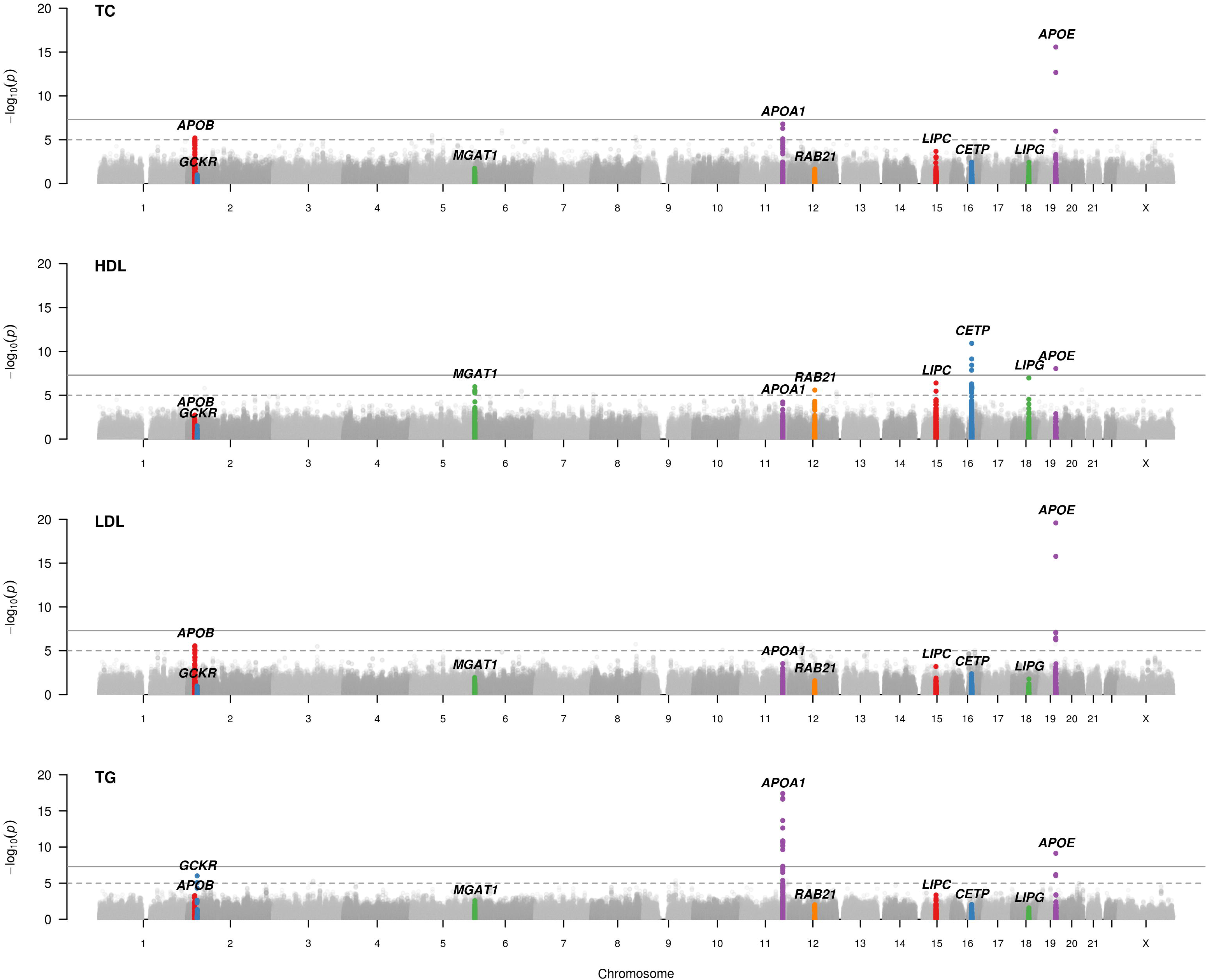
Manhattan plots for GWAS of four lipid traits in the discovery cohort of 2,849 Samoans. The dashed and solid lines denote genome-wide suggestive and genome-wide significant *P* value thresholds (*P* < 1 × 10^−5^ and *P* < 5 × 10^−8^, respectively). Peaks are labeled with the candidate gene or closest gene in the region if they have at least suggestive association in the discovery cohort for at least one trait and demonstrate evidence of replication or have been previously associated.

Genome-wide significant association was observed in the discovery cohort between all four traits and markers near *APOE*: TC and rs4420638 (*P*_D_ = 2.67 × 10^−16^, Fig 2E); HDL and rs4420638 (*P*_D_ = 9.07 × 10^−9^, Fig 2F); LDL and rs1160985 (*P*_D_ = 2.61 × 10^−20^; Fig 2G); and TG and rs4420638 (*P*_D_ = 7.44 × 10^−10^, Fig 2H). Additionally, HDL was associated with markers near *CETP* (rs289708, *P*_D_ = 1.19 × 10^−11^), and TG, with *APOA1* (rs6589566, *P*_D_ = 3.98 × 10^−18^, Fig 2C). Suggestive associations were observed between lipid levels and markers at an additional 28 loci, including the *MGAT1* and *RAB21* loci and HDL (Fig 2A,D), and *APOA1* with TC (rs3741298, *P*_D_ = 1.63 × 10^−7^, Fig 2B). We had 80% power at *ɑ* = 1 × 10^−5^ and *ɑ* = 5 × 10^−8^ to detect SNPs that account for 1.0% and 1.5%, respectively, of the residual variance in a phenotype.

**Fig 2.**
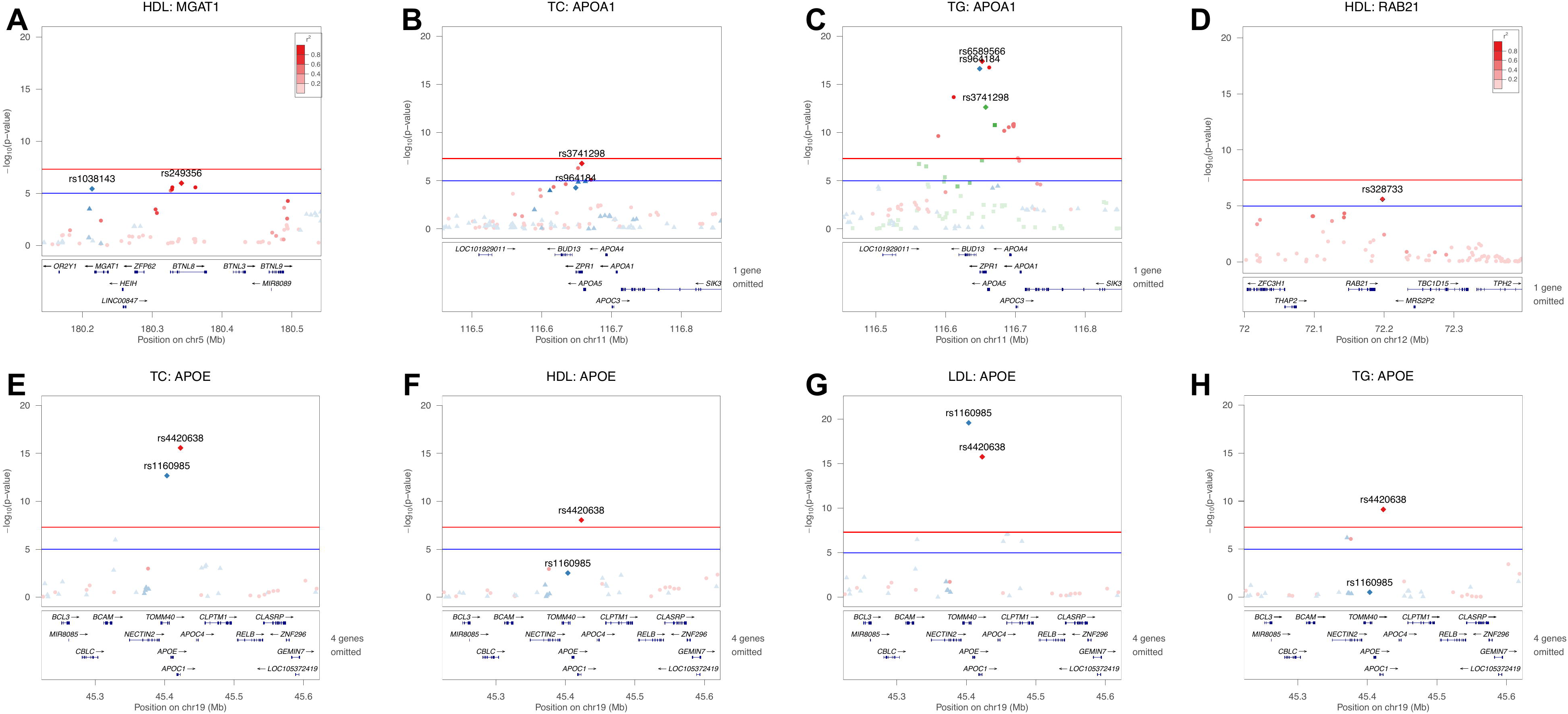
Regional association plots for selected loci. Regional association plots generated in LocusZoom [36] showing –log10(*P* values) for SNPs in the (A) *MGAT1* locus and HDL, (B) *APOA1* locus and TC, (C) *APOA1* locus and TG, (D) *RAB21* locus and HDL, and the *APOE* locus and (E) TC, (F) HDL, (G) LDL, and (H) TG. Points are color coded within each plot according to pairwise linkage disequilibrium (*r^2^*) with the labeled SNPs; the saturation of the color of each plotted SNP measures the linkage disequilibrium (*r^2^*) with the labeled SNP sharing the same color.

Gene-set enrichment analysis with MAGENTA highlighted, at a < 5% false-discovery rate (FDR), several lipid homeostasis pathways and gene ontologies for HDL and TG (Tables S8 and S14 in S1 Appendix). Four gene sets were below the FDR for both HDL and TG: HDL particle remodeling, reverse cholesterol transport, cholesterol efflux, and phospholipid efflux. An additional 12 gene sets were implicated for HDL and three gene sets for TG. The HDL particle remodeling and reverse cholesterol transport gene sets had significant enrichment for TC (Table S3 in S1 Appendix), and a single gene set was implicated with LDL, the amylase pathway (Table S11 in S1 Appendix). All four traits had significant enrichment for known TC, HDL, LDL, and TG loci using the INRICH method (Tables S5, S9, S12, and S15 in S1 Appendix).

Validation of peak SNPs was attempted for 21 loci. At loci with multiple associated variants, the most significant variant was chosen as representative of the locus. For some loci, the exclusion of participants using self-reported heart disease medication resulted in a different peak SNP. Thus, for the *APOE* locus rs1160985 was genotyped instead of rs4420638 (*P*_D_ = 2.67 × 10^−16^ for TC, *P*_D_ = 9.07× 10^−9^ for HDL, and *P*_D_ = 7.44 × 10^−10^ for TG); for the *APOA1* locus rs964184 was genotyped instead of rs3741298 (*P*_D_ = 1.63 × 10^−7^ for TC) or rs6589566 (*P*_D_ = 3.98 × 10^−18^ for TG); for the *MGAT1* locus rs1038143 was genotyped instead of rs249356 (*P*_D_ = 1.06 × 10^−6^ for HDL); for the *APOB* locus rs754523 was genotyped instead of rs1469513 (*P*_D_ = 2.71 × 10^−6^ for LDL).

We successfully genotyped the peak SNP, or a proxy SNP, in the replication cohorts for 15 loci. Two loci (*APOA1* with TG, *P*_DR_ = 1.81 × 10^−29^; *APOE* with TC, *P*_DR_ = 4.29 × 10^−21^, and LDL, *P*_DR_ = 1.53 × 10^−27^) demonstrated genome-wide significant associations in the discovery-replication meta-analysis (Table 2 and Tables S1, S7, S10, and S13 in S1 Appendix). An additional four associations demonstrated evidence of replication with consistent directions of effect and suggestive joint *P*_DR_ values (*GCKR* with TG, *P*_DR_ = 5.62 × 10^−8^; *MGAT1* with HDL, *P*_DR_ = 2.91 × 10^−7^; *APOA1* with TC, *P*_DR_ = 1.72 × 10^−6^; *RAB21* with HDL, *P*_DR_ = 5.92 × 10^−7^). Three associations had suggestive joint *P*_DR_ values driven by the discovery associations only (*APOB* with LDL, *P*_DR_ = 5.81 × 10^−6^; *LIPC* with HDL, *P*_DR_ = 9.15 × 10^−7^; *CDH4* with HDL, *P*_DR_ = 8.77 × 10^−6^); associations at *APOB* and *CDH4* had consistent directions of effect. Among the remaining loci with at least suggestive association in the discovery sample, but not in the discovery-replication meta-analysis, consistent effect directions were also seen for TC and *APOB* and *ZHX2*; LDL and *ALG10* and *CPNE8* (Table 2 and Tables S1, S7, S10, and S13 in S1 Appendix).

**Table 2.**
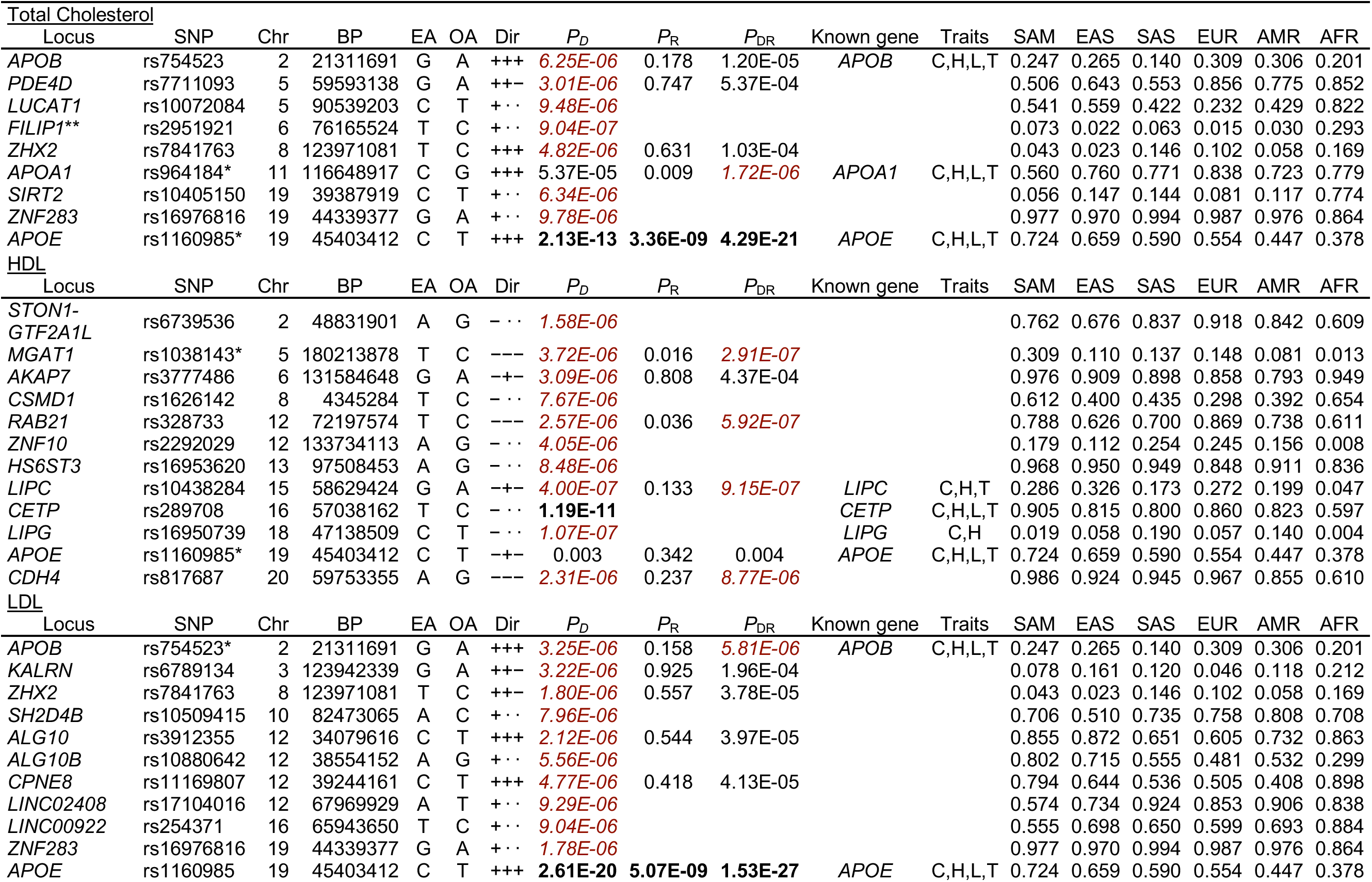
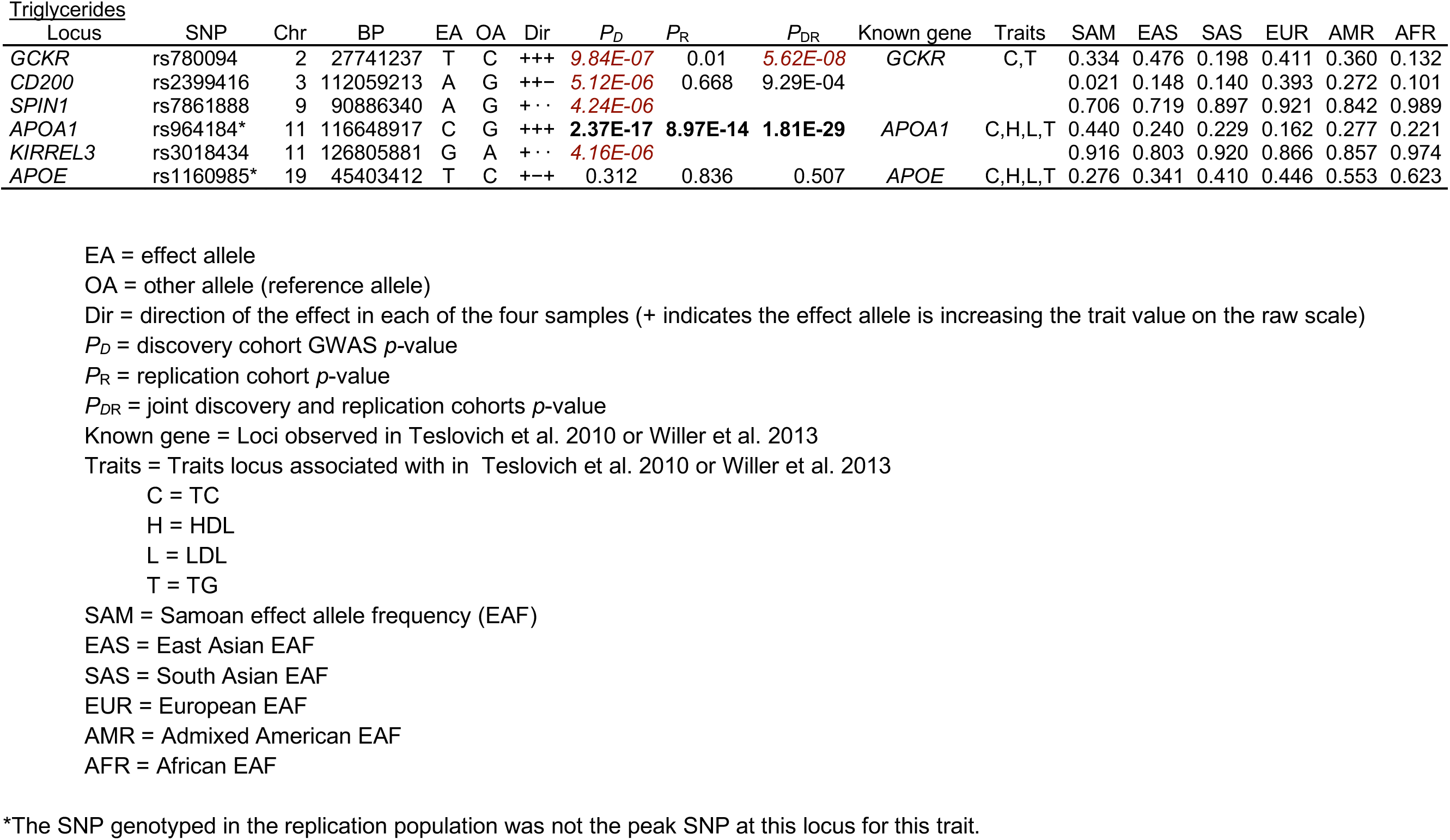
Suggestive loci and replication genotyping

We compared the directions of effect to those previously reported in Willer et al. [8] and Teslovich et al. [30] for *APOB*, *GCKR*, *APOA1*, *LIPC*, *LIPG*, and *APOE* (i.e., genome-wide suggestive loci that have been previously associated with lipid traits). We observed a consistent direction of effect for the representative SNP for all associations except for *LIPG* and HDL (Table 2).

The effect allele frequencies in the two samples—discovery and replication— were largely similar for each of the 15 successfully genotyped SNPs (Tables S1, S7, S10, and S13 Appendix). However, many loci had markedly different effect allele frequencies between Samoans and other 1000 Genomes populations (Table 2). For example, compared to 1000 Genomes populations, there were higher effect allele frequencies (EAFs) in Samoans for rs964184 near *APOA1* (G allele frequency: 0.440 in Samoans vs. < 0.277 in 1000 Genomes populations), rs1160985 near *APOE* (C allele frequency: 0.724 in Samoans vs. < 0.659 in 1000 Genomes populations), and rs1038143 near *MGAT1* (A allele frequency: 0.309 in Samoans vs. < 0.148 in 1000 Genomes populations).

## Discussion

In this study, we examined four measures of fasting lipid levels—TC, HDL, LDL and TG—for associations with 659,492 SNPs from a genome-wide array in a discovery cohort of 2,849 Samoans, with follow-up genotyping of significant and suggestive findings in a replication cohort comprising 1,798 Samoans from Samoa and American Samoa. Thirty-one loci had at least suggestive evidence of association with one or more lipid traits in the discovery cohort, of which eight have been reported to be associated with lipid levels previously: *APOB, GCKR, MGAT1, APOA1*, *LIPC*, *CETP, LIPG*, and *APOE* [8, 10, 30] although the direction of effect for the variant near LIPG was in the opposite direction from previous results. Enrichment analyses highlighted known lipid metabolism gene sets and previously associated lipid loci.

We observed a difference in the architecture of the statistical association signals between the four lipid traits and variants near *APOE* (Fig 2E-H). The peak SNP for TC and LDL was rs1160985, an intronic variant in *TOMM40* upstream of *APOE*; whereas the peak SNP for HDL and TG was rs4420638, an intergenic variant downstream of *APOC1* and *APOE*. rs1160985 demonstrated evidence of replication for TC and LDL but not for HDL and TG, consistent with the discovery findings. These markers are in low linkage disequilibrium with each other (*r^2^* = 0.093) and may represent distinct association signals. While this could support a shared genetic architecture for TC and LDL and for HDL and TG in Polynesians, this study was not positioned to adequately capture the association signal present at this locus. Future studies with sequencing or imputation of the 19q13.2 region will be necessary to dissect the genetic architecture of *APOE* and lipid levels in Polynesians.

We did not observe suggestive or genome-wide significant association with several loci which have figured prominently in multiple lipid GWAS (e.g., *LPL*, *LDLR*, *CILP2*, *FADS1*/*2*/*3*, *ANGPTL3*, *SORT1*, *PPP1R3B*, *MIXIPL*, *HNF4A*, *PCSK9*, *GALNT2*, *HMGCR*) either because we lacked sufficient power to detect their effects, the effects are negligible in Samoans, or the allele frequencies of associated variants are different enough in Samoans to hinder detection. However, it is important to note that this study was not designed to evaluate the effect of known lipids loci in Samoans, nor were previously-associated loci examined specifically.

We detected and replicated a suggestive association between HDL and a variant on 5q35.3 (Fig 2A). While the peak SNP lies within an intron of *BTNL8*, the variant selected for follow-up genotyping is intergenic, downstream of *MGAT1*. A suggestive association between variants near *MGAT1* and HDL in a GWAS in the Micronesian population of Kosrae has been previously reported [10]. Although there is no evidence of association between *MGAT1* and lipids as reported in prior studies of non-Pacific Islanders, variation near *MGAT1* has been associated with BMI, serum fatty acid levels and composition, and glucose response in Europeans [37–39]. The encoded MGAT enzyme plays a major role in the absorption of dietary fat in the intestine [40]. Due to the greater frequency of the HDL-associated risk variant observed in Samoans compared to 1000 Genomes populations, it is plausible that variation near *MGAT1* may have a unique role in or a stronger effect on the lipid metabolism of Pacific Islanders.

We also detected and replicated a novel suggestive association between HDL and a variant downstream of *RAB21* (Fig 2D). Unlike the variant downstream of *MGAT1*, we observed similar allele frequencies between Samoans and 1000 Genomes populations (i.e., < 20% difference between Samoans and another 1000 Genomes population) in the HDL-associated variant downstream of *RAB21*. This region, 12q21, was previously seen in linkage analysis with both univariate and bivariate scans of TC and LDL [12]. The individuals included in this linkage analysis are also included in our 2002–03 replication sample set, however, they do not appear to be driving the association signal near *RAB21* (Table S7 in S1 Appendix). Variation near *RAB21* has been previously associated with obesity [41]. *RAB21* belongs to the family of monomeric GTPases involved in control of cellular membrane trafficking and is involved in the targeted trafficking of integrins and the regulation of cell adhesion and migration [42, 43].

This study is limited in drawing conclusions about the genetic architecture of lipids in Samoans, as replication genotyping was unavailable for many loci and due to the lack of genome-wide imputation. Future studies, evaluating the evidence of association between associations seen here in separate cohorts as well fine-mapping loci with genotype imputation (given the availability of a relevant reference panel), are necessary to fully evaluate the genetic architecture of lipids in Samoans.

This is the first GWAS of lipid phenotypes in Samoans, and we observed association with many known lipid loci, which was further supported by the gene-set enrichment analysis highlighting lipid metabolism gene sets. However, the difference in association results near *APOE*, coupled with evidence of Pacific-Islander–specific associations with *MGAT1* and *RAB21* suggest that some, but not all, of the genetic architecture of lipids is shared between Samoans and other populations. Given this evidence of a partially distinct genetic architecture of lipids in Samoans, further investigation and fine-mapping of lipid loci, especially that across multiple ethnicities, is warranted.

## Acknowledgements

The authors would like to thank the Samoan participants of the study, local village authorities, and the many Samoan and other field workers over the years. We acknowledge the Samoan Ministry of Health, the Samoa Bureau of Statistics, and the American Samoan Department of Health for their support of this research. We give particular thanks to two research assistants, Melania Selu and Vaimoana Lupematasila, who contributed to the 2010 recruitment and continue to assist us in our work in Samoa.

## Supporting Information

**S1 Appendix. Supporting Information.**

## Data Availability Statement

The discovery cohort data are available from dbGaP (accession number: phs000914.v1.p1).

