## Supplementary material for "Genome-wide association studies in Samoans give insight into the genetic architecture of fasting serum lipid levels": S1 Appendix. Supporting Information

### Contents

1. Author Contributions 2
2. Total Cholesterol (TC) 3
3. HDL 10
4. LDL 17
5. Triglycerides (TG) 23
6. LocusZoom plots 28
6.1. Total Cholesterol LocusZoom plots 28
6.2. HDL LocusZoom plots 37
6.3. LDL LocusZoom plots 49
6.4. Triglycerides LocusZoom plots 60

### List of Tables

S1 TC results 4
S2 Descriptions of the columns in the GWAS results tables. 5
S3 MAGENTA pathway analysis of TC. 7
S4 Descriptions of the columns in the MAGENTA results tables. 7
S5 INRICH analysis of TC 8
S6 INRICH parameter settings. 9
S7 HDL results 11
S8 MAGENTA pathway analysis of HDL 14
S9 INRICH analysis of HDL 16
S10 LDL results 18
S11 MAGENTA pathway analysis of LDL 21
S12 INRICH analysis of LDL 22
S13 TG results 24
S14 MAGENTA pathway analysis of TG 26
S15 INRICH analysis of TG 27

### List of Figures

S1 TC: genome-wide association scan. 3
S2 TC: QQ plot 3
S3 HDL: genome-wide association scan 10
S4 HDL: QQ plot 10
S5 LDL: genome-wide association scan 17
S6 LDL: QQ plot. 17
S7 TG: genome-wide association scan. 23
S8 TG: QQ plot 23
S9 TC on chromosome 2 positions 21111691-21511691 28

S10 TC on chromosome 5 positions 59393138-59793138 . . . . . . . . . . . . . . . . . . 29
S11 TC on chromosome 5 positions 90339203-90739203 . . . . . . . . . . . . . . . . . . 30
S12 TC on chromosome 6 positions 75965524-76365524 . . . . . . . . . . . . . . . . . . 31
S13 TC on chromosome 8 positions 123771081-124171081 . . . . . . . . . . . . . . . . . 32
S14 TC on chromosome 11 positions 116457561-116857561. . . . . . . . . . . . . . . . . 33
S15 TC on chromosome 19 positions 39187919-39587919 . . . . . . . . . . . . . . . . . 34
S16 TC on chromosome 19 positions 44139377-44539377 . . . . . . . . . . . . . . . . . 35
S17 TC on chromosome 19 positions 45222946-45622946 . . . . . . . . . . . . . . . . . 36
S18 HDL on chromosome 2 positions 48631901-49031901 . . . . . . . . . . . . . . . . . 37
S19 HDL on chromosome 5 positions 180141895-180541895 . . . . . . . . . . . . . . . . 38
S20 HDL on chromosome 6 positions 131384648-131784648 . . . . . . . . . . . . . . . . 39
S21 HDL on chromosome 8 positions 4145284-4545284 . . . . . . . . . . . . . . . . . . 40
S22 HDL on chromosome 12 positions 71997574-72397574 . . . . . . . . . . . . . . . . . 41
S23 HDL on chromosome 12 positions 133534113-133934113 . . . . . . . . . . . . . . . . 42
S24 HDL on chromosome 13 positions 97308453-97708453 . . . . . . . . . . . . . . . . . 43
S25 HDL on chromosome 15 positions 58429424-58829424 . . . . . . . . . . . . . . . . . 44
S26 HDL on chromosome 16 positions 56838162-57238162 . . . . . . . . . . . . . . . . . 45
S27 HDL on chromosome 18 positions 46938509-47338509 . . . . . . . . . . . . . . . . . 46
S28 HDL on chromosome 19 positions 45222946-45622946 . . . . . . . . . . . . . . . . . 47
S29 HDL on chromosome 20 positions 59553355-59953355 . . . . . . . . . . . . . . . . . 48
S30 LDL on chromosome 2 positions 21059562-21459562 . . . . . . . . . . . . . . . . . 49
S31 LDL on chromosome 3 positions 123742339-124142339. . . . . . . . . . . . . . . . . 50
S32 LDL on chromosome 8 positions 123771081-124171081. . . . . . . . . . . . . . . . . 51
S33 LDL on chromosome 10 positions 82273065-82673065 . . . . . . . . . . . . . . . . . 52
S34 LDL on chromosome 12 positions 33879616-34279616 . . . . . . . . . . . . . . . . . 53
S35 LDL on chromosome 12 positions 38354152-38754152 . . . . . . . . . . . . . . . . . 54
S36 LDL on chromosome 12 positions 39044161-39444161 . . . . . . . . . . . . . . . . . 55
S37 LDL on chromosome 12 positions 67769929-68169929 . . . . . . . . . . . . . . . . . 56
S38 LDL on chromosome 16 positions 65743650-66143650 . . . . . . . . . . . . . . . . . 57
S39 LDL on chromosome 19 positions 44139377-44539377 . . . . . . . . . . . . . . . . . 58
S40 LDL on chromosome 19 positions 45203412-45603412 . . . . . . . . . . . . . . . . . 59
S41 TG on chromosome 2 positions 27541237-27941237 . . . . . . . . . . . . . . . . . . 60
S42 TG on chromosome 3 positions 111859213-112259213 . . . . . . . . . . . . . . . . . 61
S43 TG on chromosome 9 positions 90686340-91086340 . . . . . . . . . . . . . . . . . . 62
S44 TG on chromosome 11 positions 116452423-116852423. . . . . . . . . . . . . . . . . 63
S45 TG on chromosome 11 positions 126605881-127005881. . . . . . . . . . . . . . . . . 64
S46 TG on chromosome 19 positions 45222946-45622946 . . . . . . . . . . . . . . . . . 65

### 1. Author Contributions

R.L.M. performed the genotype quality control and association analyses, with guidance from D.E.W.; J.C.C. created the figures and wrote the relevant sections of the manuscript with guidance from D.E.W, R.L.M, S.T.M, and N.L.H; D.E.W. carried out the MAGENTA and INRICH analyses and assembled the Supplementary Information; N.L.H. led the field work data collection and phenotype analyses with guidance from S.T.M. G.S. led and directed genotyping experiments (using the Affymetrix 6.0 chip) and assay development for validation and replication (using the TaqMan platform) with guidance from R.D. H.C. participated extensively in DNA extraction, genotyping, and quality control of the data under the supervision of G.S. and R.D. M.S.R. facilitated fieldwork in Samoa and American Samoa. T.N. contributed to the discussion of the public health implications of the findings. All authors contributed to this work, discussed the results, and critically reviewed and revised the manuscript.

### 2. Total Cholesterol (TC)

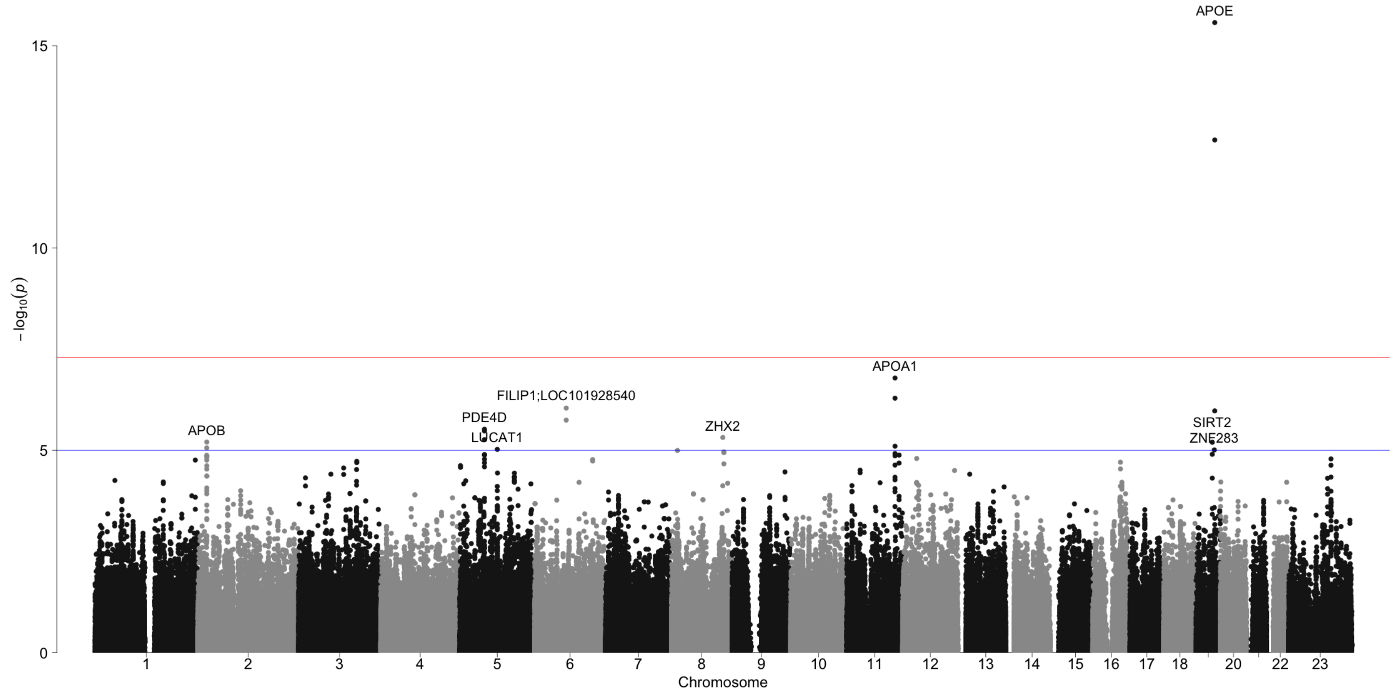


Figure S1. TC: genome-wide association scan

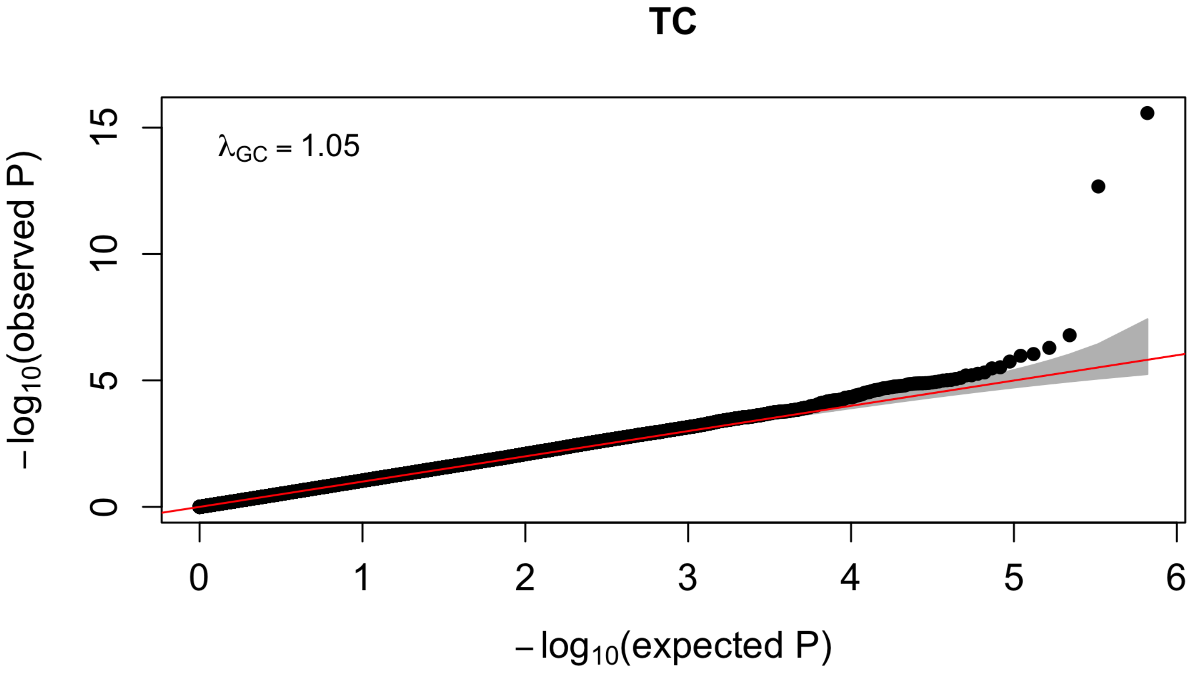


Figure S2. TC: QQ plot

Table S1. TC results. See Table S2 for a description of the column contents. Significant P values less than 5 x $10^{-8}$ are in **bold**, while suggestive P values between 1 x $10^{-5}$ and 5 x $10^{-8}$ are in *dark red italics*.

| Label | SNP | CHR | POS | $P_D$ | $P_U$ | $P_A$ | $P_R$ | $P_{DR}$ |
|---|---|---|---|---|---|---|---|---|
| APOE | rs1160985 | 19 | 45403412 | **2.13E-13** | 8.74E-05 | *9.34E-06* | **3.36E-09** | **4.29E-21** |
| APOA1 | rs964184 | 11 | 116648917 | 5.37E-05 | 0.007 | 0.237 | 0.009 | *1.72E-06* |
| APOB | rs754523 | 2 | 21311691 | *6.25E-06* | 0.116 | 0.647 | 0.178 | 1.20E-05 |
| ZHX2 | rs7841763 | 8 | 123971081 | *4.82E-06* | 0.517 | 0.925 | 0.631 | 1.03E-04 |
| PDE4D | rs7711093 | 5 | 59593138 | *3.01E-06* | 0.810 | 0.542 | 0.747 | 5.37E-04 |
| FILIP1;LOC101928540 | rs2951921 | 6 | 76165524 | *9.04E-07* | NA | NA | NA | NA |
| SIRT2 | rs10405150 | 19 | 39387919 | *6.34E-06* | NA | NA | NA | NA |
| LUCAT1 | rs10072084 | 5 | 90539203 | *9.48E-06* | NA | NA | NA | NA |
| ZNF283 | rs16976816 | 19 | 44339377 | *9.78E-06* | NA | NA | NA | NA |

| Label | SNP | CHR | POS | $z_D$ | $z_U$ | $z_A$ | Dir | $n_D$ | $n_U$ | $n_A$ | $n_R$ | $n_{DR}$ |
|---|---|---|---|---|---|---|---|---|---|---|---|---|
| APOE | rs1160985 | 19 | 45403412 | 7.34 | 3.92 | 4.43 | +++ | 2849 | 704 | 1077 | 1781 | 4630 |
| APOA1 | rs964184 | 11 | 116648917 | 4.04 | 2.68 | 1.18 | +++ | 2849 | 710 | 1080 | 1790 | 4639 |
| APOB | rs754523 | 2 | 21311691 | 4.52 | 1.57 | 0.46 | +++ | 2848 | 709 | 1078 | 1787 | 4635 |
| ZHX2 | rs7841763 | 8 | 123971081 | 4.57 | 0.65 | 0.09 | +++ | 2849 | 708 | 1078 | 1786 | 4635 |
| PDE4D | rs7711093 | 5 | 59593138 | 4.67 | 0.24 | -0.61 | ++- | 2849 | 706 | 1079 | 1785 | 4634 |
| FILIP1;LOC101928540 | rs2951921 | 6 | 76165524 | 4.91 | NA | NA | +?? | 2849 | NA | NA | NA | NA |
| SIRT2 | rs10405150 | 19 | 39387919 | 4.52 | NA | NA | +?? | 2849 | NA | NA | NA | NA |
| LUCAT1 | rs10072084 | 5 | 90539203 | 4.43 | NA | NA | +?? | 2849 | NA | NA | NA | NA |
| ZNF283 | rs16976816 | 19 | 44339377 | 4.42 | NA | NA | +?? | 2849 | NA | NA | NA | NA |

| Label | SNP | CHR | POS | eafD | eafU | eafA | SAM | EAS | SAS | EUR | AMR | AFR | eA | oA |
|---|---|---|---|---|---|---|---|---|---|---|---|---|---|---|
| APOE | rs1160985 | 19 | 45403412 | 0.732 | 0.712 | 0.709 | 0.724 | 0.659 | 0.590 | 0.554 | 0.447 | 0.378 | C | T |
| APOA1 | rs964184 | 11 | 116648917 | 0.430 | 0.453 | 0.457 | 0.440 | 0.240 | 0.229 | 0.162 | 0.277 | 0.221 | G | C |
| APOB | rs754523 | 2 | 21311691 | 0.254 | 0.248 | 0.228 | 0.247 | 0.265 | 0.140 | 0.309 | 0.306 | 0.201 | G | A |
| ZHX2 | rs7841763 | 8 | 123971081 | 0.040 | 0.047 | 0.047 | 0.043 | 0.023 | 0.146 | 0.102 | 0.058 | 0.169 | T | C |
| PDE4D | rs7711093 | 5 | 59593138 | 0.509 | 0.522 | 0.488 | 0.506 | 0.643 | 0.553 | 0.856 | 0.775 | 0.852 | G | A |
| FILIP1;LOC101928540 | rs2951921 | 6 | 76165524 | 0.073 | NA | NA | 0.073 | 0.022 | 0.063 | 0.015 | 0.030 | 0.293 | T | C |
| SIRT2 | rs10405150 | 19 | 39387919 | 0.056 | NA | NA | 0.056 | 0.147 | 0.144 | 0.081 | 0.117 | 0.774 | C | T |
| LUCAT1 | rs10072084 | 5 | 90539203 | 0.541 | NA | NA | 0.541 | 0.559 | 0.422 | 0.232 | 0.429 | 0.822 | C | T |
| ZNF283 | rs16976816 | 19 | 44339377 | 0.977 | NA | NA | 0.977 | 0.970 | 0.994 | 0.987 | 0.976 | 0.864 | G | A |

| Label | SNP | CHR | POS | known | traitW | GeneUp | DistanceUp | GeneDown | DistanceDown |
|---|---|---|---|---|---|---|---|---|---|
| APOE | rs1160985 | 19 | 45403412 | APOE | C H L T | TOMM40 | 0 | TOMM40 | 0 |
| APOA1 | rs964184 | 11 | 116648917 | APOA1 | C H L T | ZPR1 | 0 | ZPR1 | 0 |
| APOB | rs754523 | 2 | 21311691 | APOB | C H L T | APOB | 44746 | TDRD15 | 35166 |
| ZHX2 | rs7841763 | 8 | 123971081 | NA | NA | ZHX2 | 0 | ZHX2 | 0 |
| PDE4D | rs7711093 | 5 | 59593138 | NA | NA | PDE4D | 0 | PDE4D | 0 |
| FILIP1;LOC101928540 | rs2951921 | 6 | 76165524 | NA | NA | FILIP1;LOC101928540 | 0 | FILIP1;LOC101928540 | 0 |
| SIRT2 | rs10405150 | 19 | 39387919 | NA | NA | SIRT2 | 0 | SIRT2 | 0 |
| LUCAT1 | rs10072084 | 5 | 90539203 | NA | NA | ADGRV1 | 79170 | LUCAT1 | 59599 |
| ZNF283 | rs16976816 | 19 | 44339377 | NA | NA | ZNF283 | 0 | ZNF283 | 0 |

| Label | SNP | CHR | POS | $P_D$ | $P_R$ | $P_{DR}$ | gwasPeak | gwasP |
|---|---|---|---|---|---|---|---|---|
| APOE | rs1160985 | 19 | 45403412 | **2.13E-13** | **3.36E-09** | **4.29E-21** | rs4420638 | **2.67E-16** |
| APOA1 | rs964184 | 11 | 116648917 | 5.37E-05 | 0.009 | *1.72E-06* | rs3741298 | *1.63E-07* |
| APOB | rs754523 | 2 | 21311691 | *6.25E-06* | 0.178 | 1.20E-05 | NA | NA |
| ZHX2 | rs7841763 | 8 | 123971081 | *4.82E-06* | 0.631 | 1.03E-04 | NA | NA |
| PDE4D | rs7711093 | 5 | 59593138 | *3.01E-06* | 0.747 | 5.37E-04 | NA | NA |
| FILIP1;LOC101928540 | rs2951921 | 6 | 76165524 | *9.04E-07* | NA | NA | NA | NA |
| SIRT2 | rs10405150 | 19 | 39387919 | *6.34E-06* | NA | NA | NA | NA |
| LUCAT1 | rs10072084 | 5 | 90539203 | *9.48E-06* | NA | NA | NA | NA |
| ZNF283 | rs16976816 | 19 | 44339377 | *9.78E-06* | NA | NA | NA | NA |

Table S2. Descriptions of the columns in the GWAS results tables.

| Column | Description |
|---|---|
| Label | Label for the locus derived from the list of known loci as labeled by Teslovich et al. and Willer et al., or if novel, the nearest gene. |
| SNP | SNP RS ID |
| CHR | chromosome |
| POS | physical position (hg19) in base pairs |
| $P_D$ | p-value in the GWAS Samoans from the 2010s (the discovery cohort) |
| $P_U$ | p-value in the unrelated Samoans from 1990s (the 1990-95 replication cohort) |
| $P_A$ | p-value in the adult Samoans from the 2000s (the 2002-03 replication cohort) |
| $P_R$ | meta-analysis p-value of the 1990s and 2000s samples |
| $P_{DR}$ | meta-analysis p-value of the 1990s, 2000s and 2010s samples. |
| $z_D$ | z-score, computed from the p-value, in the GWAS (all z-scores are for the transformed trait) |
| $z_U$ | z-score, computed from the p-value, in the unrelated Samoans from the 1990s |

| | |
|---|---|
| $z_A$ | z-score, computed from the p-value, in the adult Samoans from the 2000s |
| Dir | direction of the effect in each of the four samples (+ indicates the effect allele is increasing the trait value on the raw scale) |
| $n_D$ | sample size in the GWAS Samoans from the 2010s |
| $n_U$ | sample size in the unrelated Samoans from 1990s |
| $n_A$ | sample size in the adult Samoans from the 2000s |
| $n_R$ | meta-analysis sample size of the 1990s and 2000s samples |
| $n_{DR}$ | meta-analysis sample size of the 1990s, 2000s and 2010s samples. |
| eafD | effect allele frequency in the GWAS Samoans from the 2010s |
| eafU | effect allele frequency in the unrelated Samoans from 1990s |
| eafA | effect allele frequency in the adult Samoans from the 2000s |
| SAM | effect allele frequency in all Samoans |
| EAS | effect allele frequency in individuals of East Asian descent from the 1000 Genomes Project |
| SAS | effect allele frequency in individuals of South Asian descent from the 1000 Genomes Project |
| EUR | effect allele frequency in individuals of European descent from the 1000 Genomes Project |
| AMR | effect allele frequency in individuals of admixed American descent from the 1000 Genomes Project |
| AFR | effect allele frequency in individuals of African descent from the 1000 Genomes Project |
| eA | effect allele (the effect allele is the one associated with a poorer health outcome) |
| oA | other allele |
| known | known lipid-levels locus from Teslovich et al. 2010 and Willer et al. 2013 |
| traitW | traits associated with the locus listed in the 'known' column (C = Cholesterol, H = HDL, L = LDL, T = Triglycerides) |
| GeneUp | nearest gene upstream of the SNP |
| DistanceUp | distance in base pairs to nearest upstream gene (if 0 the SNP is within the transcription boundaries of the gene(s)) |
| GeneDown | nearest gene downstream of the SNP |
| DistanceDown | distance in base pairs to nearest downstream gene (if 0 the SNP is within the transcription boundaries of the gene(s)) |
| gwasPeak | sometimes the most significant SNP at a particular locus was not examined using OpenArray, if another SNP at that same locus was more strongly associated with another trait. E.g., rs4420638 was the peak SNP associated with total cholesterol at the APOE locus. However, rs1160985, a SNP also at the APOE locus was associated with LDL with a smaller p value. We selected a single SNP at each locus to replicate across all the traits and so we replicated rs1160985 for total cholesterol. This column (gwasPeak) contains the SNP that was the peak SNP at this locus for this trait if it is other than the SNP that was replicated. |
| gwasP | GWAS p value for gwasPeak |

Table S3. MAGENTA pathway analysis of TC. See Table S4 for a description of the column contents.

| Database | Gene Set | Effective Gene Set Size | FDR (95% cutoff) | Exp. # Genes (>95% cutoff) | Obs. # Genes (>95% cutoff) | Trait-specific Known Genes |
|---|---|---|---|---|---|---|
| GOTERM | high-density lipoprotein particle remodeling | 10 | 0.0186 | 1 | 5 | APOA1, APOE, CETP, LIPC, LIPG |
| GOTERM | reverse cholesterol transport | 13 | 0.03866667 | 1 | 5 | ABCA1, APOA1, APOE, CETP, LIPC, HNF1A, LIPG |

Table S4. Descriptions of the columns in the MAGENTA results tables.

| Column | Description |
|---|---|
| Database | Pathway database |
| Gene Set | Gene set name |
| Effective Gene Set Size | Effective number of genes per gene set analyzed by GSEA, after removing genes that were not assigned a gene score (e.g. no SNPs in their region), or after adjusting for physcial clustering of genes in a given gene set (removing all but one gene from a subset of genes assigned the same best SNP, keeping the gene with the most significant gene score) |
| FDR (95% cutoff) | Estimated false discovery rate (q-value) using the 95 percentile enrichment cutoff |
| Exp. # Genes (>95% cutoff) | Expected number of genes with a corrected gene p-value above the 95 percentile enrichment cutoff |
| Obs. # Genes (>95% cutoff) | Observed number of genes with a corrected gene p-value above the 95 percentile enrichment cutoff |
| Trait-specific Known Genes | Genes in the trait-specific list of known genes from Willer et al (2013) that belong to each gene set |

Table S5. INRICH analysis of TC. See Table S6 for a description of the INRICH parameters used.

| Gene Set Size | Number overlapping | Empirical P value | Corrected P value | Gene Set | Known overlapping genes |
|---|---|---|---|---|---|
| 68 | 4 | 0.002931 | 0.00379981 | Known TC genes | APOB, TRIB1, APOA1, APOE |
| 67 | 6 | 3.5e-05 | 9.9995e-05 | Known HDL genes | APOB, TRIB1, APOA1, SCARB1, LCAT, APOE |
| 52 | 4 | 0.000792999 | 0.00164992 | Known LDL genes | APOB, TRIB1, APOA1, APOE |
| 41 | 4 | 0.000705999 | 0.00119994 | Known TG genes | APOB, TRIB1, APOA1, APOE |

Table S6. INRICH parameter settings.

| Parameter | Value | Comment |
|---|---|---|
| gene-list (-g) | entrez_gene.hg19.map | Entrez genes |
| background-genes (-b) | --no background set-- | |
| range-file (-x) | --no ranges-- | |
| compact (-k) | YES | |
| target size-filter (-i,j) | 2..200 | |
| min-obs threshold (-z) | 2 | |
| test-type (-1) | INTERVALS | |
| top-N-regions (-n) | --all-- | |
| kb-window (-w) | 50000 | Extend target gene regions 50 kb up/downstream |
| match-density (-d) | 0.1 | Allow mapping SNP density of 90-110% |
| pre-compute (-c) | YES | |
| match-genes (-e) | YES | |
| num-replicates (-r) | 1000000 | |
| num-bootstraps (-q) | 20000 | |
| display-p (-p) | 1 | |

### 3. HDL

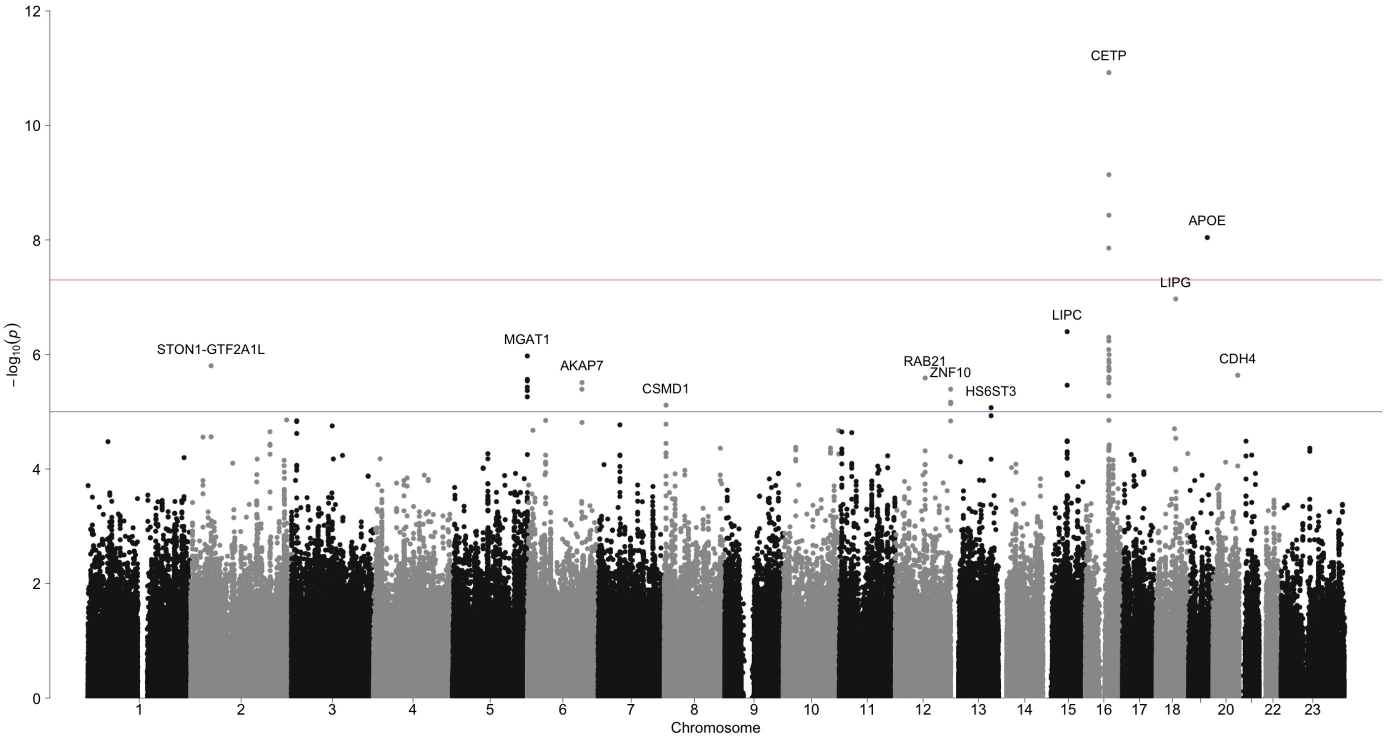


FIGURE S3. HDL: genome-wide association scan

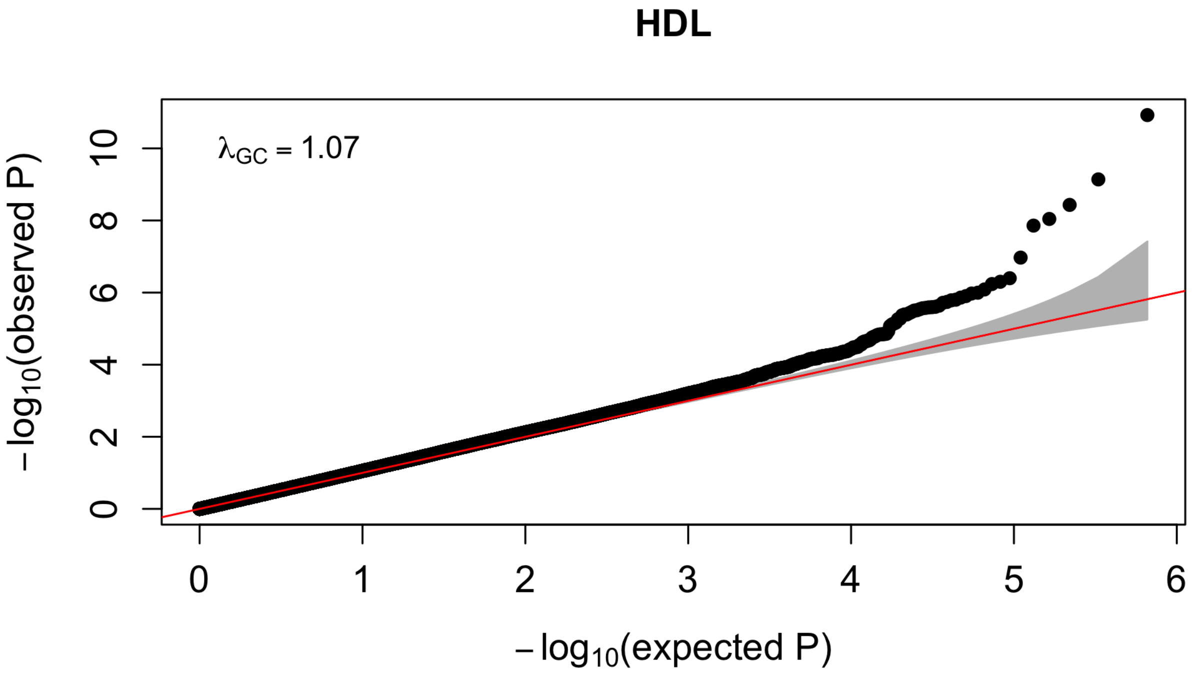


FIGURE S4. HDL: QQ plot

Table S7. HDL results. See Table S2 for a description of the column contents. Significant P values less than 5 x $10^{-8}$ are in **bold**, while suggestive P values between 1 x $10^{-5}$ and 5 x $10^{-8}$ are in *dark red italics*.

| Label | SNP | CHR | POS | $P_D$ | $P_U$ | $P_A$ | $P_R$ | $P_{DR}$ |
|---|---|---|---|---|---|---|---|---|
| MGAT1 | rs1038143 | 5 | 180213878 | *3.72E-06* | 0.010 | 0.301 | 0.016 | *2.91E-07* |
| RAB21 | rs328733 | 12 | 72197574 | *2.57E-06* | 0.011 | 0.517 | 0.036 | *5.92E-07* |
| LIPC | rs10438284 | 15 | 58629424 | *4.00E-07* | 0.905 | 0.043 | 0.133 | *9.15E-07* |
| CDH4 | rs817687 | 20 | 59753355 | *2.31E-06* | 0.206 | 0.615 | 0.237 | *8.77E-06* |
| AKAP7 | rs3777486 | 6 | 131584648 | *3.09E-06* | 0.050 | 0.212 | 0.808 | 4.37E-04 |
| APOE | rs1160985 | 19 | 45403412 | 0.003 | 0.722 | 0.133 | 0.342 | 0.004 |
| CETP | rs289708 | 16 | 57038162 | **1.19E-11** | NA | NA | NA | NA |
| LIPG | rs16950739 | 18 | 47138509 | *1.07E-07* | NA | NA | NA | NA |
| STON1-GTF2A1L | rs6739536 | 2 | 48831901 | *1.58E-06* | NA | NA | NA | NA |
| ZNF10 | rs2292029 | 12 | 133734113 | *4.05E-06* | NA | NA | NA | NA |
| CSMD1 | rs1626142 | 8 | 4345284 | *7.67E-06* | NA | NA | NA | NA |
| HS6ST3 | rs16953620 | 13 | 97508453 | *8.48E-06* | NA | NA | NA | NA |

| Label | SNP | CHR | POS | $z_D$ | $z_U$ | $z_A$ | Dir | $n_D$ | $n_U$ | $n_A$ | $n_R$ | $n_{DR}$ |
|---|---|---|---|---|---|---|---|---|---|---|---|---|
| MGAT1 | rs1038143 | 5 | 180213878 | -4.63 | -2.57 | -1.04 | --- | 2849 | 695 | 1071 | 1766 | 4615 |
| RAB21 | rs328733 | 12 | 72197574 | -4.70 | -2.55 | -0.65 | --- | 2849 | 697 | 1076 | 1773 | 4622 |
| LIPC | rs10438284 | 15 | 58629424 | -5.07 | 0.12 | -2.02 | -+- | 2847 | 697 | 1077 | 1774 | 4621 |
| CDH4 | rs817687 | 20 | 59753355 | -4.72 | -1.27 | -0.50 | --- | 2840 | 678 | 1074 | 1752 | 4592 |
| AKAP7 | rs3777486 | 6 | 131584648 | -4.67 | 1.96 | -1.25 | -+- | 2847 | 682 | 1077 | 1759 | 4606 |
| APOE | rs1160985 | 19 | 45403412 | -2.96 | 0.36 | -1.50 | -+- | 2849 | 692 | 1075 | 1767 | 4616 |
| CETP | rs289708 | 16 | 57038162 | -6.78 | NA | NA | -?? | 2849 | NA | NA | NA | NA |
| LIPG | rs16950739 | 18 | 47138509 | -5.31 | NA | NA | -?? | 2849 | NA | NA | NA | NA |
| STON1-GTF2A1L | rs6739536 | 2 | 48831901 | -4.80 | NA | NA | -?? | 2847 | NA | NA | NA | NA |
| ZNF10 | rs2292029 | 12 | 133734113 | -4.61 | NA | NA | -?? | 2848 | NA | NA | NA | NA |
| CSMD1 | rs1626142 | 8 | 4345284 | -4.47 | NA | NA | -?? | 2756 | NA | NA | NA | NA |
| HS6ST3 | rs16953620 | 13 | 97508453 | -4.45 | NA | NA | -?? | 2847 | NA | NA | NA | NA |

| Label | SNP | CHR | POS | eafD | eafU | eafA | SAM | EAS | SAS | EUR | AMR | AFR | eA | oA |
|---|---|---|---|---|---|---|---|---|---|---|---|---|---|---|
| MGAT1 | rs1038143 | 5 | 180213878 | 0.310 | 0.303 | 0.311 | 0.309 | 0.110 | 0.137 | 0.148 | 0.081 | 0.013 | A | G |
| RAB21 | rs328733 | 12 | 72197574 | 0.788 | 0.799 | 0.782 | 0.788 | 0.626 | 0.700 | 0.869 | 0.738 | 0.611 | T | C |
| LIPC | rs10438284 | 15 | 58629424 | 0.290 | 0.289 | 0.276 | 0.286 | 0.326 | 0.173 | 0.272 | 0.199 | 0.047 | A | G |
| CDH4 | rs817687 | 20 | 59753355 | 0.985 | 0.987 | 0.989 | 0.986 | 0.924 | 0.945 | 0.967 | 0.855 | 0.610 | C | T |
| AKAP7 | rs3777486 | 6 | 131584648 | 0.975 | 0.978 | 0.976 | 0.976 | 0.909 | 0.898 | 0.858 | 0.793 | 0.949 | T | C |
| APOE | rs1160985 | 19 | 45403412 | 0.732 | 0.712 | 0.709 | 0.724 | 0.659 | 0.590 | 0.554 | 0.447 | 0.378 | C | T |
| CETP | rs289708 | 16 | 57038162 | 0.905 | NA | NA | 0.905 | 0.815 | 0.800 | 0.860 | 0.823 | 0.597 | G | A |
| LIPG | rs16950739 | 18 | 47138509 | 0.019 | NA | NA | 0.019 | 0.058 | 0.190 | 0.057 | 0.140 | 0.004 | T | C |
| STON1-GTF2A1L | rs6739536 | 2 | 48831901 | 0.762 | NA | NA | 0.762 | 0.676 | 0.837 | 0.918 | 0.842 | 0.609 | T | A |
| ZNF10 | rs2292029 | 12 | 133734113 | 0.179 | NA | NA | 0.179 | 0.112 | 0.254 | 0.245 | 0.156 | 0.008 | T | C |
| CSMD1 | rs1626142 | 8 | 4345284 | 0.612 | NA | NA | 0.612 | 0.400 | 0.435 | 0.298 | 0.392 | 0.654 | G | A |
| HS6ST3 | rs16953620 | 13 | 97508453 | 0.968 | NA | NA | 0.968 | 0.950 | 0.949 | 0.848 | 0.911 | 0.836 | A | G |

| Label | SNP | CHR | POS | known | traitW | GeneUp | DistanceUp | GeneDown | DistanceDown |
|---|---|---|---|---|---|---|---|---|---|
| MGAT1 | rs1038143 | 5 | 180213878 | NA | NA | OR2Y1 | 46820 | MGAT1 | 3662 |
| RAB21 | rs328733 | 12 | 72197574 | NA | NA | RAB21 | 10318 | TBC1D15 | 35912 |
| LIPC | rs10438284 | 15 | 58629424 | LIPC | C H T | AQP9 | 151314 | LIPC | 94750 |
| CDH4 | rs817687 | 20 | 59753355 | NA | NA | LINC01718 | 98120 | CDH4 | 74126 |
| AKAP7 | rs3777486 | 6 | 131584648 | NA | NA | AKAP7 | 0 | AKAP7 | 0 |
| APOE | rs1160985 | 19 | 45403412 | APOE | C H L T | TOMM40 | 0 | TOMM40 | 0 |
| CETP | rs289708 | 16 | 57038162 | CETP | C H L T | NLRC5 | 0 | NLRC5 | 0 |
| LIPG | rs16950739 | 18 | 47138509 | LIPG | C H | LIPG | 19230 | ACAA2 | 171364 |
| STON1-GTF2A1L | rs6739536 | 2 | 48831901 | NA | NA | STON1-GTF2A1L | 0 | STON1-GTF2A1L | 0 |
| ZNF10 | rs2292029 | 12 | 133734113 | NA | NA | ZNF10 | 0 | ZNF10 | 0 |
| CSMD1 | rs1626142 | 8 | 4345284 | NA | NA | CSMD1 | 0 | CSMD1 | 0 |
| HS6ST3 | rs16953620 | 13 | 97508453 | NA | NA | HS6ST3 | 16637 | LINC00359 | 85081 |

| Label | SNP | CHR | POS | $P_D$ | $P_R$ | $P_{DR}$ | gwasPeak | gwasP |
|---|---|---|---|---|---|---|---|---|
| MGAT1 | rs1038143 | 5 | 180213878 | *3.72E-06* | 0.016 | *2.91E-07* | rs249356 | *1.06E-06* |
| RAB21 | rs328733 | 12 | 72197574 | *2.57E-06* | 0.036 | *5.92E-07* | NA | NA |
| LIPC | rs10438284 | 15 | 58629424 | *4.00E-07* | 0.133 | *9.15E-07* | NA | NA |
| CDH4 | rs817687 | 20 | 59753355 | *2.31E-06* | 0.237 | *8.77E-06* | NA | NA |
| AKAP7 | rs3777486 | 6 | 131584648 | *3.09E-06* | 0.808 | 4.37E-04 | NA | NA |
| APOE | rs1160985 | 19 | 45403412 | 0.003 | 0.342 | 0.004 | rs4420638 | **9.07E-09** |
| CETP | rs289708 | 16 | 57038162 | **1.19E-11** | NA | NA | NA | NA |
| LIPG | rs16950739 | 18 | 47138509 | *1.07E-07* | NA | NA | NA | NA |
| STON1-GTF2A1L | rs6739536 | 2 | 48831901 | *1.58E-06* | NA | NA | NA | NA |
| ZNF10 | rs2292029 | 12 | 133734113 | *4.05E-06* | NA | NA | NA | NA |
| CSMD1 | rs1626142 | 8 | 4345284 | *7.67E-06* | NA | NA | NA | NA |
| HS6ST3 | rs16953620 | 13 | 97508453 | *8.48E-06* | NA | NA | NA | NA |

Table S8. MAGENTA pathway analysis of HDL. See Table S4 for a description of the column contents.

| Database | Gene Set | Effective Gene Set Size | FDR (95% cutoff) | Exp. # Genes (>95% cutoff) | Obs. # Genes (>95% cutoff) | Trait-specific Known Genes |
|---|---|---|---|---|---|---|
| GOTERM | high-density lipoprotein particle remodeling | 10 | 0 | 1 | 8 | APOA1, APOE, SCARB1, CETP, LCAT, LIPC, LIPG |
| GOTERM | reverse cholesterol transport | 13 | 0 | 1 | 9 | ABCA1, APOA1, APOE, SCARB1, CETP, LCAT, LIPC, LIPG |
| GOTERM | high-density lipoprotein particle | 15 | 3.333333e-05 | 1 | 8 | APOA1, APOE, CETP, LCAT, LIPC |
| REACTOME | LIPOPROTEIN METABOLISM | 24 | 8e-04 | 1 | 9 | ABCA1, APOA1, APOB, APOE, SCARB1, CETP, LCAT, LIPC, LPL |
| REACTOME | HDL MEDIATED LIPID TRANSPORT | 11 | 0.00315 | 1 | 5 | ABCA1, APOA1, SCARB1, CETP, LCAT |
| GOTERM | cholesterol homeostasis | 36 | 0.00508 | 2 | 10 | ABCA1, APOA1, APOB, APOE, SCARB1, CETP, LCAT, LIPC, LIPG |
| GOTERM | phospholipid efflux | 7 | 0.005225 | 0 | 4 | ABCA1, APOA1, APOE |

| | | | | | | |
|---|---|---|---|---|---|---|
| GOTERM | cholesterol efflux | 16 | 0.006466667 | 1 | 6 | ABCA1, APOA1, APOE, SCARB1 |
| GOTERM | cholesterol transporter activity | 12 | 0.007114286 | 1 | 5 | ABCA1, APOA1, APOB, APOE, CETP, STARD3 |
| GOTERM | cholesterol transport | 12 | 0.0090625 | 1 | 5 | APOA1, APOB, CETP, LCAT |
| GOTERM | cholesterol binding | 18 | 0.009444444 | 1 | 6 | ABCA1, APOA1, CETP, STARD3 |
| Ingenuity | LXR.RXR.Activation | 34 | 0.0129 | 2 | 8 | ABCA1, APOA1, APOE, CETP, LCAT, LPL, PLTP |
| PANTHER BIOLOGICAL PROCESS | Other apoptosis | 10 | 0.0146 | 1 | 4 | SCARB1 |
| REACTOME | CHYLOMICRON MEDIATED LIPID TRANSPORT | 14 | 0.01463333 | 1 | 5 | APOA1, APOB, APOE, LIPC, LPL |
| GOTERM | cholesterol catabolic process | 10 | 0.0215 | 1 | 4 | APOE, SCARB1 |
| PANTHER MOLECULAR FUNCTION | Apolipoprotein | 15 | 0.025 | 1 | 5 | APOA1, APOB, APOE |

Table S9. INRICH analysis of HDL. See Table S6 for a description of the INRICH parameters used.

| Gene Set Size | Number overlapping | Empirical P value | Corrected P value | Gene Set | Known overlapping genes |
|---|---|---|---|---|---|
| 68 | 7 | 7.99999e-06 | 4.99975e-05 | Known TC genes | RAF1, GPAM, APOA1, LIPC, CETP, LIPG, APOE |
| 67 | 7 | 3.5e-05 | 9.9995e-05 | Known HDL genes | VEGFA, APOA1, LIPC, CETP, LCAT, LIPG, APOE |
| 52 | 4 | 0.002151 | 0.00654967 | Known LDL genes | GPAM, APOA1, CETP, APOE |
| 41 | 5 | 9.59999e-05 | 0.000349983 | Known TG genes | VEGFA, APOA1, LIPC, CETP, APOE |

### 4. LDL

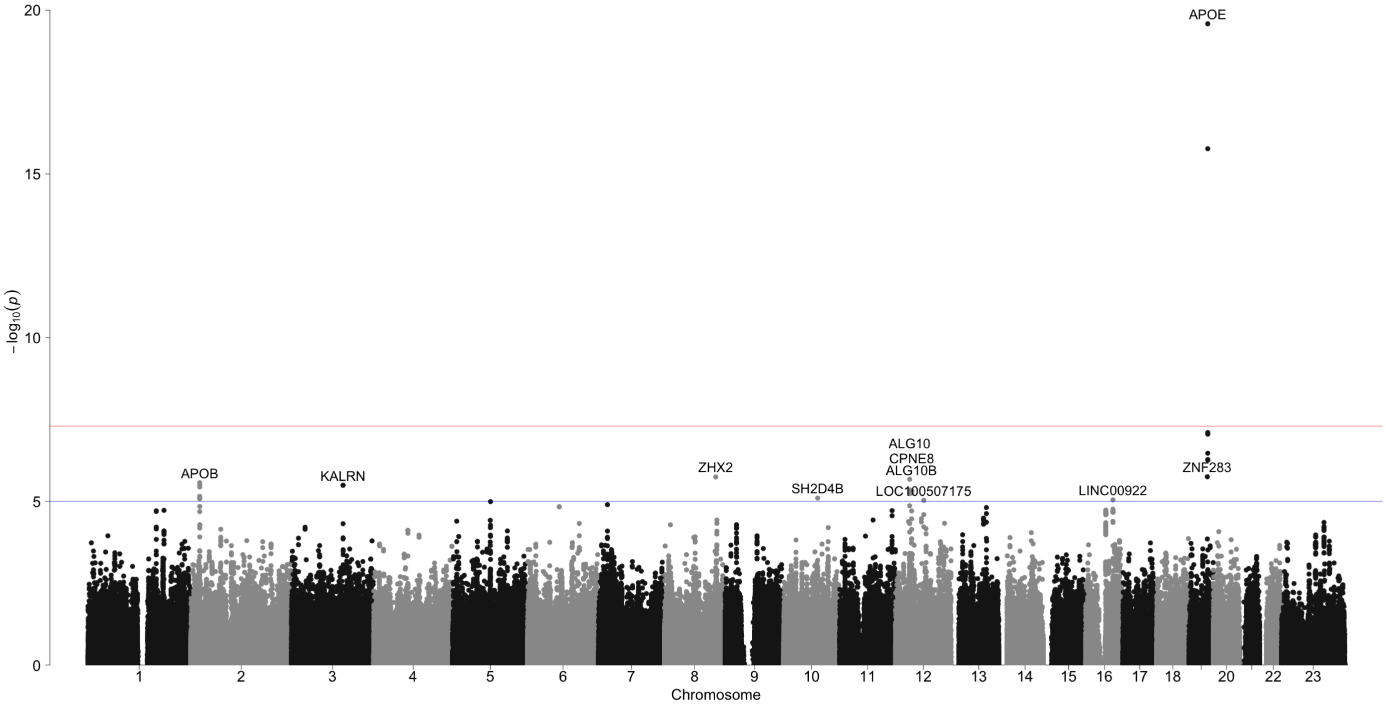


FIGURE S5. LDL: genome-wide association scan

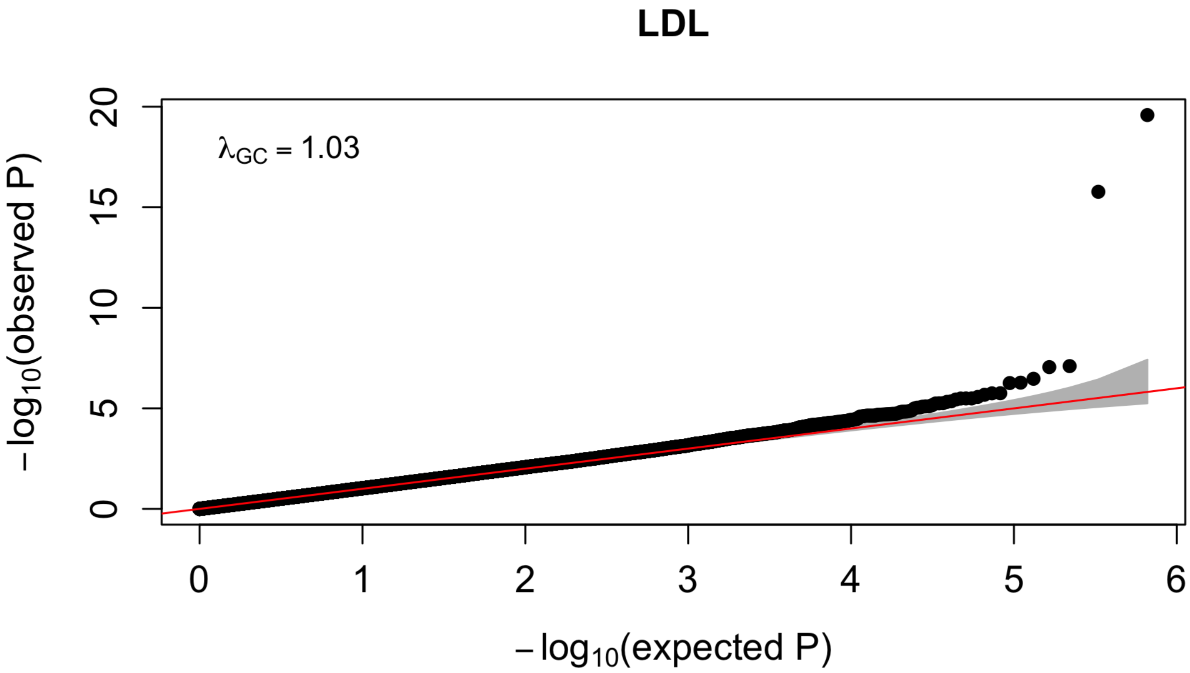


FIGURE S6. LDL: QQ plot

Table S10. LDL results. See Table S2 for a description of the column contents. Significant P values less than 5 x $10^{-8}$ are in **bold**, while suggestive P values between 1 x $10^{-5}$ and 5 x $10^{-8}$ are in *dark red italics*.

| Label | SNP | CHR | POS | $P_D$ | $P_U$ | $P_A$ | $P_R$ | $P_{DR}$ |
|---|---|---|---|---|---|---|---|---|
| APOE | rs1160985 | 19 | 45403412 | **2.61E-20** | 0.001 | *7.88E-07* | **5.07E-09** | **1.53E-27** |
| APOB | rs754523 | 2 | 21311691 | *3.25E-06* | 0.247 | 0.381 | 0.158 | *5.81E-06* |
| ZHX2 | rs7841763 | 8 | 123971081 | *1.80E-06* | 0.293 | 0.920 | 0.557 | 3.78E-05 |
| ALG10 | rs3912355 | 12 | 34079616 | *2.12E-06* | 0.514 | 0.801 | 0.544 | 3.97E-05 |
| CPNE8 | rs11169807 | 12 | 39244161 | *4.77E-06* | 0.464 | 0.654 | 0.418 | 4.13E-05 |
| KALRN | rs6789134 | 3 | 123942339 | *3.22E-06* | 0.497 | 0.667 | 0.925 | 1.96E-04 |
| ZNF283 | rs16976816 | 19 | 44339377 | *1.78E-06* | NA | NA | NA | NA |
| ALG10B | rs10880642 | 12 | 38554152 | *5.56E-06* | NA | NA | NA | NA |
| SH2D4B | rs10509415 | 10 | 82473065 | *7.96E-06* | NA | NA | NA | NA |
| LINC00922 | rs254371 | 16 | 65943650 | *9.04E-06* | NA | NA | NA | NA |
| LINC02408 | rs17104016 | 12 | 67969929 | *9.29E-06* | NA | NA | NA | NA |

| Label | SNP | CHR | POS | $z_D$ | $z_U$ | $z_A$ | Dir | $n_D$ | $n_U$ | $n_A$ | $n_R$ | $n_{DR}$ |
|---|---|---|---|---|---|---|---|---|---|---|---|---|
| APOE | rs1160985 | 19 | 45403412 | 9.23 | 3.19 | 4.94 | +++ | 2842 | 692 | 1047 | 1739 | 4581 |
| APOB | rs754523 | 2 | 21311691 | 4.65 | 1.16 | 0.88 | +++ | 2841 | 697 | 1048 | 1745 | 4586 |
| ZHX2 | rs7841763 | 8 | 123971081 | 4.78 | 1.05 | -0.10 | ++- | 2842 | 696 | 1048 | 1744 | 4586 |
| ALG10 | rs3912355 | 12 | 34079616 | 4.74 | 0.65 | 0.25 | +++ | 2833 | 688 | 1046 | 1734 | 4567 |
| CPNE8 | rs11169807 | 12 | 39244161 | 4.58 | 0.73 | 0.45 | +++ | 2842 | 695 | 1050 | 1745 | 4587 |
| KALRN | rs6789134 | 3 | 123942339 | 4.66 | 0.68 | -0.43 | ++- | 2842 | 692 | 1049 | 1741 | 4583 |
| ZNF283 | rs16976816 | 19 | 44339377 | 4.78 | NA | NA | +?? | 2842 | NA | NA | NA | NA |
| ALG10B | rs10880642 | 12 | 38554152 | 4.54 | NA | NA | +?? | 2830 | NA | NA | NA | NA |
| SH2D4B | rs10509415 | 10 | 82473065 | 4.47 | NA | NA | +?? | 2842 | NA | NA | NA | NA |
| LINC00922 | rs254371 | 16 | 65943650 | 4.44 | NA | NA | +?? | 2842 | NA | NA | NA | NA |
| LINC02408 | rs17104016 | 12 | 67969929 | 4.43 | NA | NA | +?? | 2806 | NA | NA | NA | NA |

| Label | SNP | CHR | POS | eafD | eafU | eafA | SAM | EAS | SAS | EUR | AMR | AFR | eA | oA |
|---|---|---|---|---|---|---|---|---|---|---|---|---|---|---|
| APOE | rs1160985 | 19 | 45403412 | 0.732 | 0.712 | 0.709 | 0.724 | 0.659 | 0.590 | 0.554 | 0.447 | 0.378 | C | T |
| APOB | rs754523 | 2 | 21311691 | 0.254 | 0.247 | 0.229 | 0.247 | 0.265 | 0.140 | 0.309 | 0.306 | 0.201 | G | A |
| ZHX2 | rs7841763 | 8 | 123971081 | 0.040 | 0.047 | 0.049 | 0.043 | 0.023 | 0.146 | 0.102 | 0.058 | 0.169 | T | C |
| ALG10 | rs3912355 | 12 | 34079616 | 0.855 | 0.856 | 0.857 | 0.855 | 0.872 | 0.651 | 0.605 | 0.732 | 0.863 | C | T |
| CPNE8 | rs11169807 | 12 | 39244161 | 0.802 | 0.760 | 0.794 | 0.794 | 0.644 | 0.536 | 0.505 | 0.408 | 0.898 | C | T |
| KALRN | rs6789134 | 3 | 123942339 | 0.079 | 0.074 | 0.076 | 0.078 | 0.161 | 0.120 | 0.046 | 0.118 | 0.212 | G | A |
| ZNF283 | rs16976816 | 19 | 44339377 | 0.977 | NA | NA | 0.977 | 0.970 | 0.994 | 0.987 | 0.976 | 0.864 | G | A |
| ALG10B | rs10880642 | 12 | 38554152 | 0.802 | NA | NA | 0.802 | 0.715 | 0.555 | 0.481 | 0.532 | 0.299 | A | G |
| SH2D4B | rs10509415 | 10 | 82473065 | 0.706 | NA | NA | 0.706 | 0.510 | 0.735 | 0.758 | 0.808 | 0.708 | A | C |
| LINC00922 | rs254371 | 16 | 65943650 | 0.555 | NA | NA | 0.555 | 0.698 | 0.650 | 0.599 | 0.693 | 0.884 | T | C |
| LINC02408 | rs17104016 | 12 | 67969929 | 0.574 | NA | NA | 0.574 | 0.734 | 0.924 | 0.853 | 0.906 | 0.838 | A | T |

| Label | SNP | CHR | POS | known | traitW | GeneUp | DistanceUp | GeneDown | DistanceDown |
|---|---|---|---|---|---|---|---|---|---|
| APOE | rs1160985 | 19 | 45403412 | APOE | C H L T | TOMM40 | 0 | TOMM40 | 0 |
| APOB | rs754523 | 2 | 21311691 | APOB | C H L T | APOB | 44746 | TDRD15 | 35166 |
| ZHX2 | rs7841763 | 8 | 123971081 | NA | NA | ZHX2 | 0 | ZHX2 | 0 |
| ALG10 | rs3912355 | 12 | 34079616 | NA | NA | SYT10 | 486862 | ALG10 | 95599 |
| CPNE8 | rs11169807 | 12 | 39244161 | NA | NA | CPNE8 | 0 | CPNE8 | 0 |
| KALRN | rs6789134 | 3 | 123942339 | NA | NA | KALRN | 0 | KALRN | 0 |
| ZNF283 | rs16976816 | 19 | 44339377 | NA | NA | ZNF283 | 0 | ZNF283 | 0 |
| ALG10B | rs10880642 | 12 | 38554152 | NA | NA | ALG10 | 4372916 | ALG10B | 156404 |
| SH2D4B | rs10509415 | 10 | 82473065 | NA | NA | SH2D4B | 66749 | NRG3 | 1162004 |
| LINC00922 | rs254371 | 16 | 65943650 | NA | NA | LINC00922 | 333447 | CDH5 | 456859 |
| LINC02408 | rs17104016 | 12 | 67969929 | NA | NA | LINC02408 | 9018 | DYRK2 | 72582 |

| Label | SNP | CHR | POS | $P_D$ | $P_R$ | $P_{DR}$ | gwasPeak | gwasP |
|---|---|---|---|---|---|---|---|---|
| APOE | rs1160985 | 19 | 45403412 | **2.61E-20** | **5.07E-09** | **1.53E-27** | NA | NA |
| APOB | rs754523 | 2 | 21311691 | *3.25E-06* | 0.158 | *5.81E-06* | rs1469513 | *2.71E-06* |
| ZHX2 | rs7841763 | 8 | 123971081 | *1.80E-06* | 0.557 | 3.78E-05 | NA | NA |
| ALG10 | rs3912355 | 12 | 34079616 | *2.12E-06* | 0.544 | 3.97E-05 | NA | NA |
| CPNE8 | rs11169807 | 12 | 39244161 | *4.77E-06* | 0.418 | 4.13E-05 | NA | NA |
| KALRN | rs6789134 | 3 | 123942339 | *3.22E-06* | 0.925 | 1.96E-04 | NA | NA |
| ZNF283 | rs16976816 | 19 | 44339377 | *1.78E-06* | NA | NA | NA | NA |
| ALG10B | rs10880642 | 12 | 38554152 | *5.56E-06* | NA | NA | NA | NA |
| SH2D4B | rs10509415 | 10 | 82473065 | *7.96E-06* | NA | NA | NA | NA |
| LINC00922 | rs254371 | 16 | 65943650 | *9.04E-06* | NA | NA | NA | NA |
| LINC02408 | rs17104016 | 12 | 67969929 | *9.29E-06* | NA | NA | NA | NA |

Table S11. MAGENTA pathway analysis of LDL. See Table S4 for a description of the column contents.

| Database | Gene Set | Effective Gene Set Size | FDR (95% cutoff) | Exp. # Genes (>95% cutoff) | Obs. # Genes (>95% cutoff) | Trait-specific Known Genes |
|---|---|---|---|---|---|---|
| PANTHER MOLECULAR FUNCTION | Amylase | 4 | 0.19 | 0 | 2 | - |

Table S12. INRICH analysis of LDL. See Table S6 for a description of the INRICH parameters used.

| Gene Set Size | Number overlapping | Empirical P value | Corrected P value | Gene Set | Known overlapping genes |
|---|---|---|---|---|---|
| 68 | 5 | 8.29999e-05 | 0.000249988 | Known TC genes | APOB, CMTM6, DNAH11, TRIB1, APOE |
| 67 | 3 | 0.00803399 | 0.0148493 | Known HDL genes | APOB, TRIB1, APOE |
| 52 | 6 | 3e-06 | 4.99975e-05 | Known LDL genes | APOB, CMTM6, DNAH11, TRIB1, APOE, SPTLC3 |
| 41 | 3 | 0.00177 | 0.00439978 | Known TG genes | APOB, TRIB1, APOE |

### 5. Triglycerides (TG)

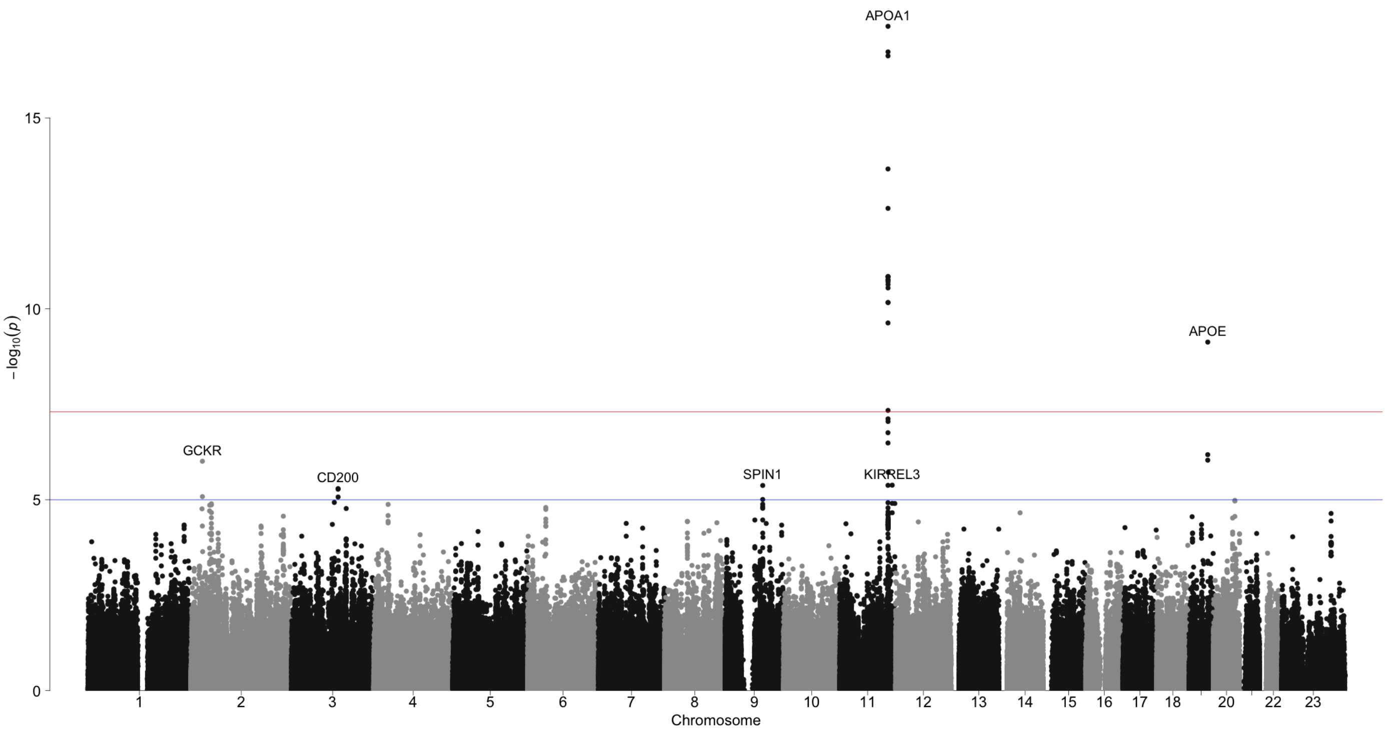


Figure S7. TG: genome-wide association scan

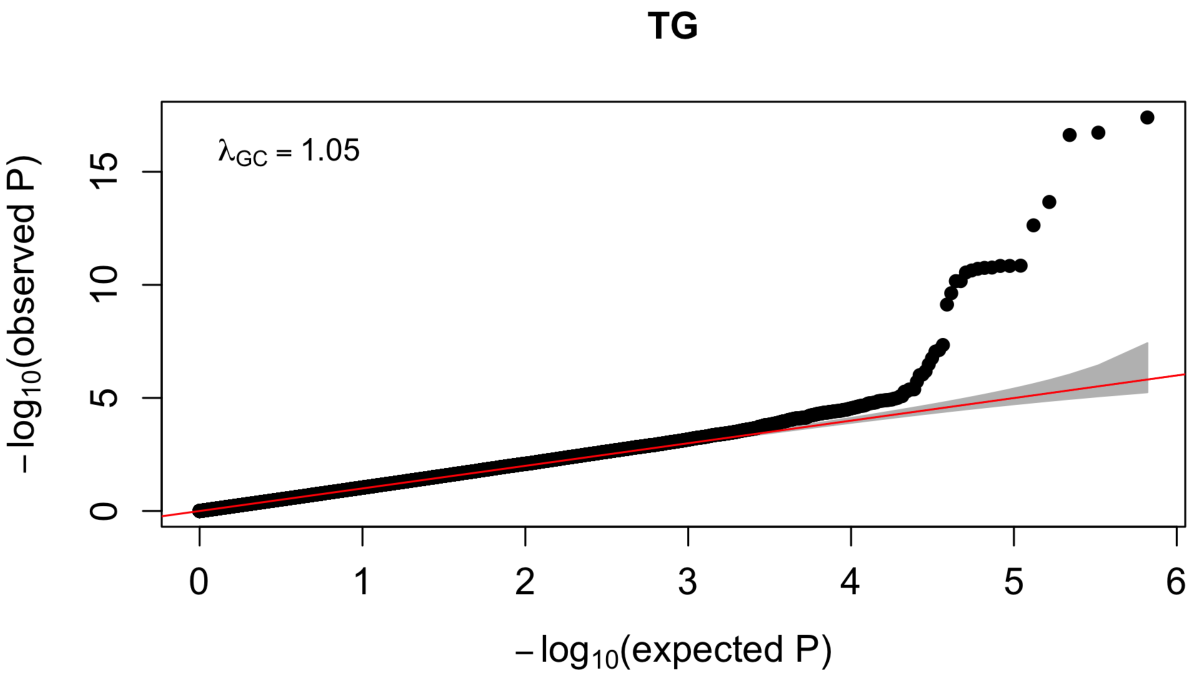


Figure S8. TG: QQ plot

TABLE S13. TG results. See Table S2 for a description of the column contents. Significant P values less than 5 x $10^{-8}$ are in **bold**, while suggestive P values between 1 x $10^{-5}$ and 5 x $10^{-8}$ are in *dark red italics*.

| Label | SNP | CHR | POS | $P_D$ | $P_U$ | $P_A$ | $P_R$ | $P_{DR}$ |
|---|---|---|---|---|---|---|---|---|
| APOA1 | rs964184 | 11 | 116648917 | **2.37E-17** | *7.27E-07* | **2.39E-08** | **8.97E-14** | **1.81E-29** |
| GCKR | rs780094 | 2 | 27741237 | *9.84E-07* | 0.016 | 0.175 | 0.010 | *5.62E-08* |
| CD200 | rs2399416 | 3 | 112059213 | *5.12E-06* | 0.635 | 0.353 | 0.668 | 9.29E-04 |
| APOE | rs1160985 | 19 | 45403412 | 0.312 | 0.672 | 0.940 | 0.836 | 0.507 |
| KIRREL3 | rs3018434 | 11 | 126805881 | *4.16E-06* | NA | NA | NA | NA |
| SPIN1 | rs7861888 | 9 | 90886340 | *4.24E-06* | NA | NA | NA | NA |

| Label | SNP | CHR | POS | $z_D$ | $z_U$ | $z_A$ | Dir | $n_D$ | $n_U$ | $n_A$ | $n_R$ | $n_{DR}$ |
|---|---|---|---|---|---|---|---|---|---|---|---|---|
| APOA1 | rs964184 | 11 | 116648917 | 8.47 | 4.95 | 5.58 | +++ | 2849 | 710 | 1080 | 1790 | 4639 |
| GCKR | rs780094 | 2 | 27741237 | 4.90 | 2.40 | 1.36 | +++ | 2847 | 710 | 1080 | 1790 | 4637 |
| CD200 | rs2399416 | 3 | 112059213 | 4.56 | 0.47 | -0.93 | ++- | 2832 | 692 | 1078 | 1770 | 4602 |
| APOE | rs1160985 | 19 | 45403412 | 1.01 | -0.42 | 0.08 | +-+ | 2849 | 704 | 1077 | 1781 | 4630 |
| KIRREL3 | rs3018434 | 11 | 126805881 | 4.60 | NA | NA | +?? | 2844 | NA | NA | NA | NA |
| SPIN1 | rs7861888 | 9 | 90886340 | 4.60 | NA | NA | +?? | 2816 | NA | NA | NA | NA |

| Label | SNP | CHR | POS | eafD | eafU | eafA | SAM | EAS | SAS | EUR | AMR | AFR | eA | oA |
|---|---|---|---|---|---|---|---|---|---|---|---|---|---|---|
| APOA1 | rs964184 | 11 | 116648917 | 0.430 | 0.453 | 0.457 | 0.440 | 0.240 | 0.229 | 0.162 | 0.277 | 0.221 | G | C |
| GCKR | rs780094 | 2 | 27741237 | 0.332 | 0.347 | 0.331 | 0.334 | 0.476 | 0.198 | 0.411 | 0.360 | 0.132 | T | C |
| CD200 | rs2399416 | 3 | 112059213 | 0.020 | 0.015 | 0.028 | 0.021 | 0.148 | 0.140 | 0.393 | 0.272 | 0.101 | A | G |
| APOE | rs1160985 | 19 | 45403412 | 0.268 | 0.288 | 0.291 | 0.276 | 0.341 | 0.410 | 0.446 | 0.553 | 0.623 | T | C |
| KIRREL3 | rs3018434 | 11 | 126805881 | 0.916 | NA | NA | 0.916 | 0.803 | 0.920 | 0.866 | 0.857 | 0.974 | G | A |
| SPIN1 | rs7861888 | 9 | 90886340 | 0.706 | NA | NA | 0.706 | 0.719 | 0.897 | 0.921 | 0.842 | 0.989 | A | G |

| Label | SNP | CHR | POS | known | traitW | GeneUp | DistanceUp | GeneDown | DistanceDown |
|---|---|---|---|---|---|---|---|---|---|
| APOA1 | rs964184 | 11 | 116648917 | APOA1 | C H L T | ZPR1 | 0 | ZPR1 | 0 |
| GCKR | rs780094 | 2 | 27741237 | GCKR | C T | GCKR | 0 | GCKR | 0 |
| CD200 | rs2399416 | 3 | 112059213 | NA | NA | CD200 | 0 | CD200 | 0 |
| APOE | rs1160985 | 19 | 45403412 | APOE | C H L T | TOMM40 | 0 | TOMM40 | 0 |
| KIRREL3 | rs3018434 | 11 | 126805881 | NA | NA | KIRREL3 | 0 | KIRREL3 | 0 |
| SPIN1 | rs7861888 | 9 | 90886340 | NA | NA | SPATA31C2 | 136440 | SPIN1 | 116956 |

| Label | SNP | CHR | POS | $P_D$ | $P_R$ | $P_{DR}$ | gwasPeak | gwasP |
|---|---|---|---|---|---|---|---|---|
| APOA1 | rs964184 | 11 | 116648917 | **2.37E-17** | **8.97E-14** | **1.81E-29** | rs6589566 | **3.98E-18** |
| GCKR | rs780094 | 2 | 27741237 | *9.84E-07* | 0.010 | *5.62E-08* | NA | NA |
| CD200 | rs2399416 | 3 | 112059213 | *5.12E-06* | 0.668 | 9.29E-04 | NA | NA |
| APOE | rs1160985 | 19 | 45403412 | 0.312 | 0.836 | 0.507 | rs4420638 | **7.44E-10** |
| KIRREL3 | rs3018434 | 11 | 126805881 | *4.16E-06* | NA | NA | NA | NA |
| SPIN1 | rs7861888 | 9 | 90886340 | *4.24E-06* | NA | NA | NA | NA |

Table S14. MAGENTA pathway analysis of TG. See Table S4 for a description of the column contents.

| Database | Gene Set | Effective Gene Set Size | FDR (95% cutoff) | Exp. # Genes (>95% cutoff) | Obs. # Genes (>95% cutoff) | Trait-specific Known Genes |
|---|---|---|---|---|---|---|
| GOTERM | cholesterol efflux | 15 | 0.0018 | 1 | 7 | APOA1, APOE |
| GOTERM | reverse cholesterol transport | 13 | 0.00405 | 1 | 6 | APOA1, APOE, CETP, LIPC |
| GOTERM | phospholipid efflux | 7 | 0.005866667 | 0 | 4 | APOA1, APOE |
| GOTERM | thyroid hormone receptor binding | 24 | 0.030375 | 1 | 7 | JMJD1C |
| GOTERM | ligand-dependent nuclear receptor transcription coactivator activity | 26 | 0.0389 | 1 | 7 | - |
| GOTERM | high-density lipoprotein particle remodeling | 10 | 0.04408 | 1 | 4 | APOA1, APOE, CETP, LIPC |
| GOTERM | fatty acid biosynthetic process | 48 | 0.04907143 | 2 | 10 | LIPC, LPL |

Table S15. INRICH analysis of TG. See Table S6 for a description of the INRICH parameters used.

| Gene Set Size | Number overlapping | Empirical P value | Corrected P value | Gene Set | Known overlapping genes |
|---|---|---|---|---|---|
| 68 | 5 | 0.000306 | 0.00039998 | Known TC genes | GCKR, SOX17, TRIB1, APOA1, APOE |
| 67 | 5 | 0.000446 | 0.00099995 | Known HDL genes | VEGFA, TRIB1, APOA1, SCARB1, APOE |
| 52 | 4 | 0.000987999 | 0.00214989 | Known LDL genes | SOX17, TRIB1, APOA1, APOE |
| 41 | 5 | 2.4e-05 | 0.000149993 | Known TG genes | GCKR, VEGFA, TRIB1, APOA1, APOE |

### 6. LocusZoom plots

**6.1. Total Cholesterol LocusZoom plots.**

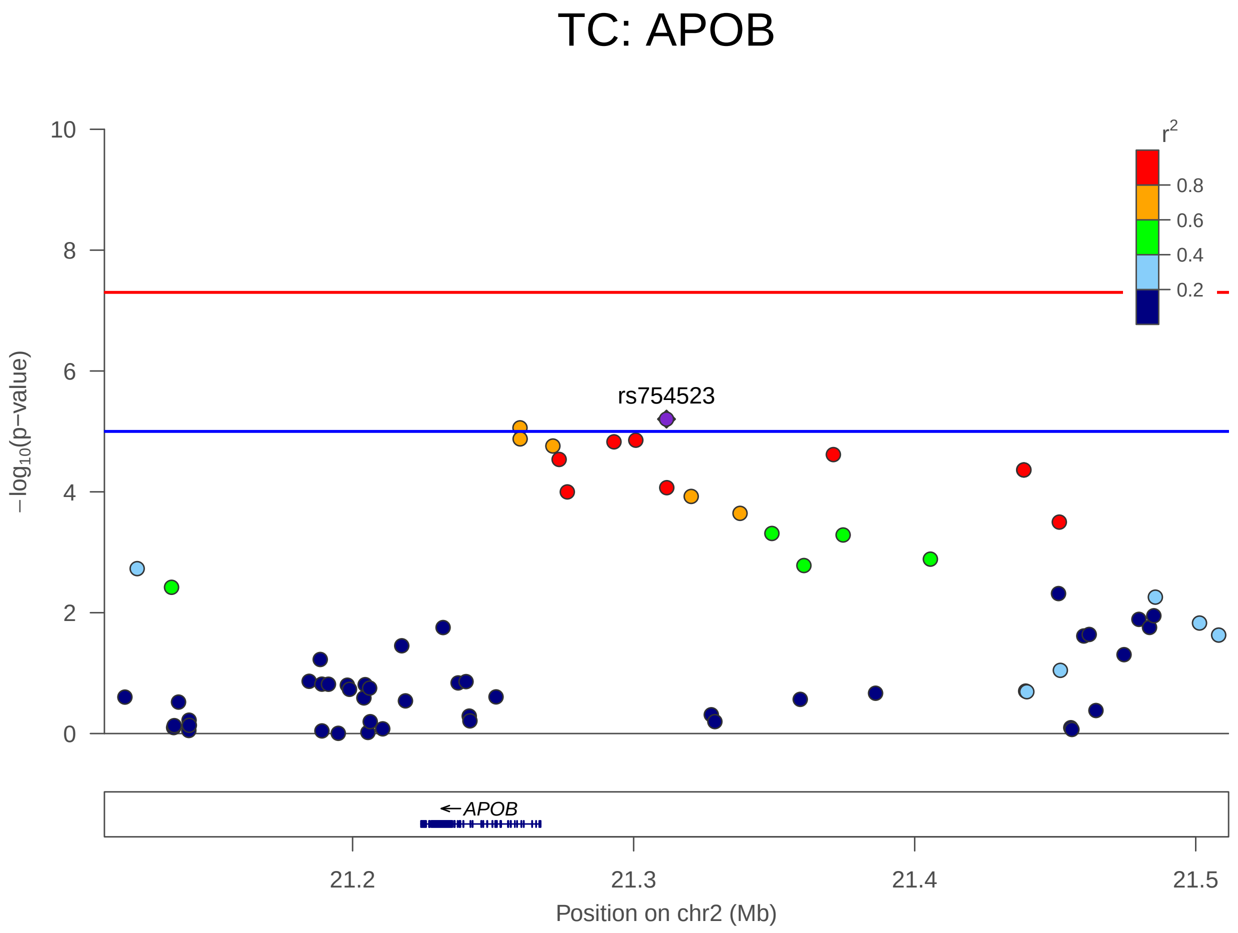


Figure S9. TC on chromosome 2 positions 21111691-21511691

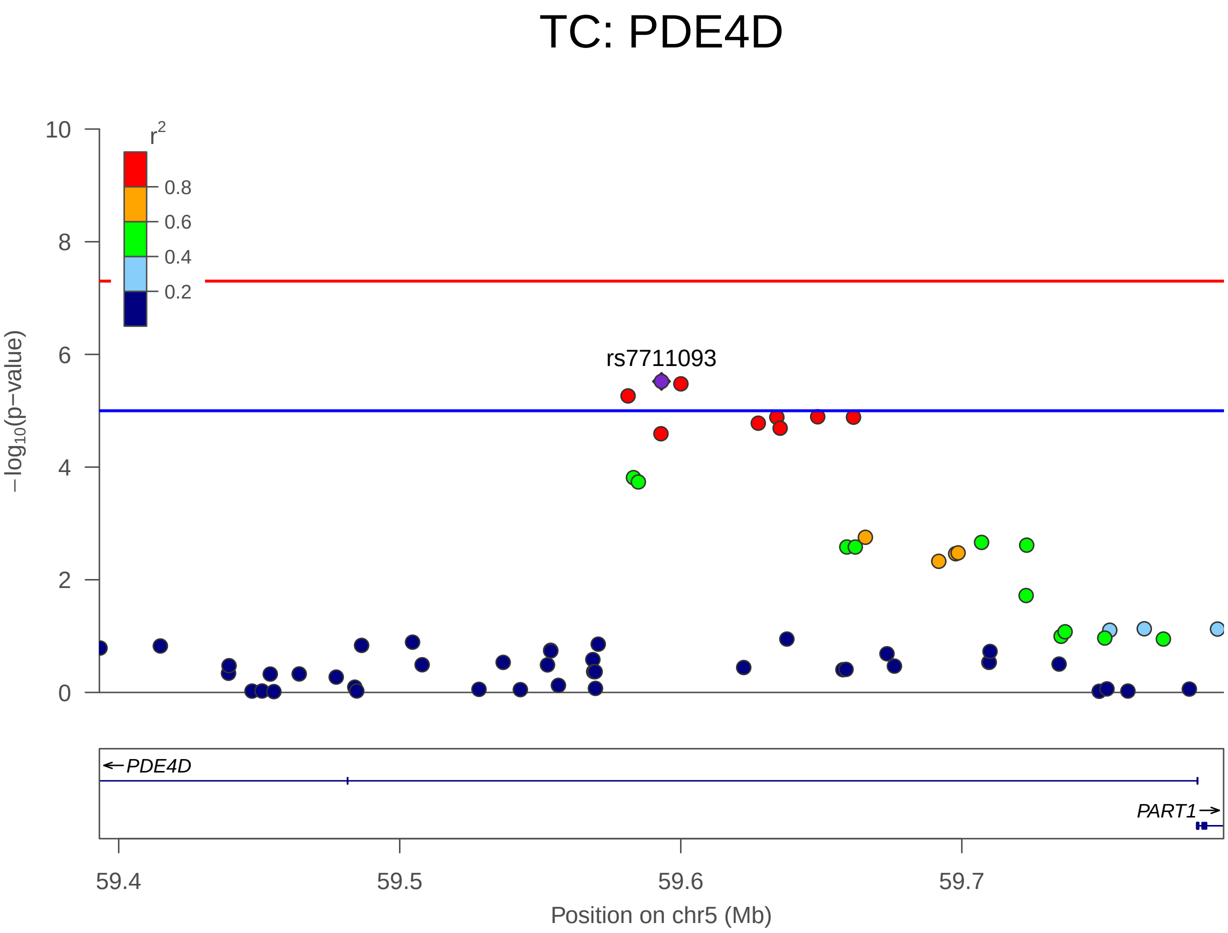


Figure S10. TC on chromosome 5 positions 59393138-59793138

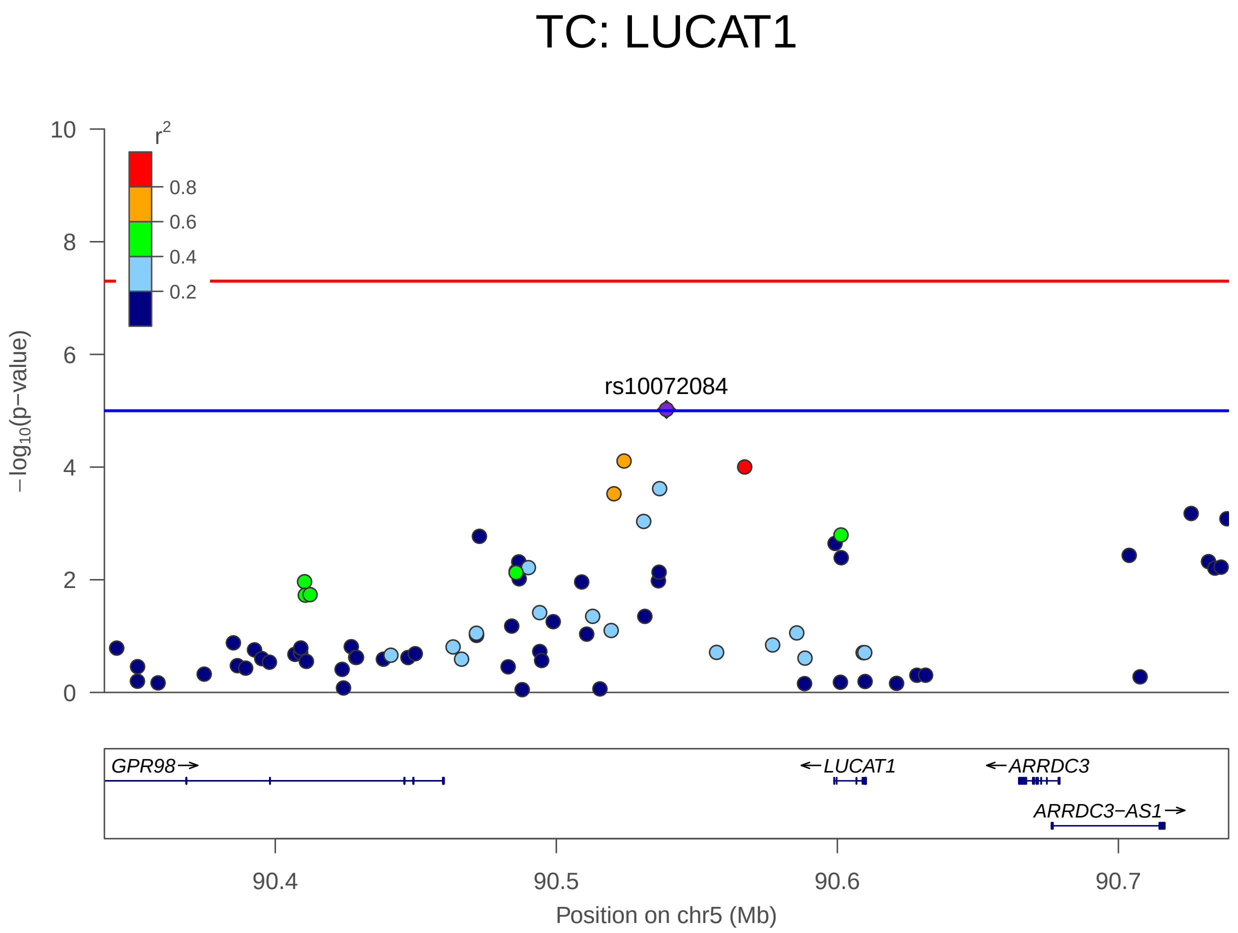


Figure S11. TC on chromosome 5 positions 90339203-90739203

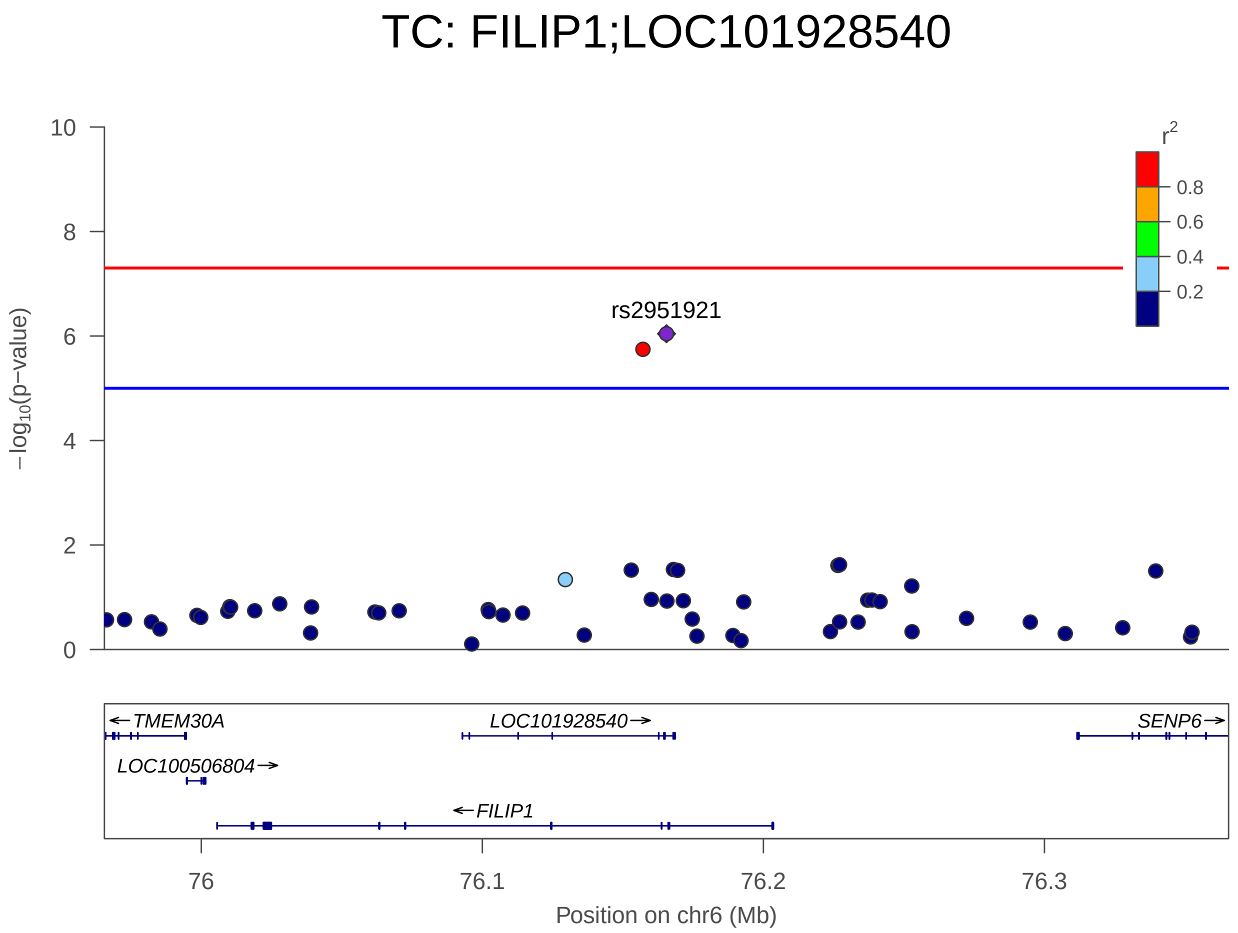


FIGURE S12. TC on chromosome 6 positions 75965524-76365524

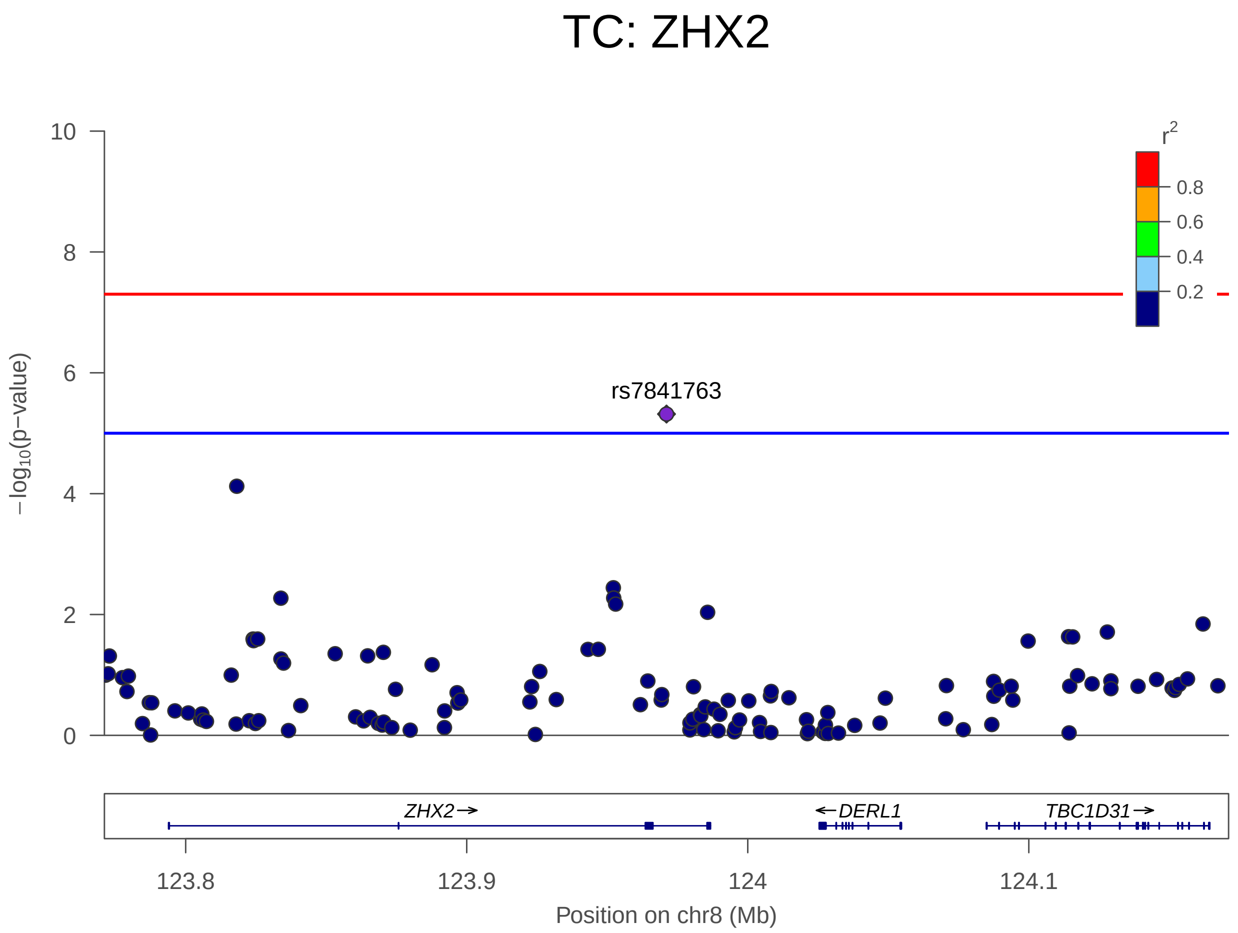


FIGURE S13. TC on chromosome 8 positions 123771081-124171081

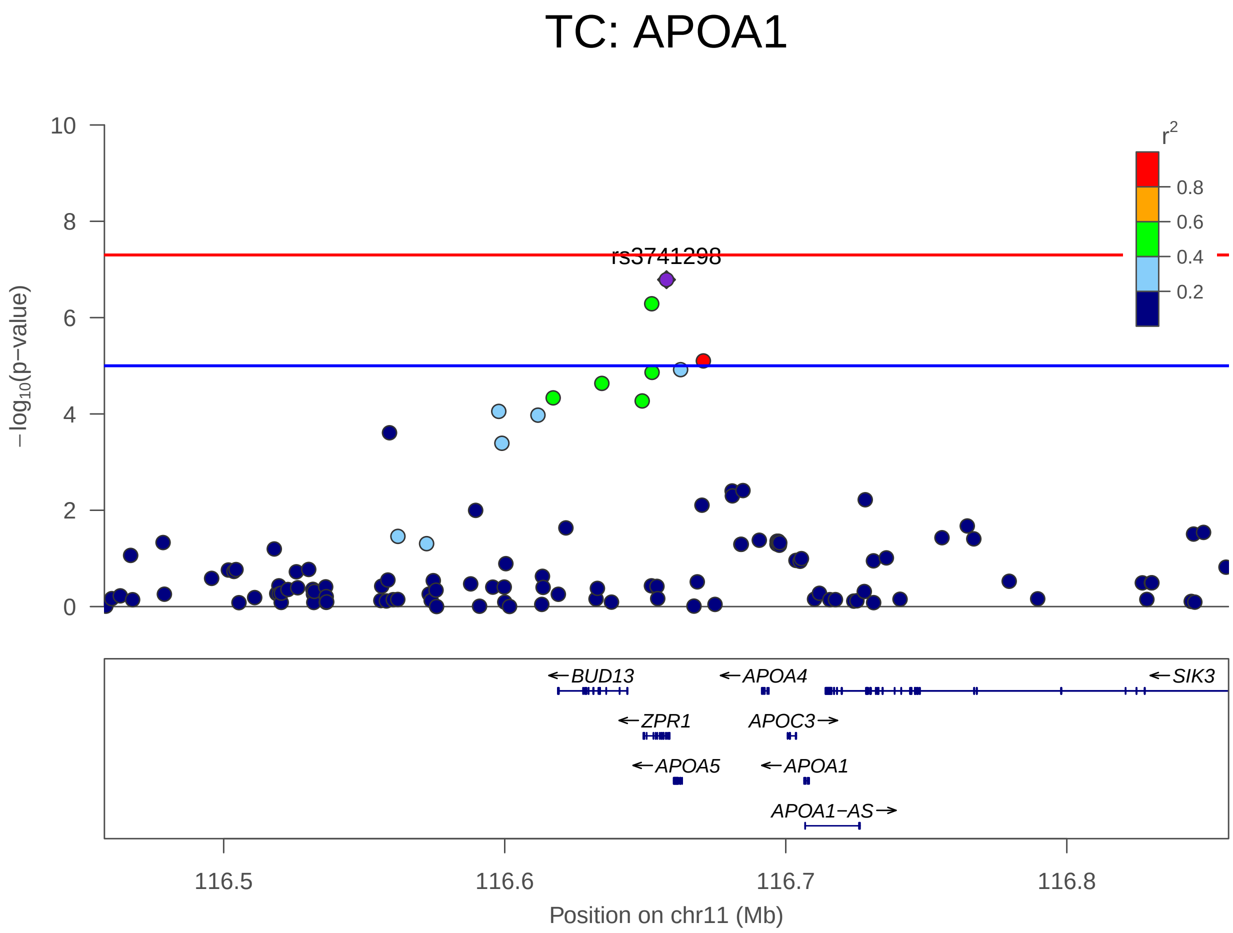


FIGURE S14. TC on chromosome 11 positions 116457561-116857561

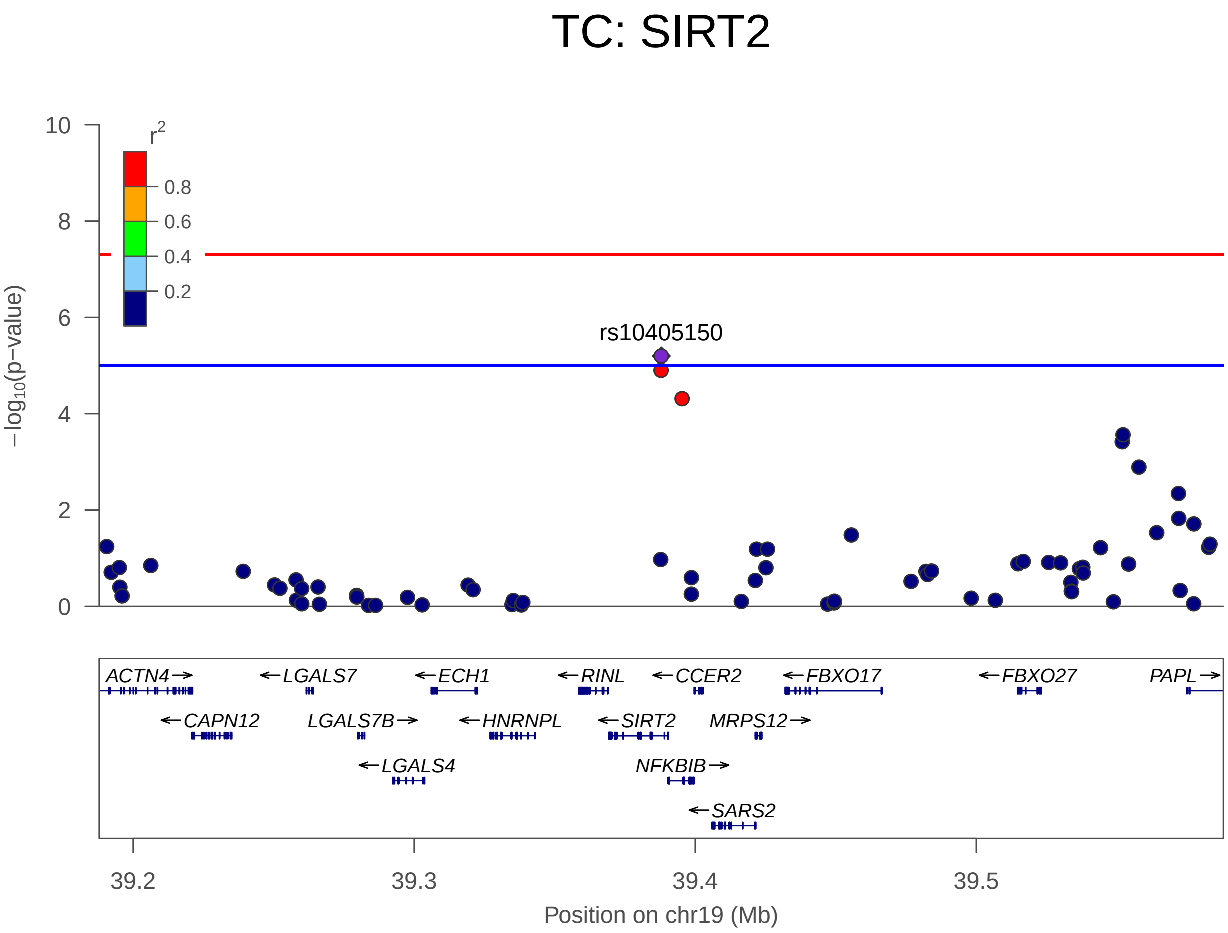


FIGURE S15. TC on chromosome 19 positions 39187919-39587919

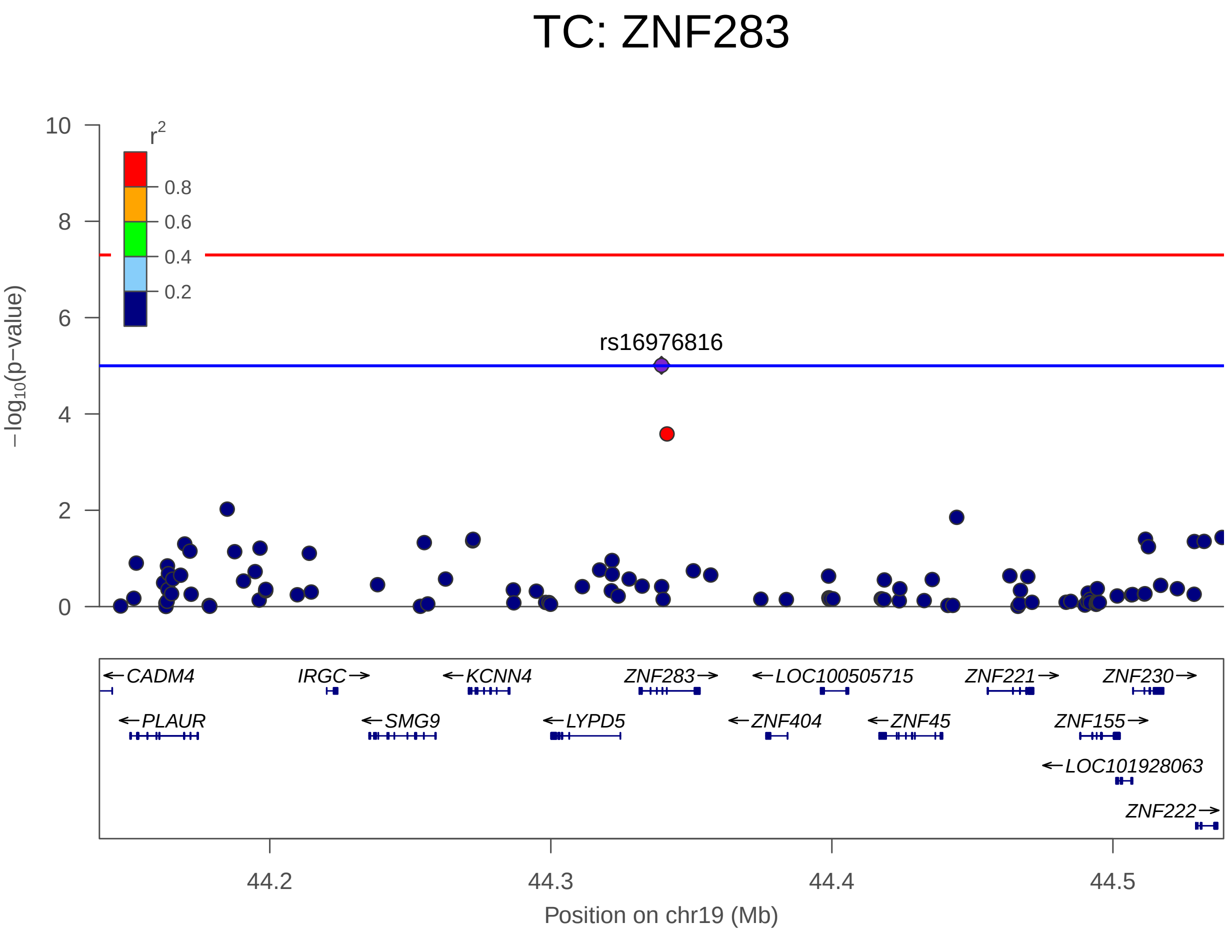


FIGURE S16. TC on chromosome 19 positions 44139377-44539377

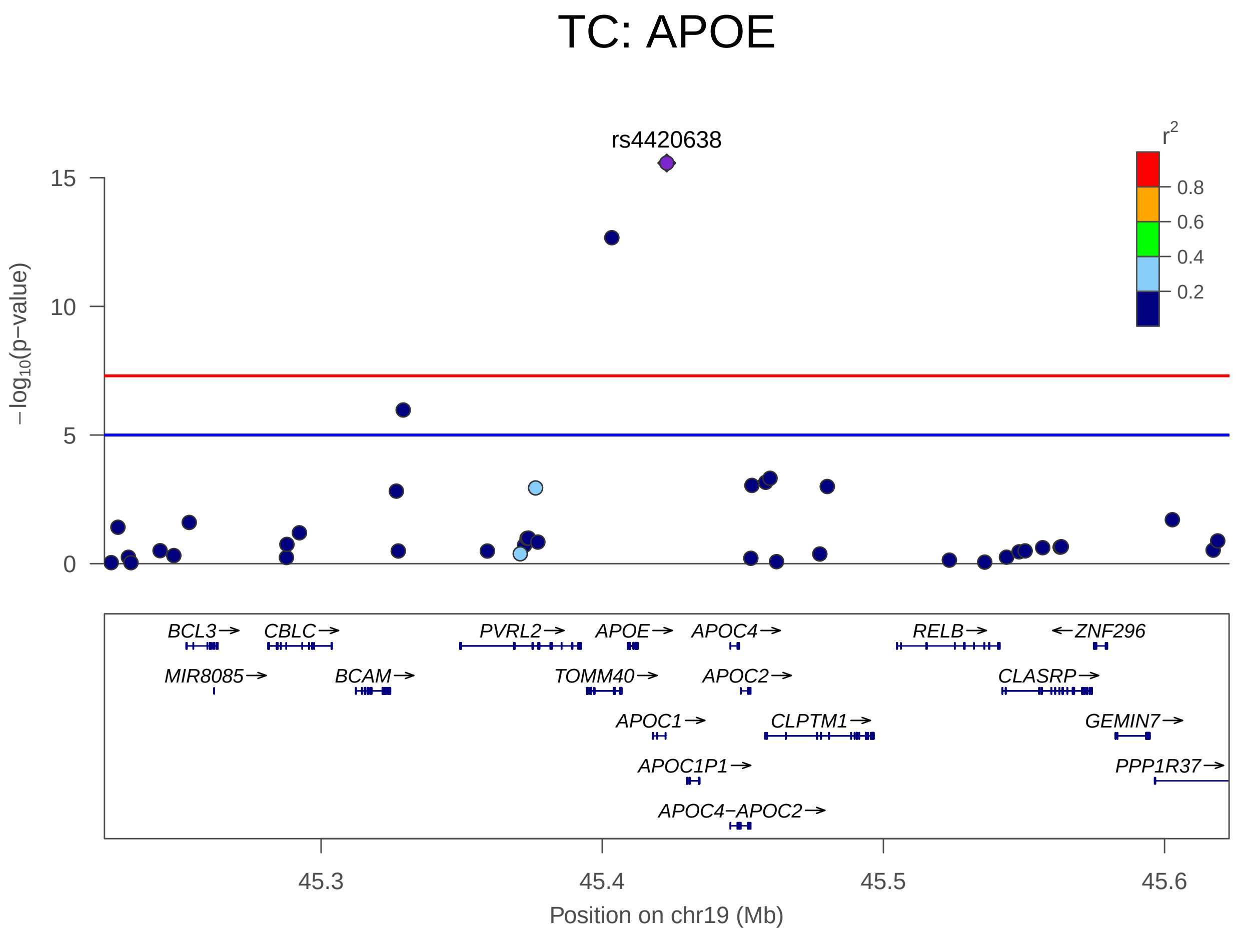


Figure S17. TC on chromosome 19 positions 45222946-45622946

### 6.2. **HDL LocusZoom plots.**

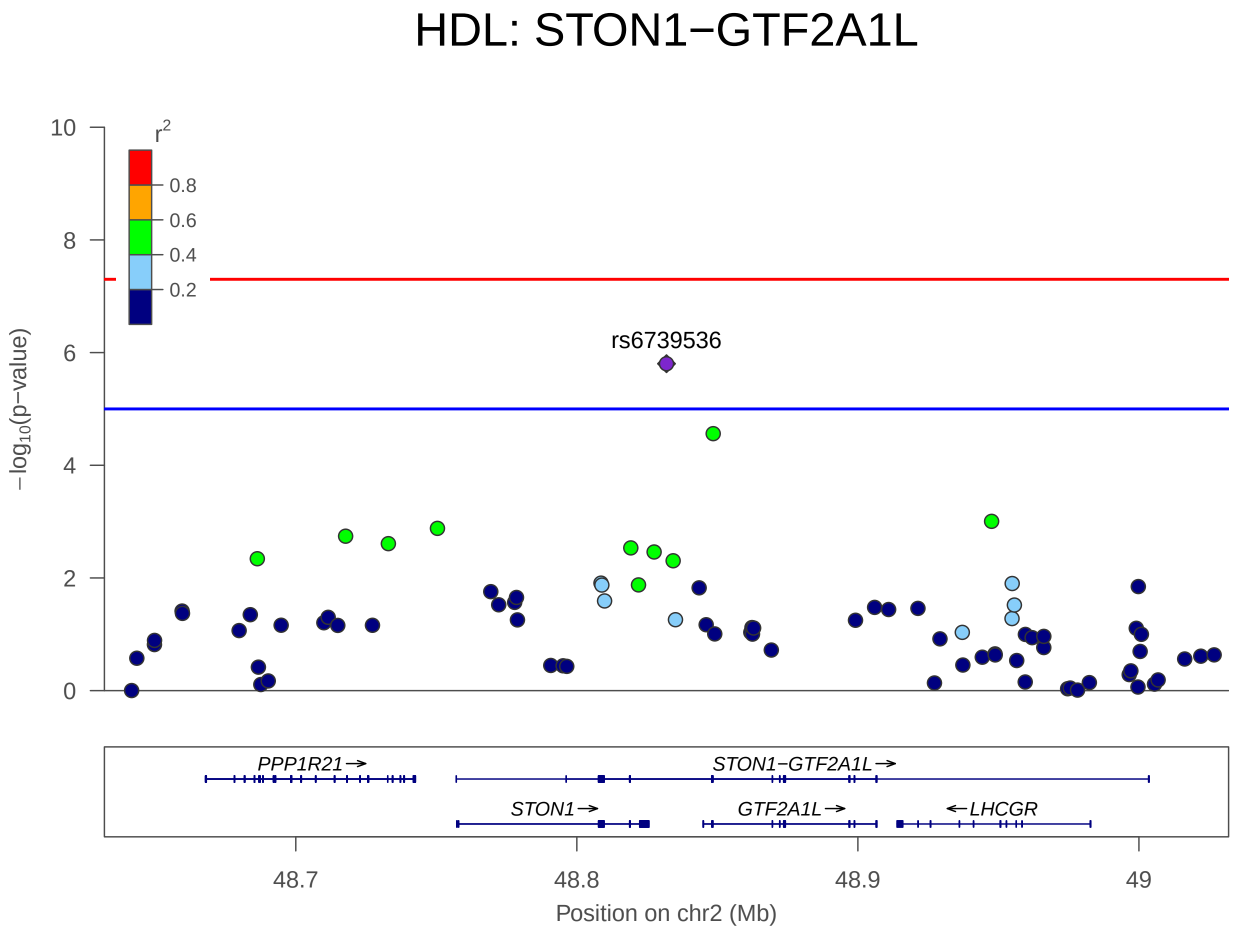


FIGURE S18. HDL on chromosome 2 positions 48631901-49031901

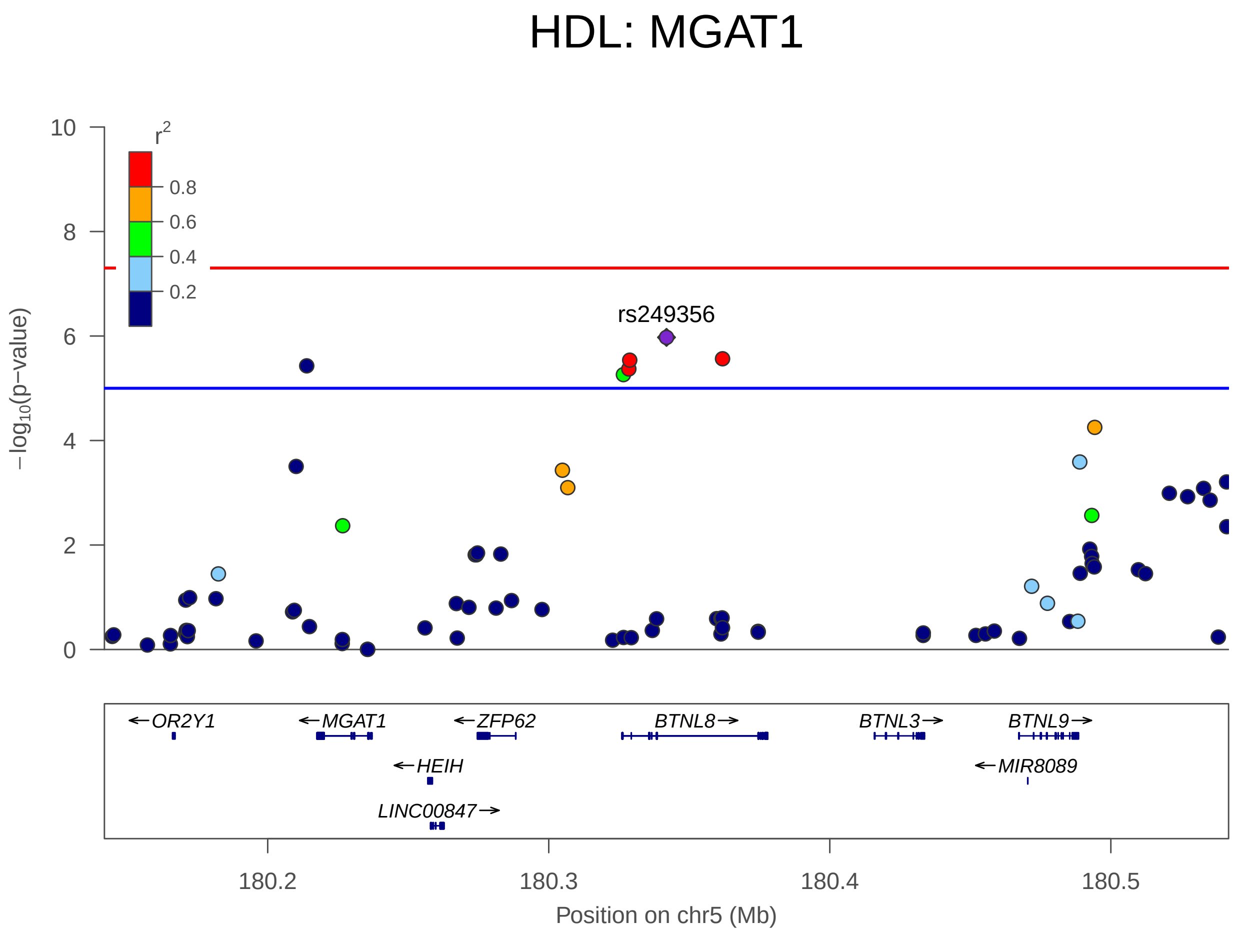


FIGURE S19. HDL on chromosome 5 positions 180141895-180541895

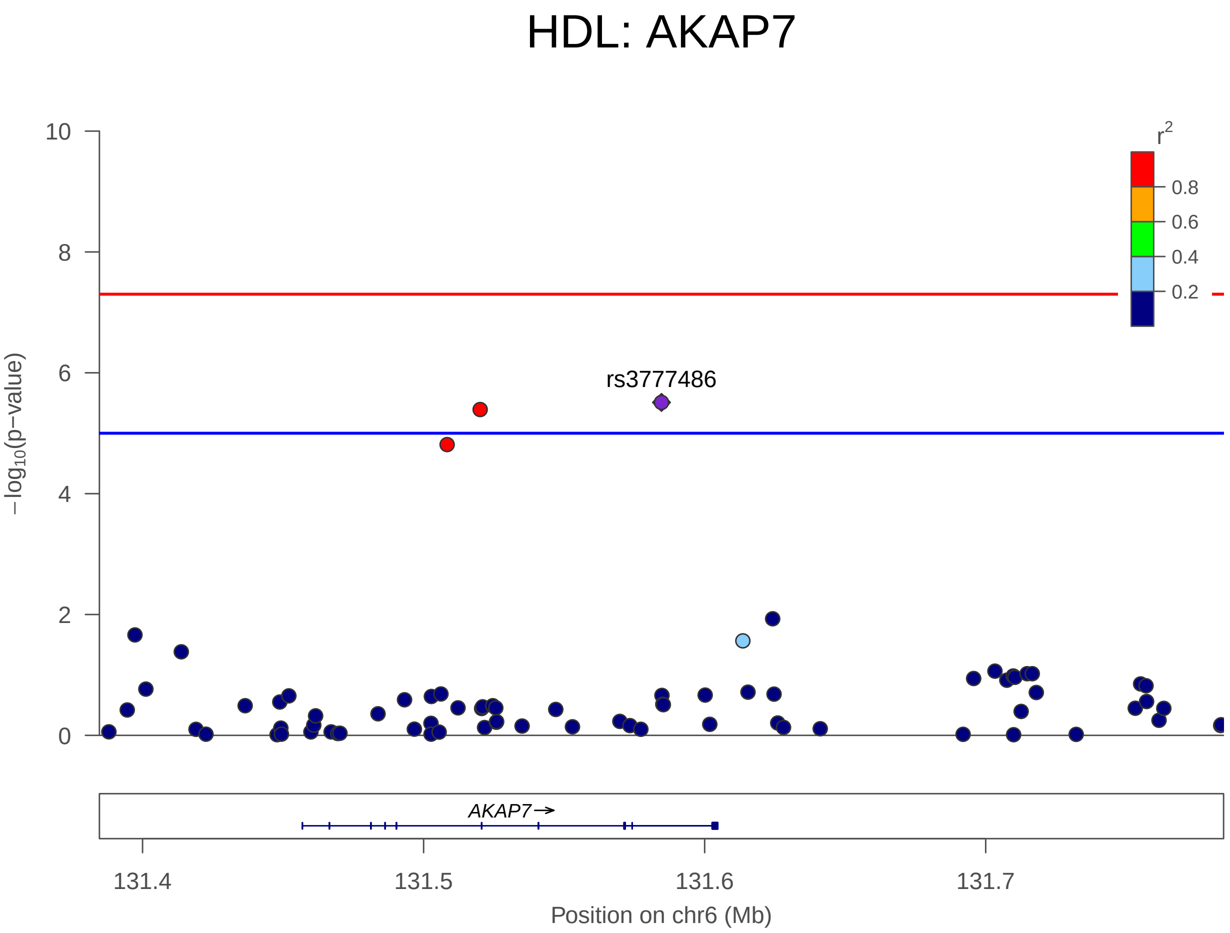


Figure S20. HDL on chromosome 6 positions 131384648-131784648

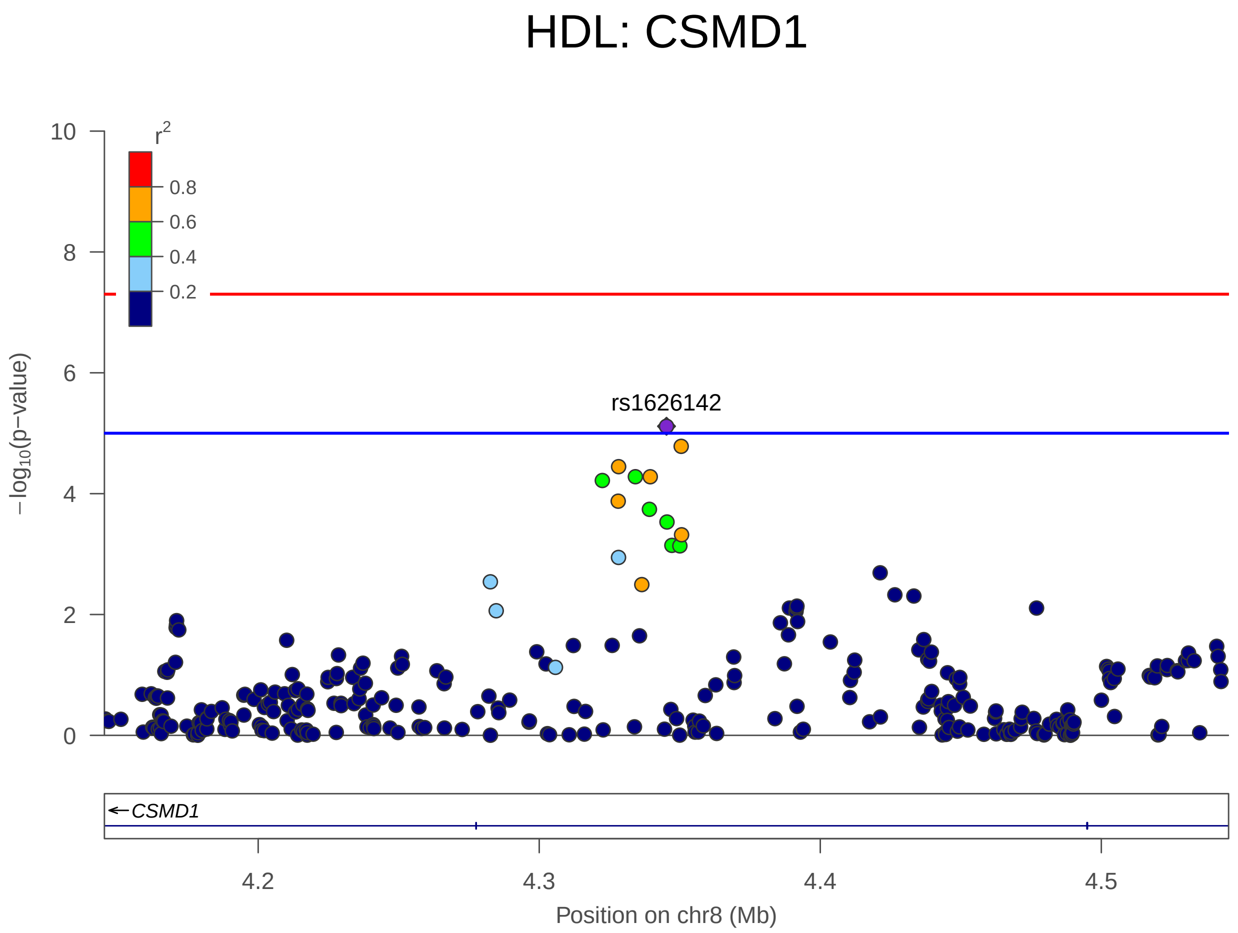


FIGURE S21. HDL on chromosome 8 positions 4145284-4545284

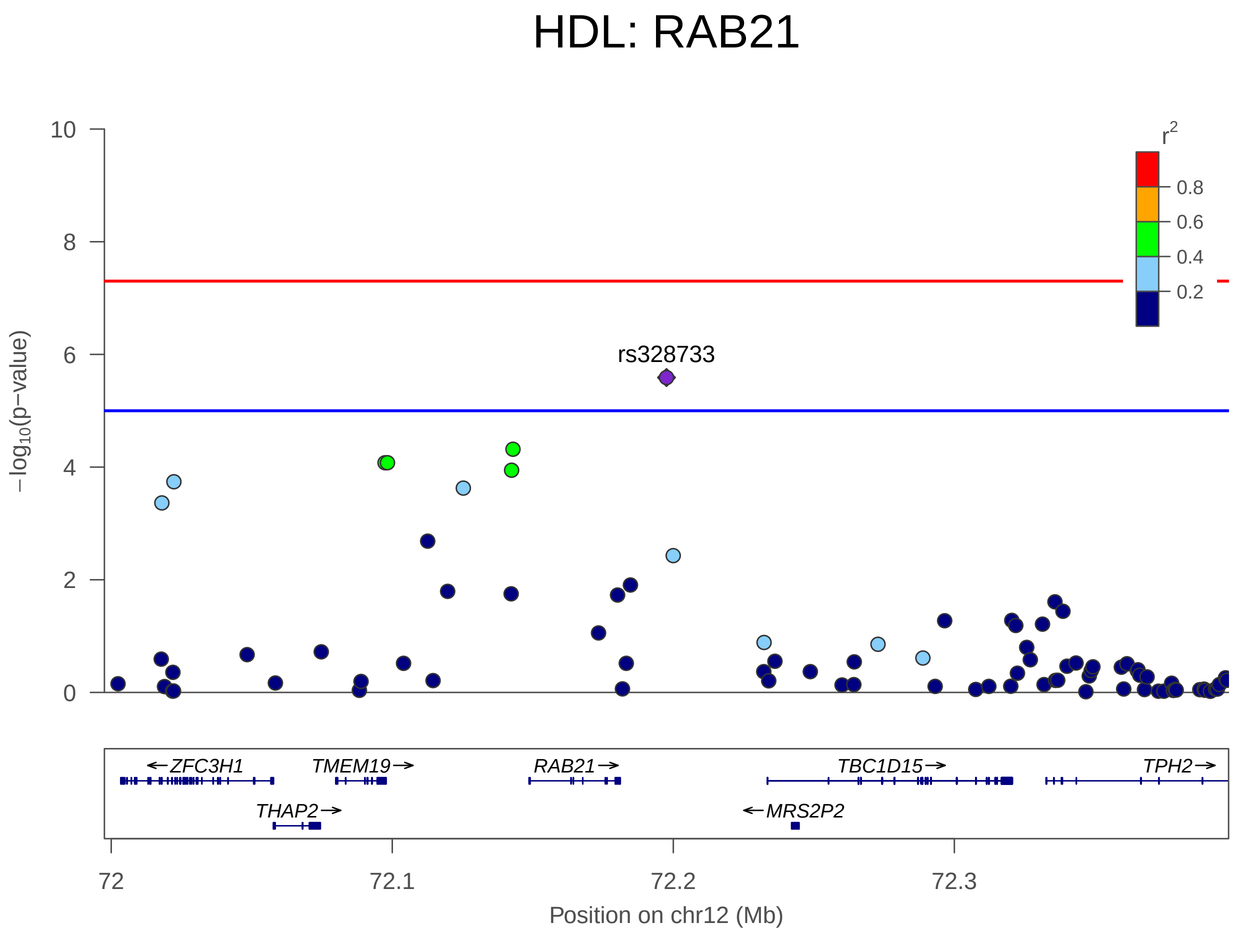


FIGURE S22. HDL on chromosome 12 positions 71997574-72397574

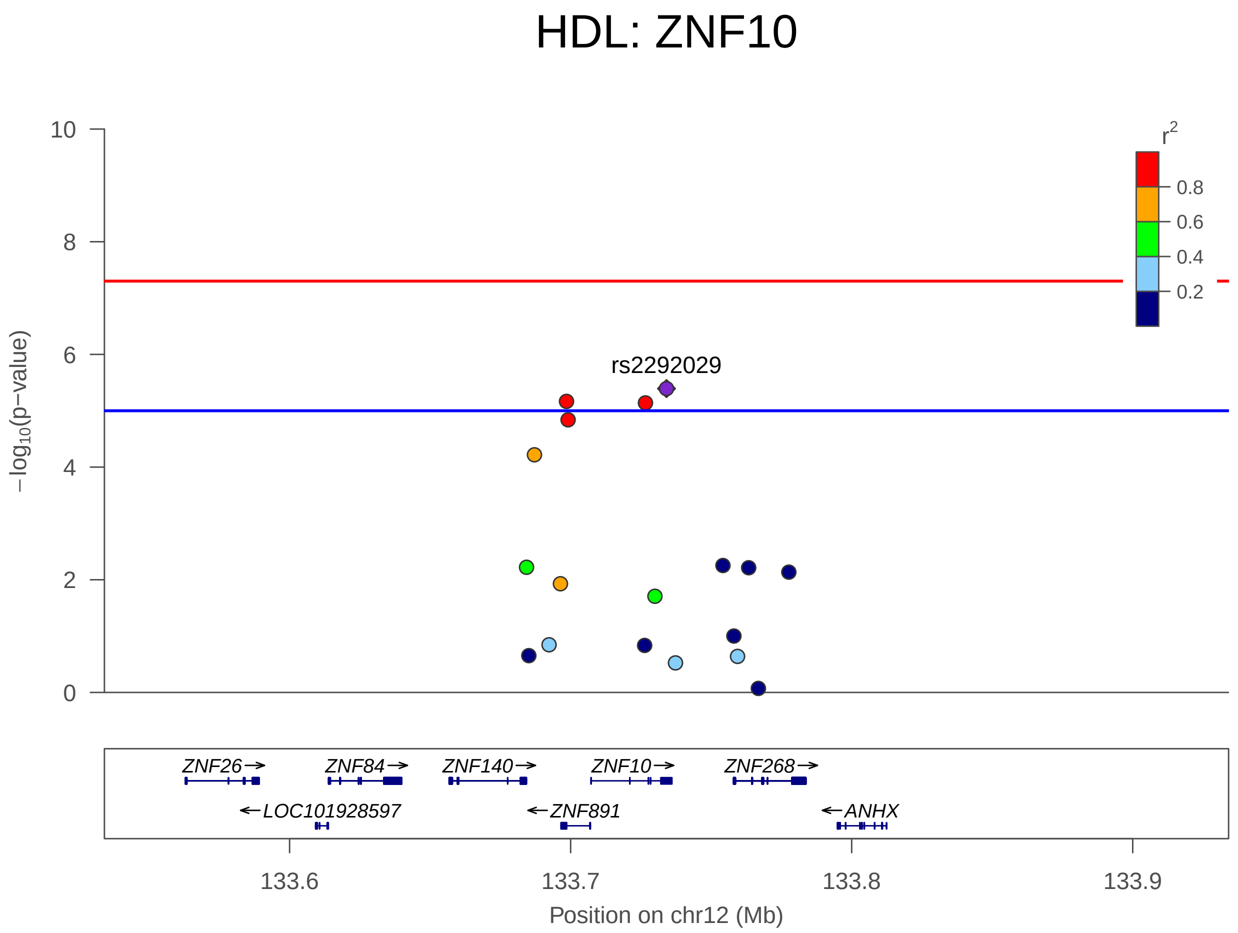


FIGURE S23. HDL on chromosome 12 positions 133534113-133934113

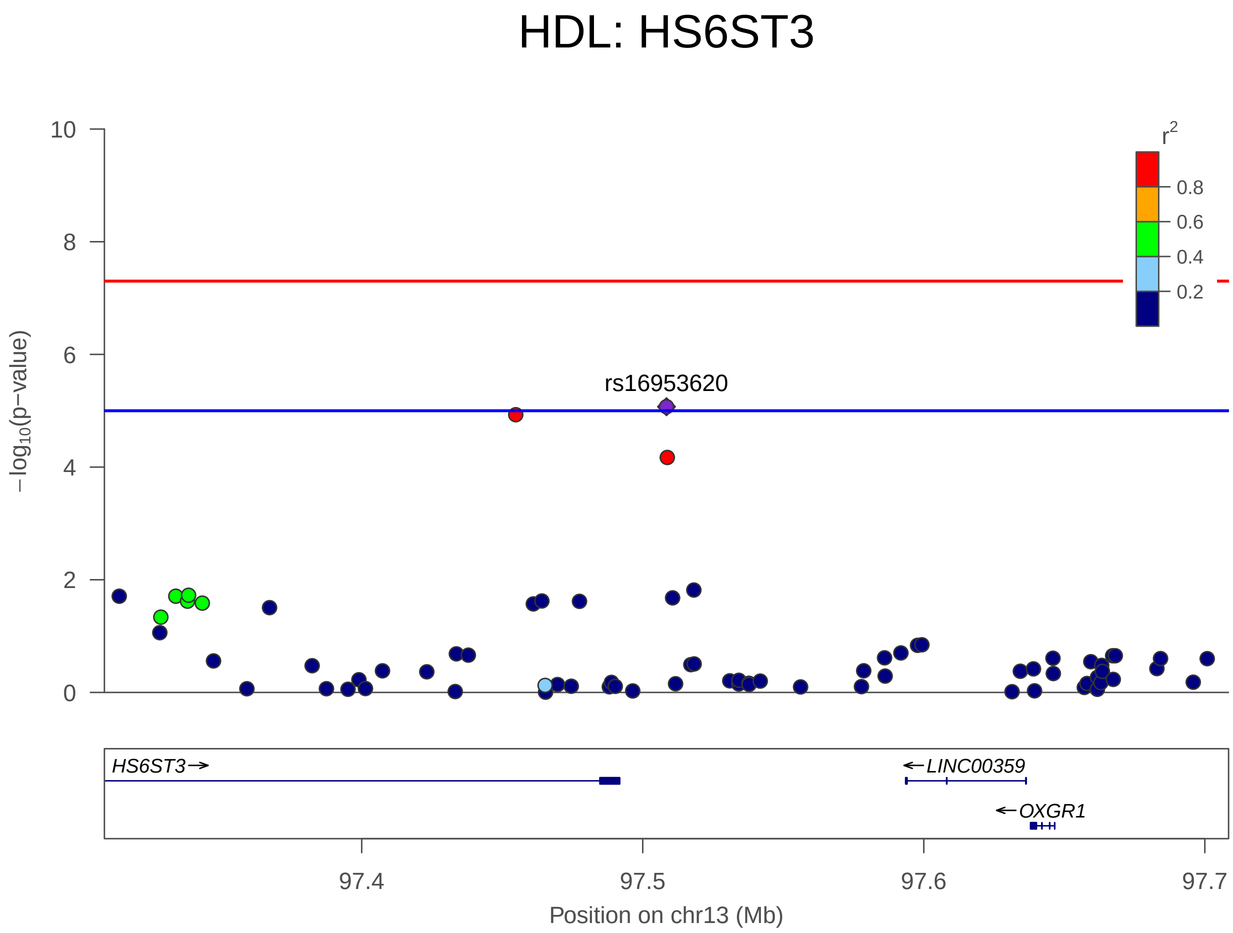


Figure S24. HDL on chromosome 13 positions 97308453-97708453

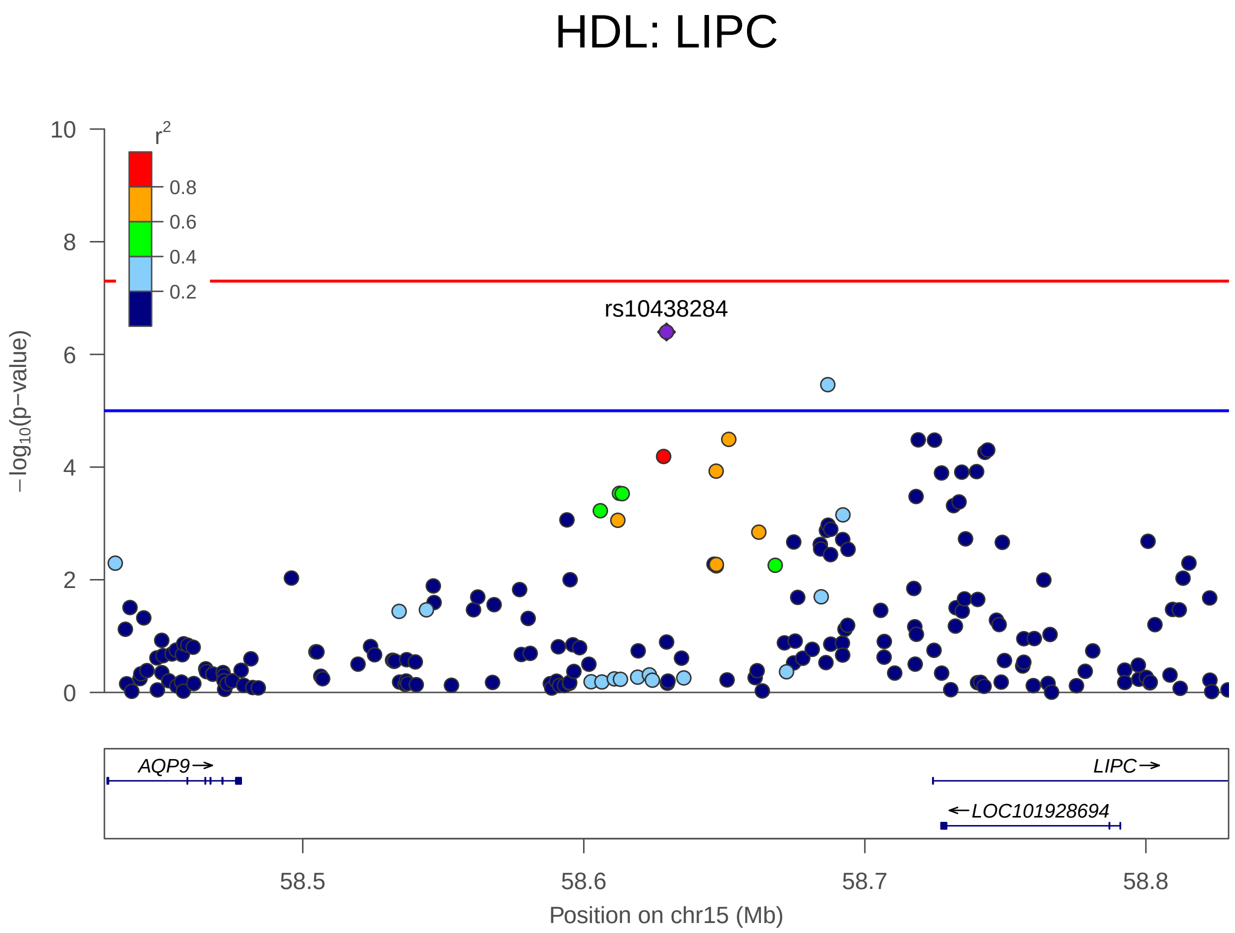


FIGURE S25. HDL on chromosome 15 positions 58429424-58829424

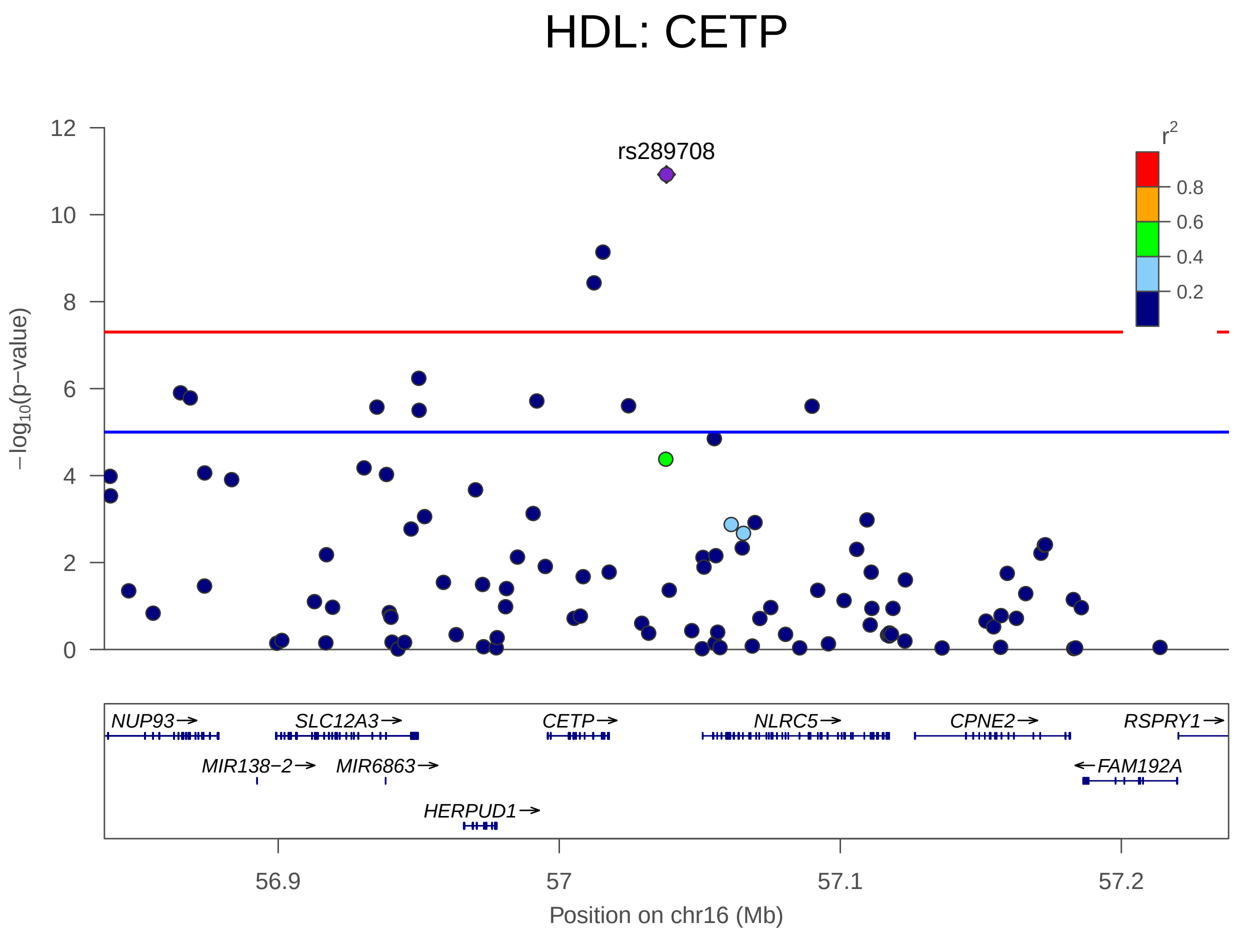


Figure S26. HDL on chromosome 16 positions 56838162-57238162

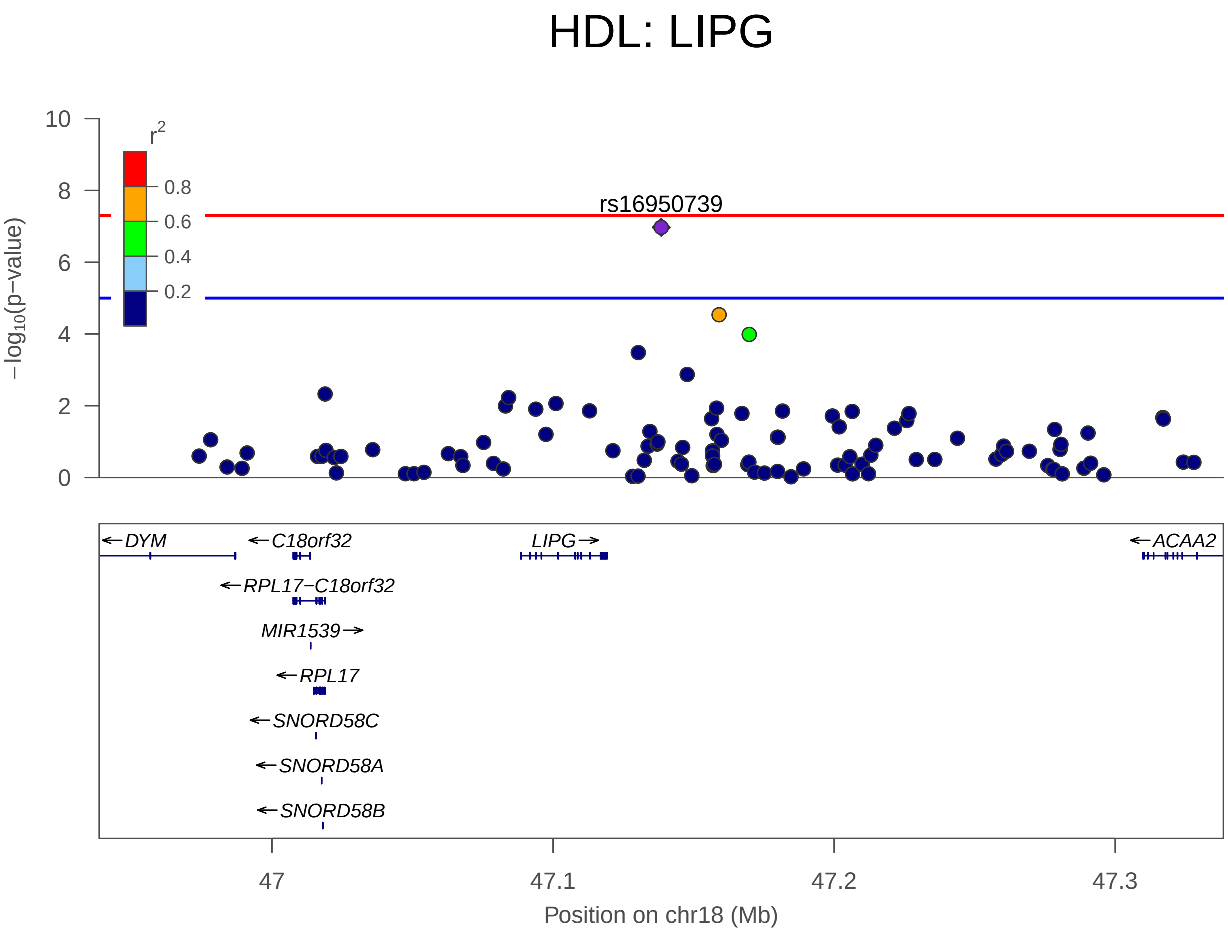


Figure S27. HDL on chromosome 18 positions 46938509-47338509

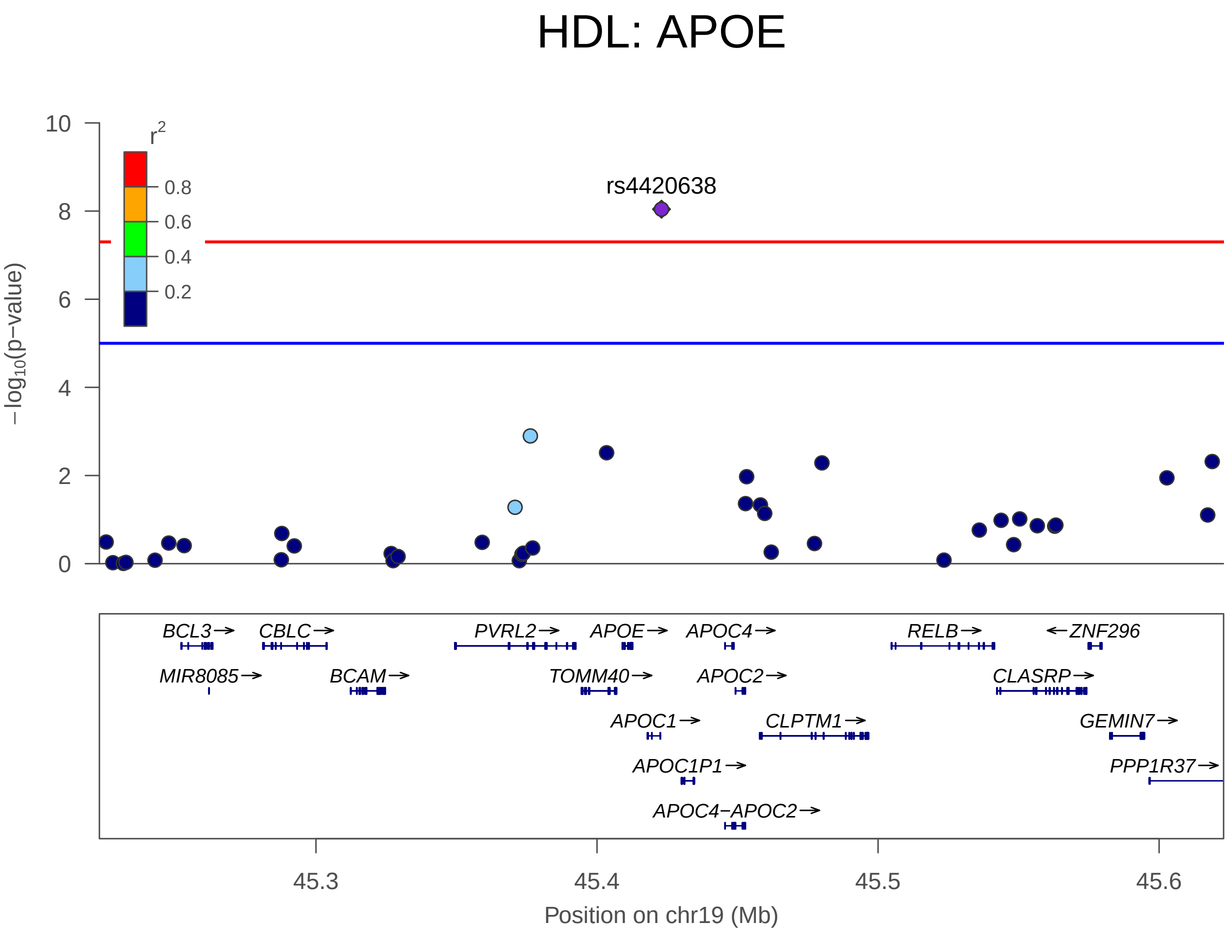


FIGURE S28. HDL on chromosome 19 positions 45222946-45622946

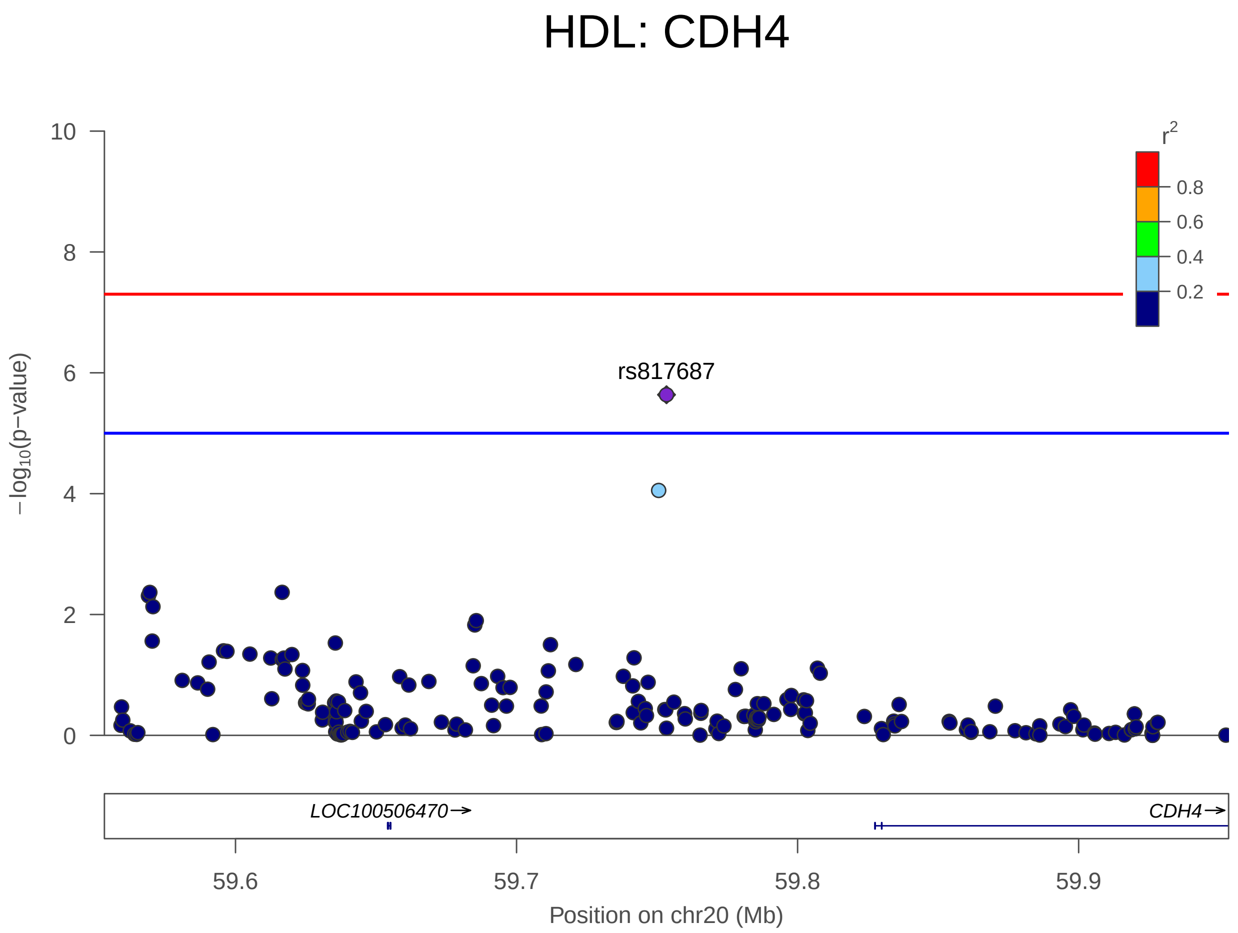


FIGURE S29. HDL on chromosome 20 positions 59553355-59953355

6.3. **LDL LocusZoom plots.**

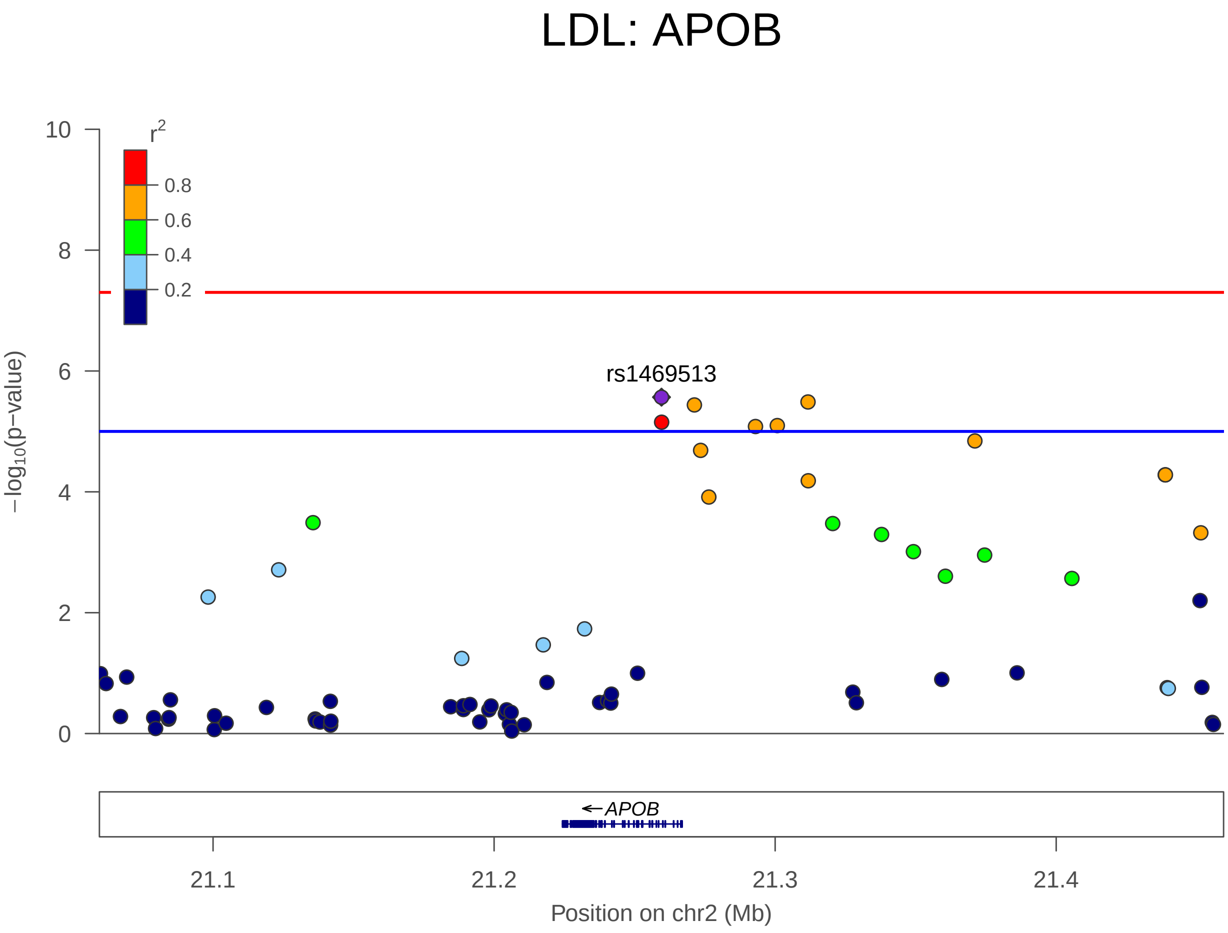


Figure S30. LDL on chromosome 2 positions 21059562-21459562

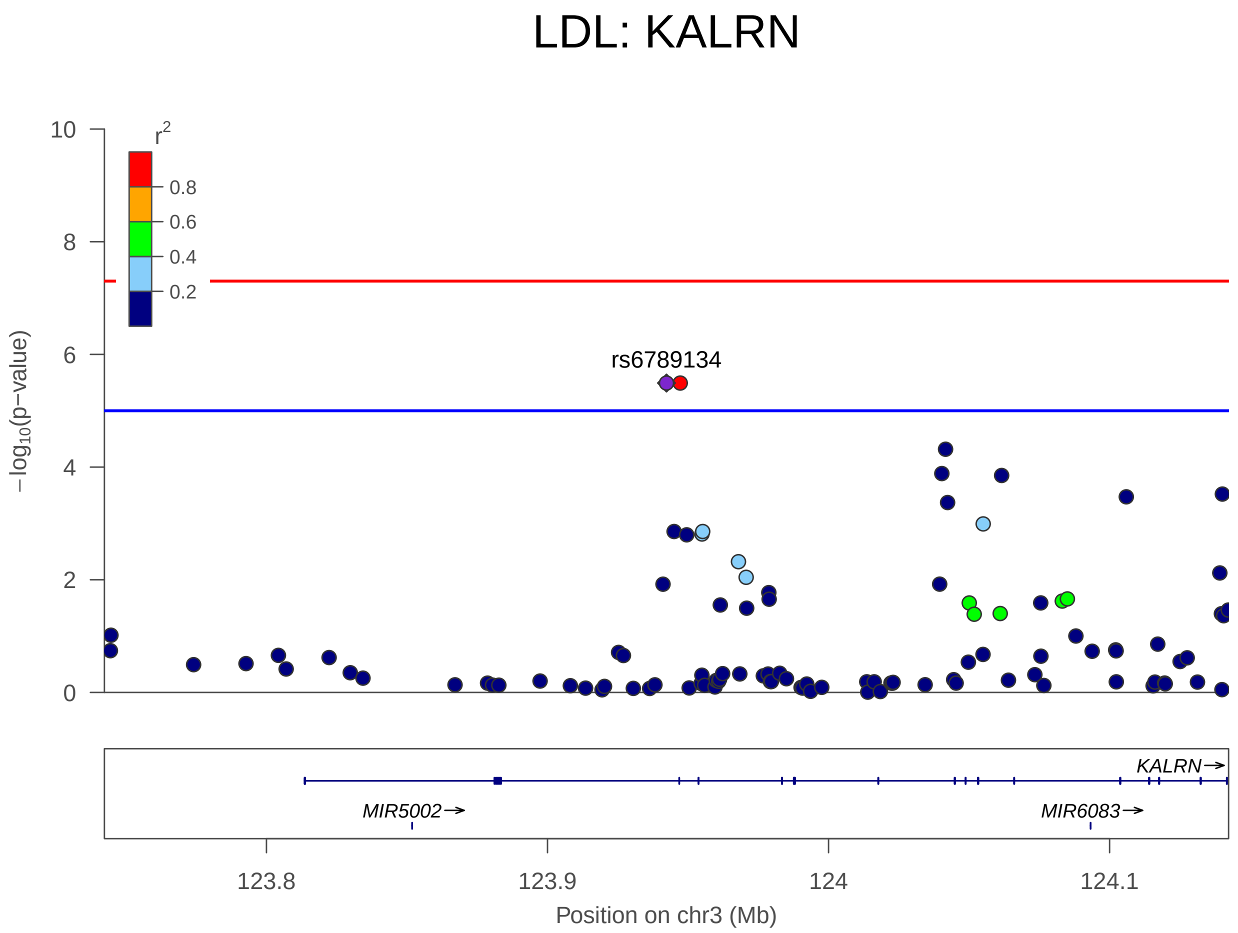


Figure S31. LDL on chromosome 3 positions 123742339-124142339

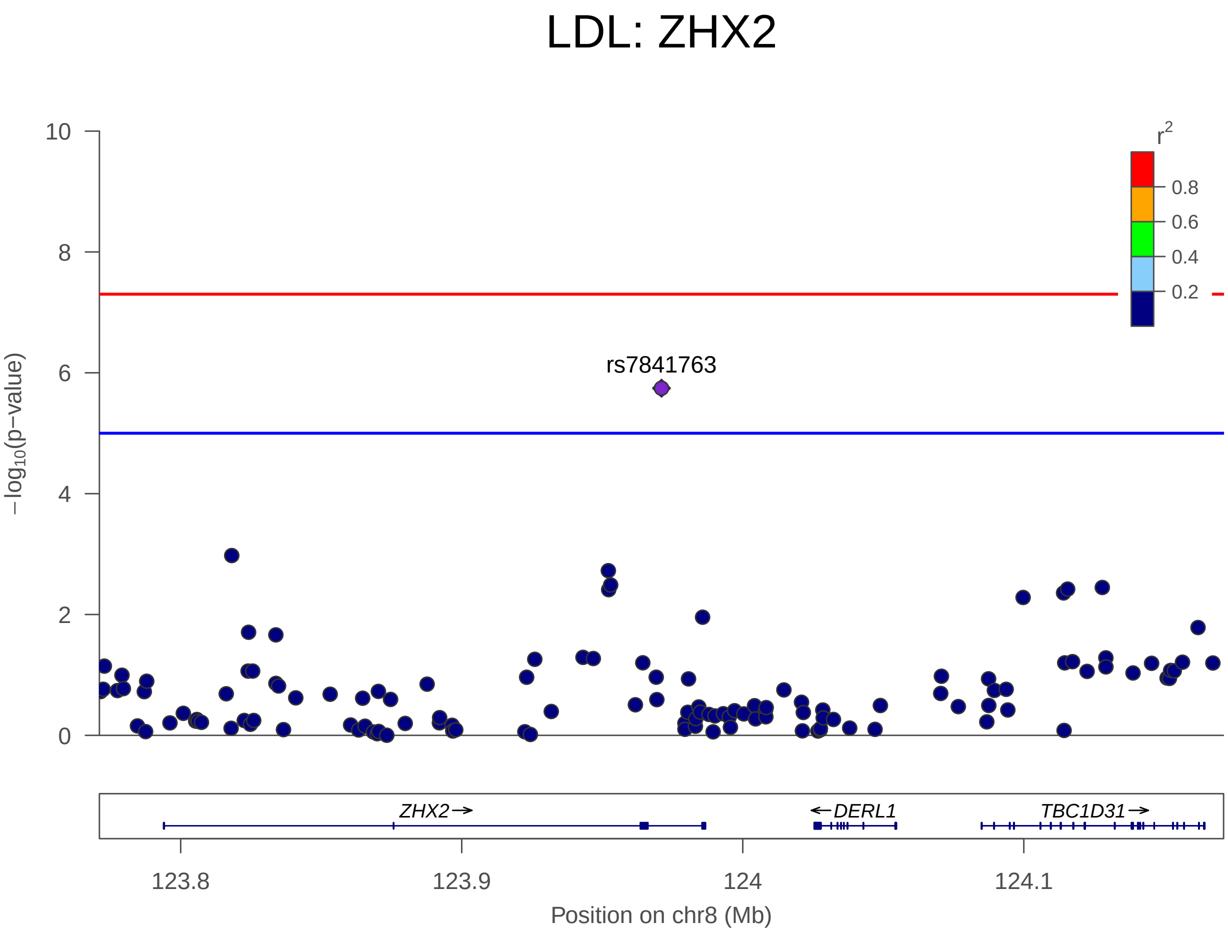


Figure S32. LDL on chromosome 8 positions 123771081-124171081

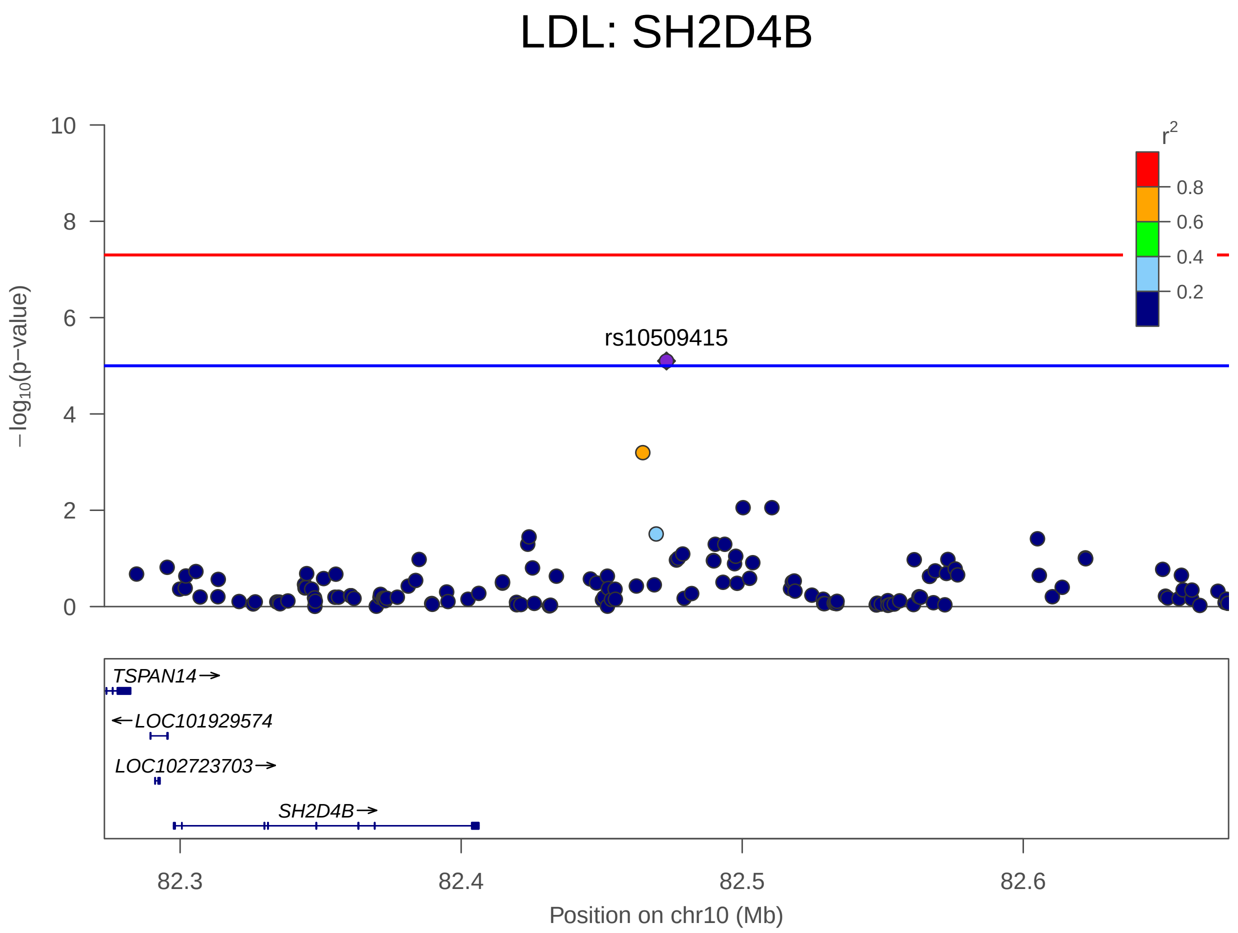


Figure S33. LDL on chromosome 10 positions 82273065-82673065

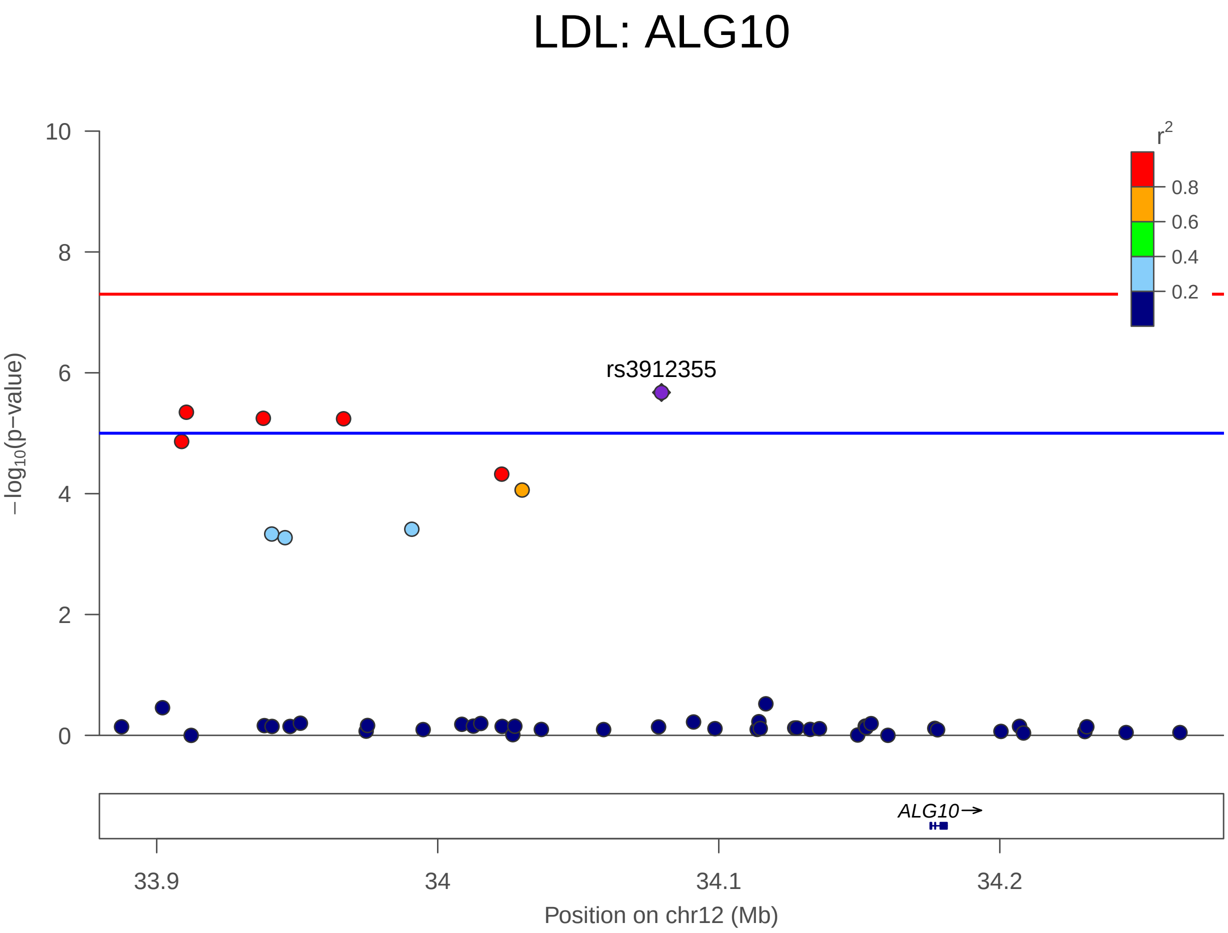


Figure S34. LDL on chromosome 12 positions 33879616-34279616

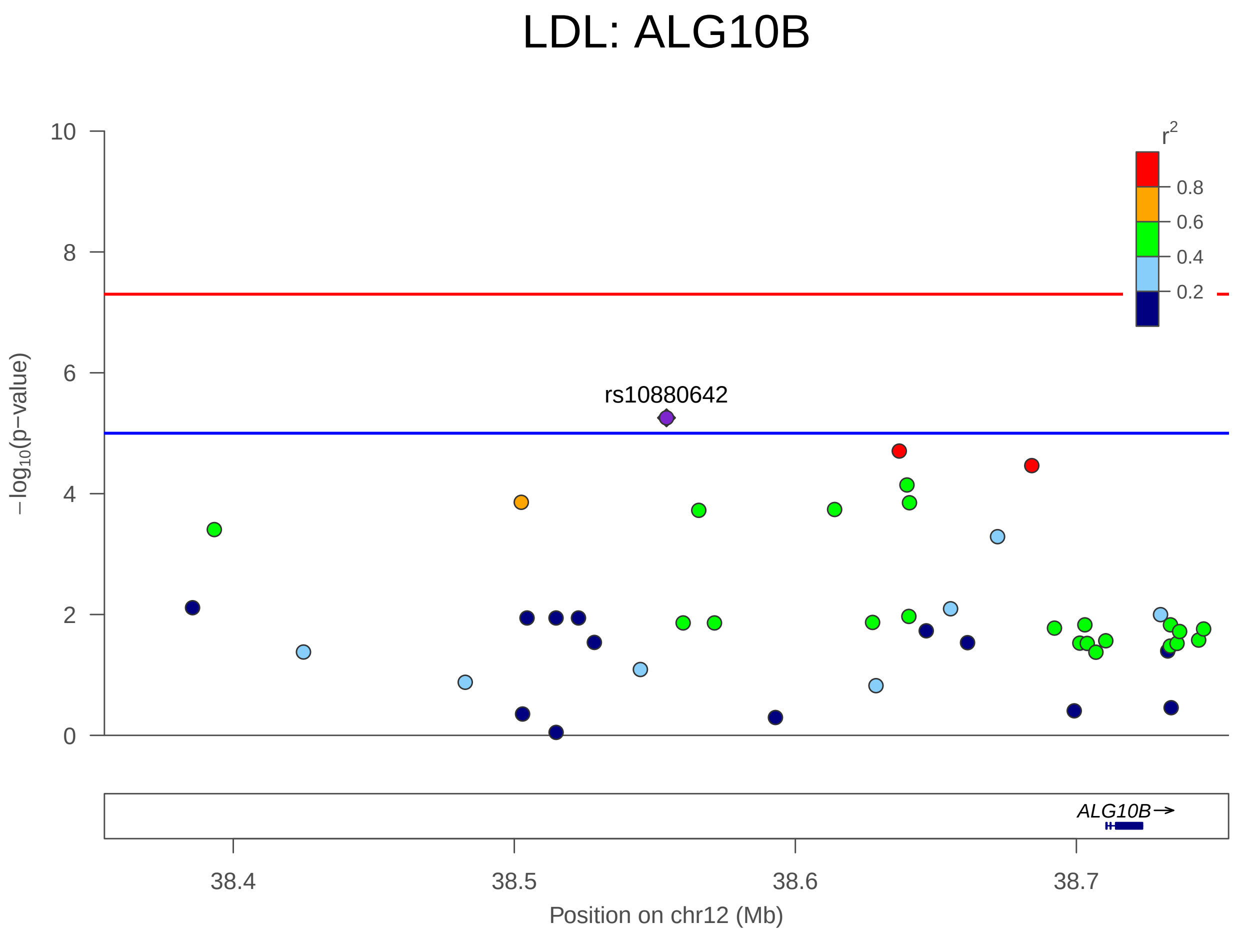


Figure S35. LDL on chromosome 12 positions 38354152-38754152

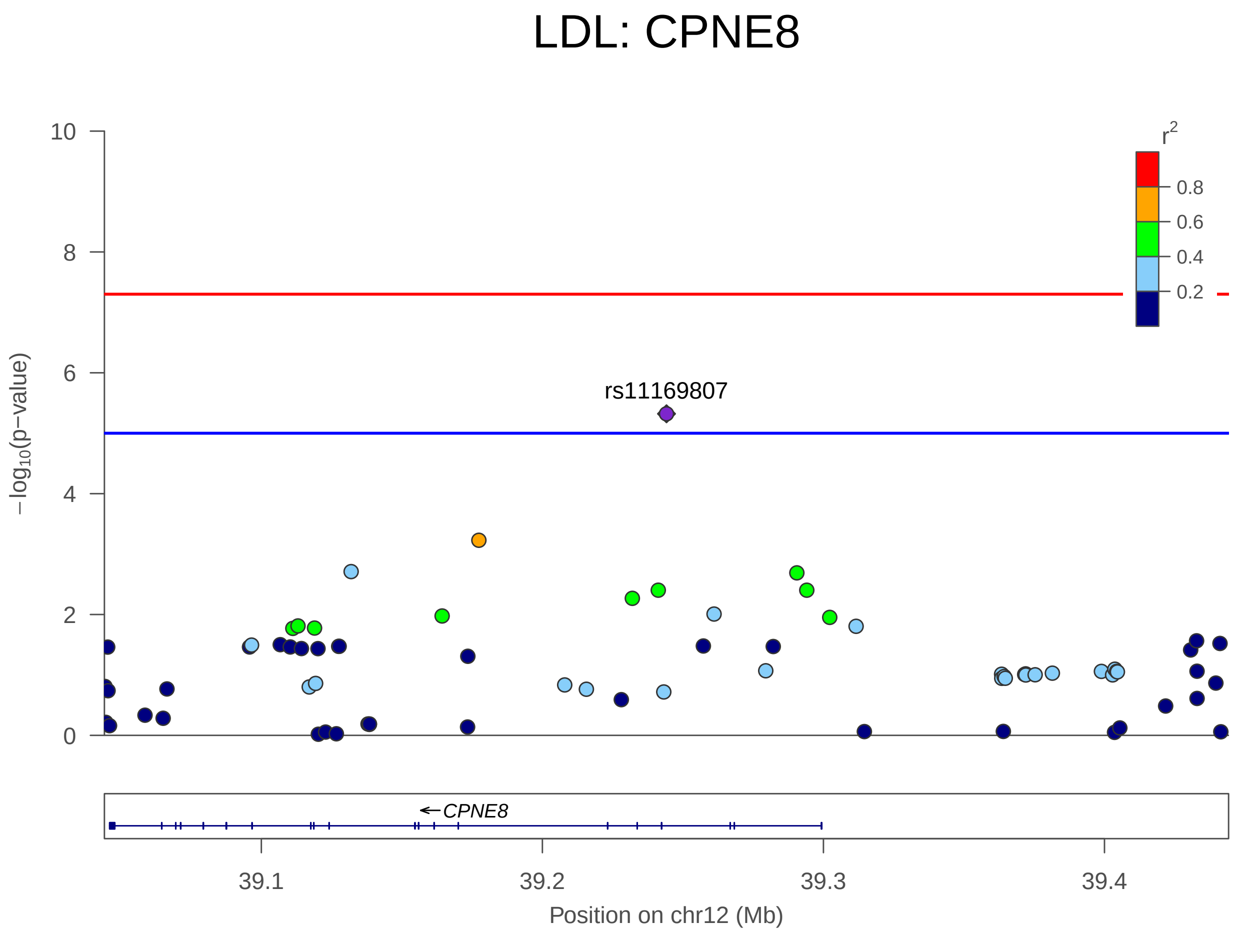


FIGURE S36. LDL on chromosome 12 positions 39044161-39444161

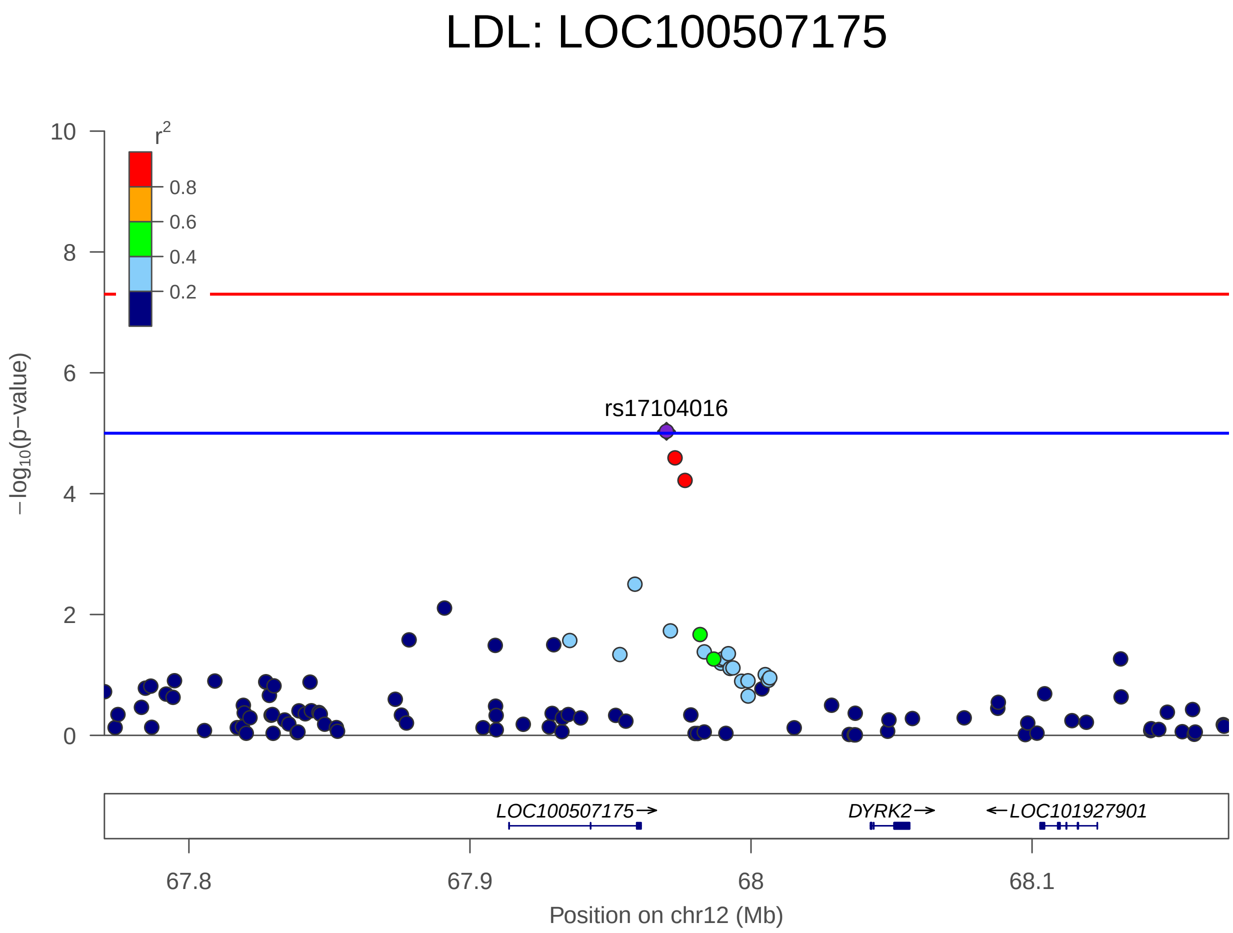


FIGURE S37. LDL on chromosome 12 positions 67769929-68169929

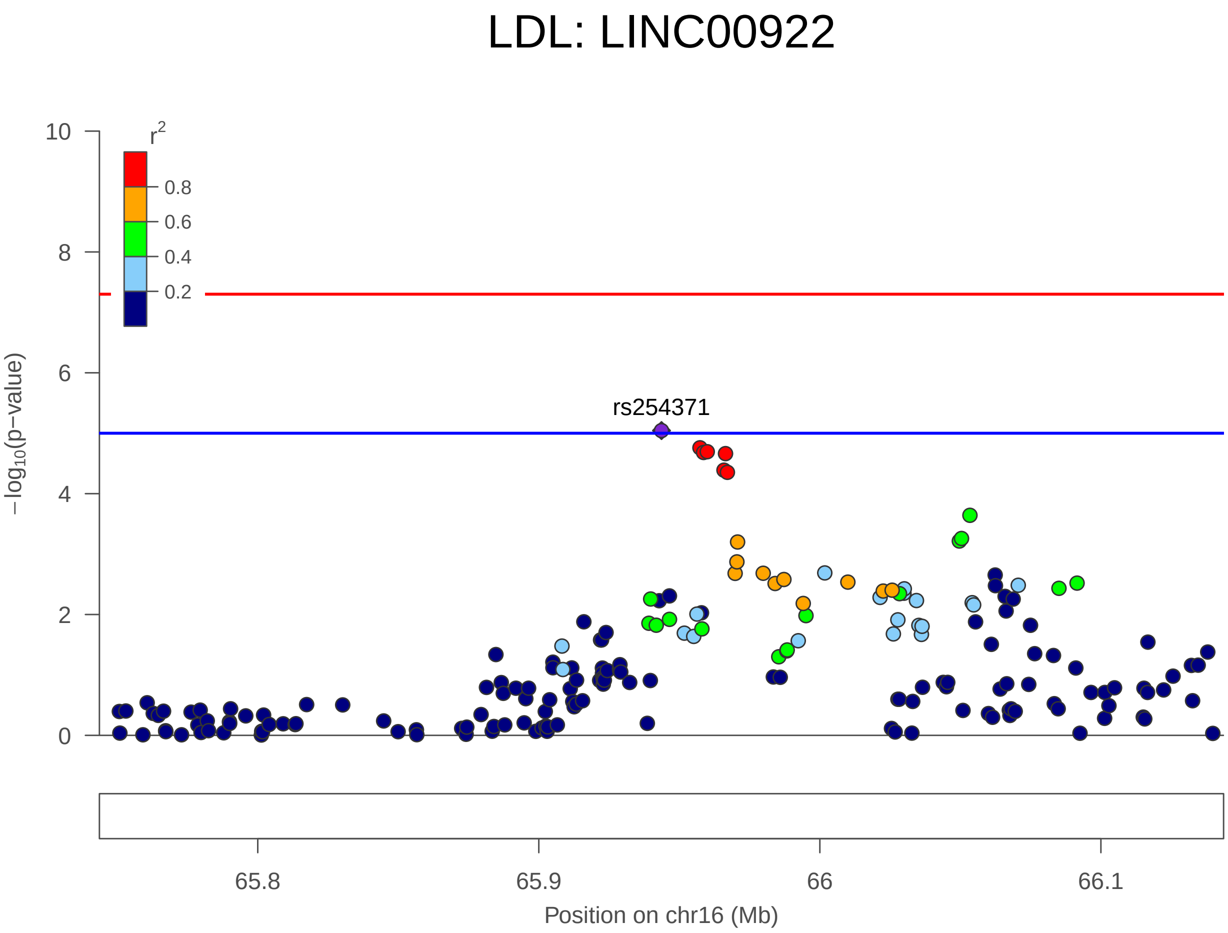


FIGURE S38. LDL on chromosome 16 positions 65743650-66143650

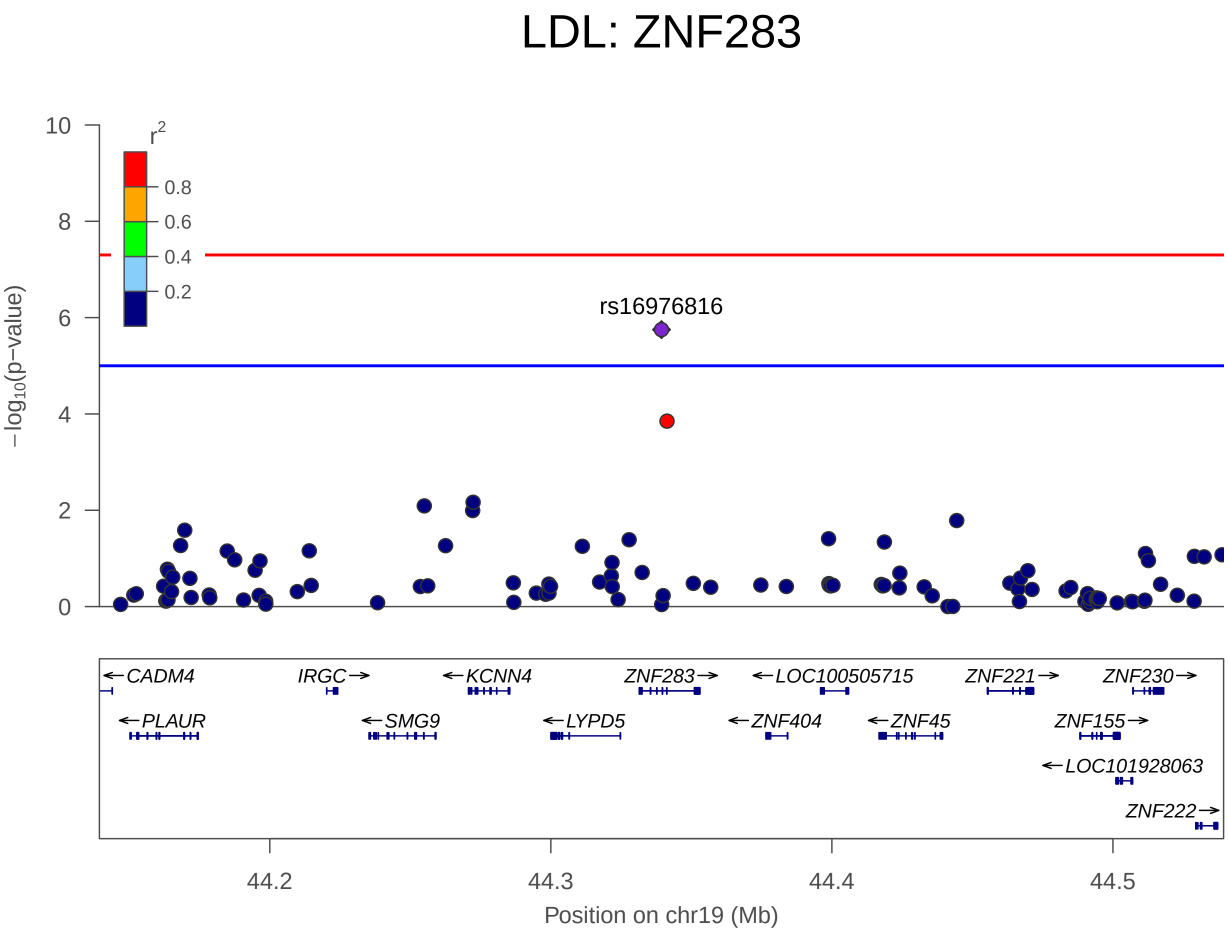


FIGURE S39. LDL on chromosome 19 positions 44139377-44539377

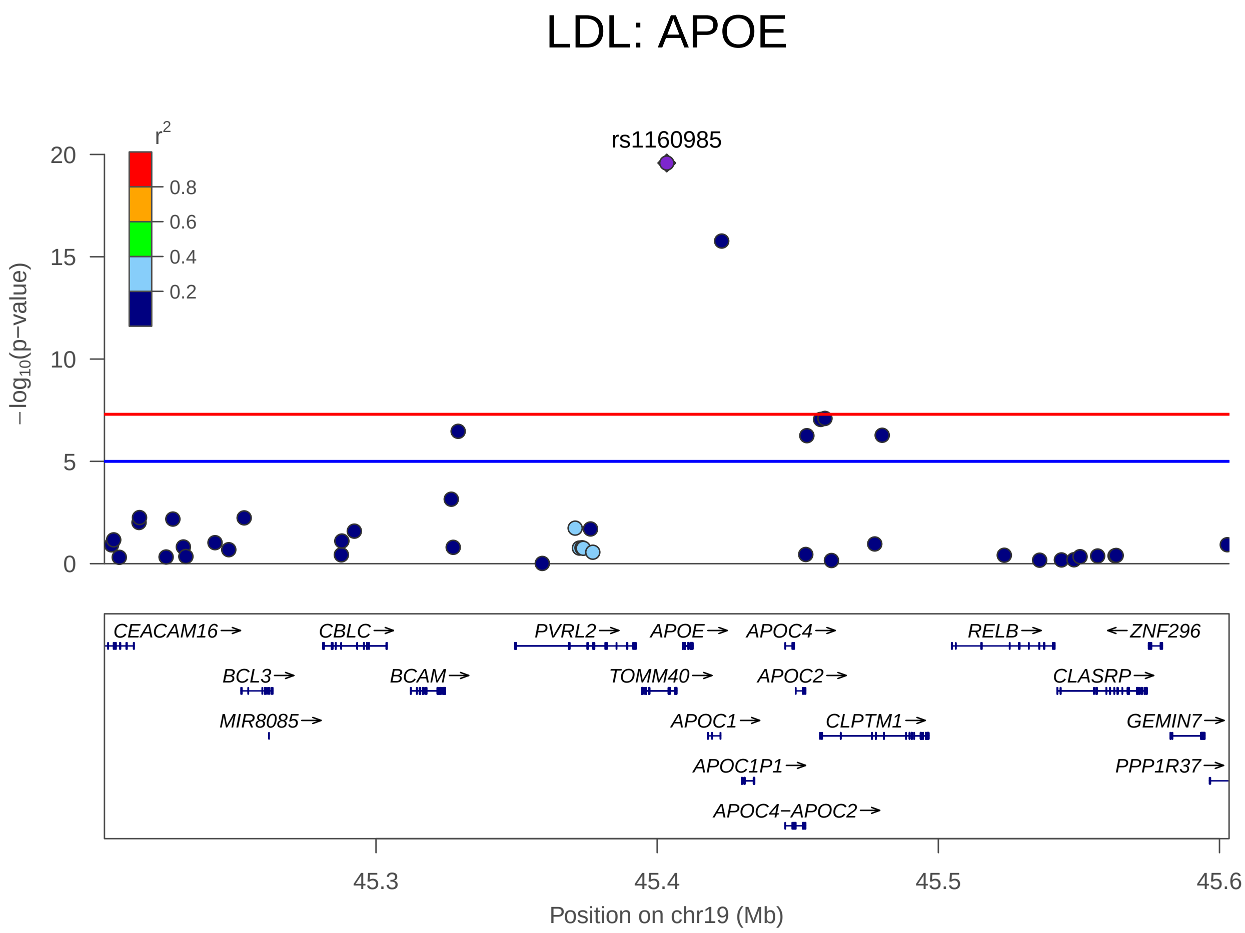


FIGURE S40. LDL on chromosome 19 positions 45203412-45603412

6.4. **Triglycerides LocusZoom plots.**

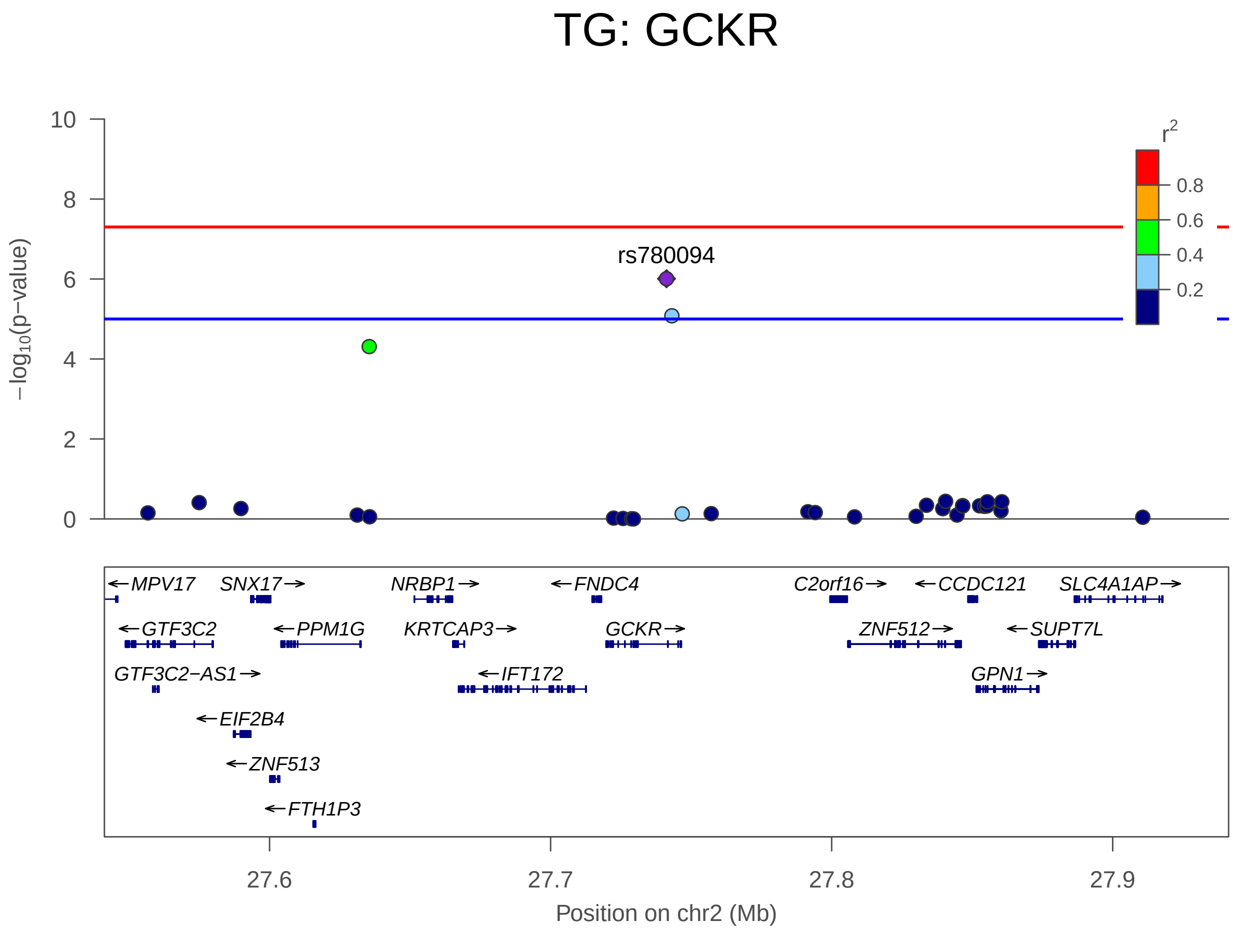


Figure S41. TG on chromosome 2 positions 27541237-27941237

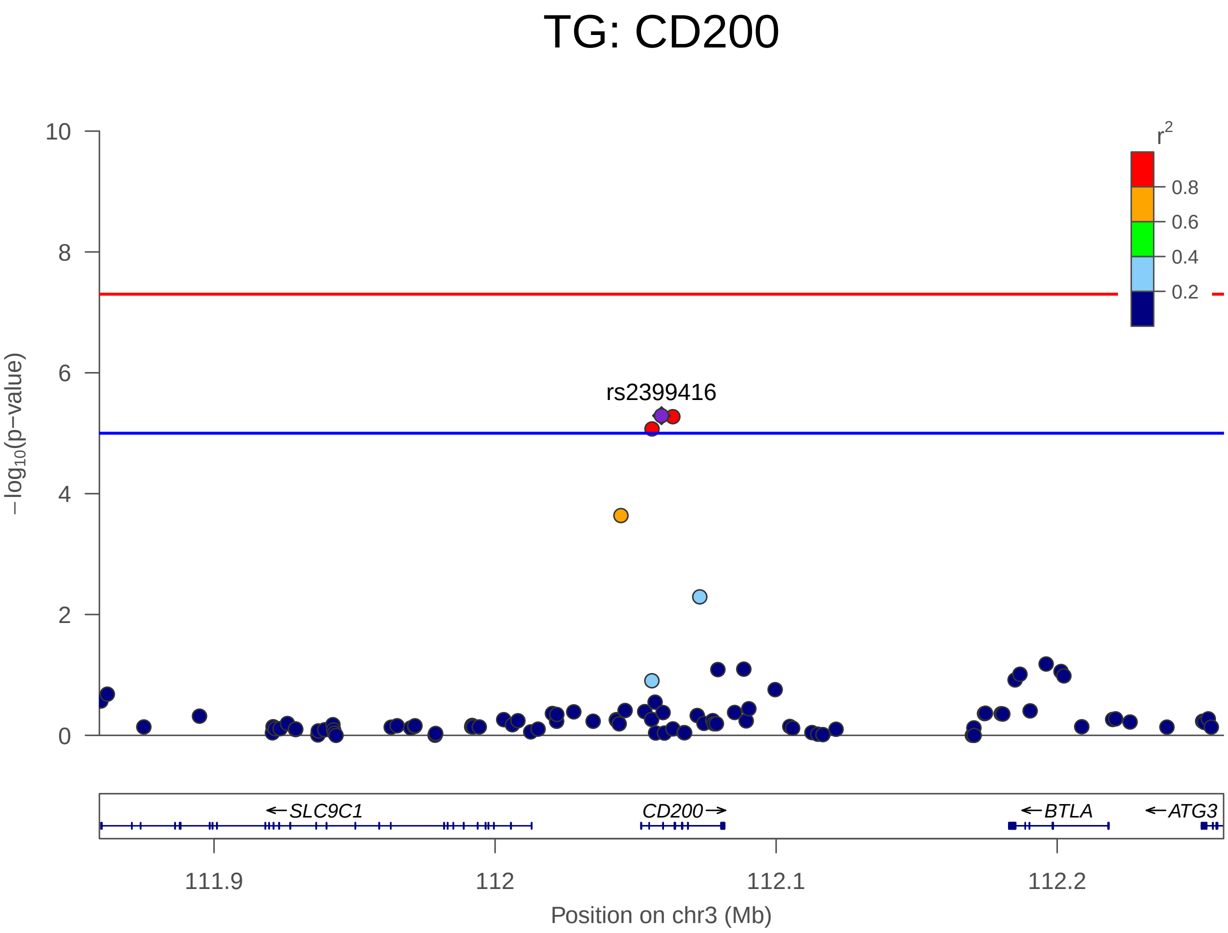


FIGURE S42. TG on chromosome 3 positions 111859213-112259213

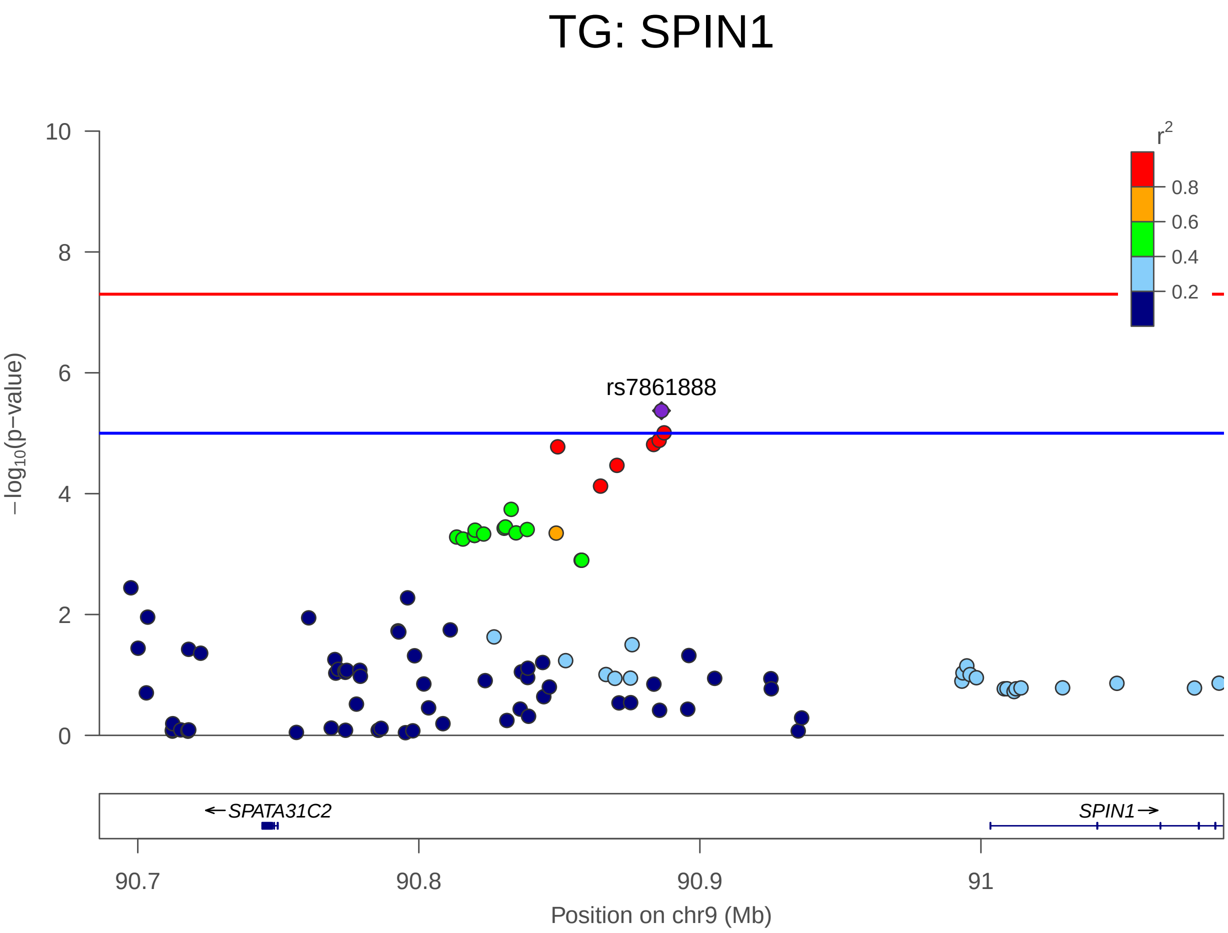


FIGURE S43. TG on chromosome 9 positions 90686340-91086340

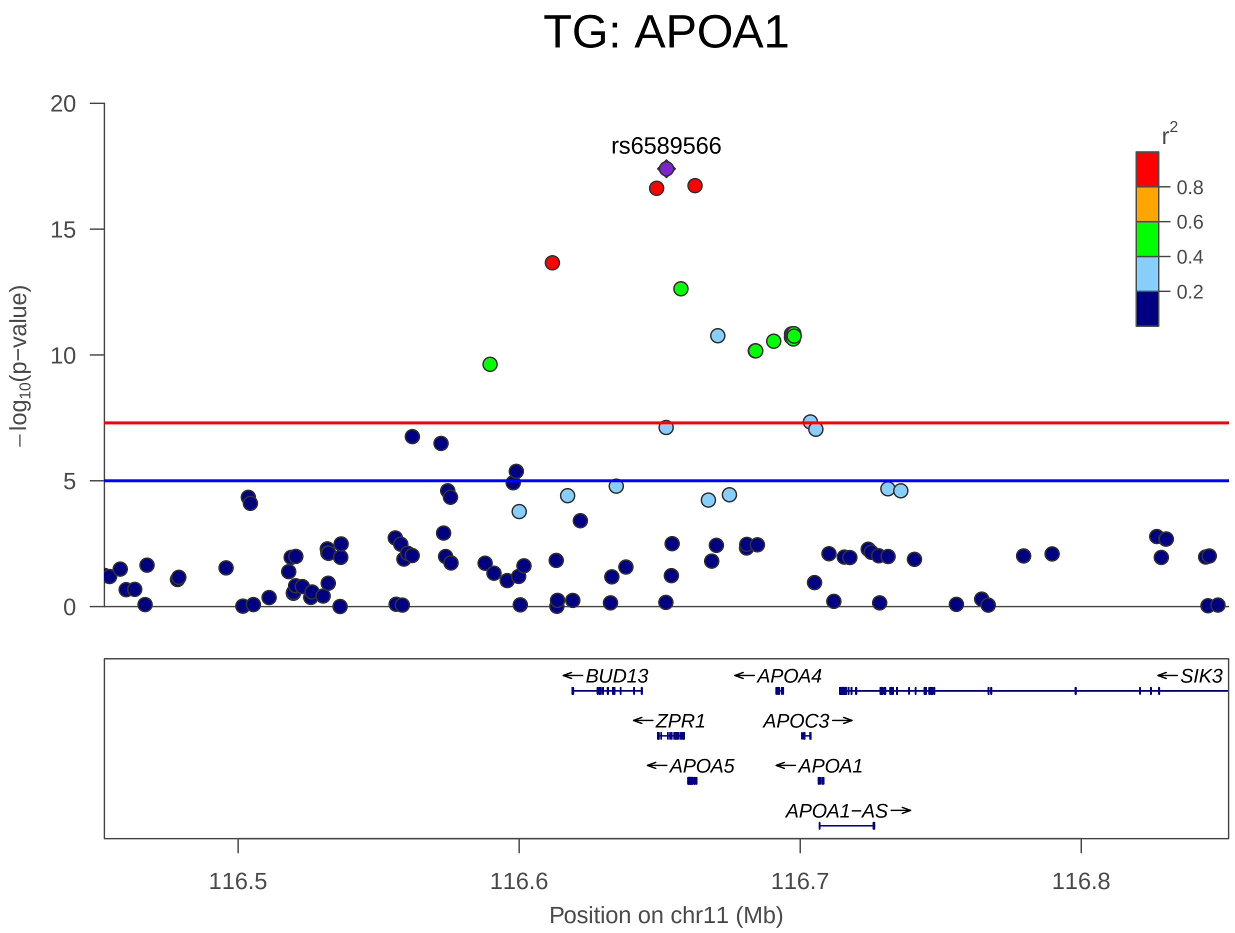


FIGURE S44. TG on chromosome 11 positions 116452423-116852423

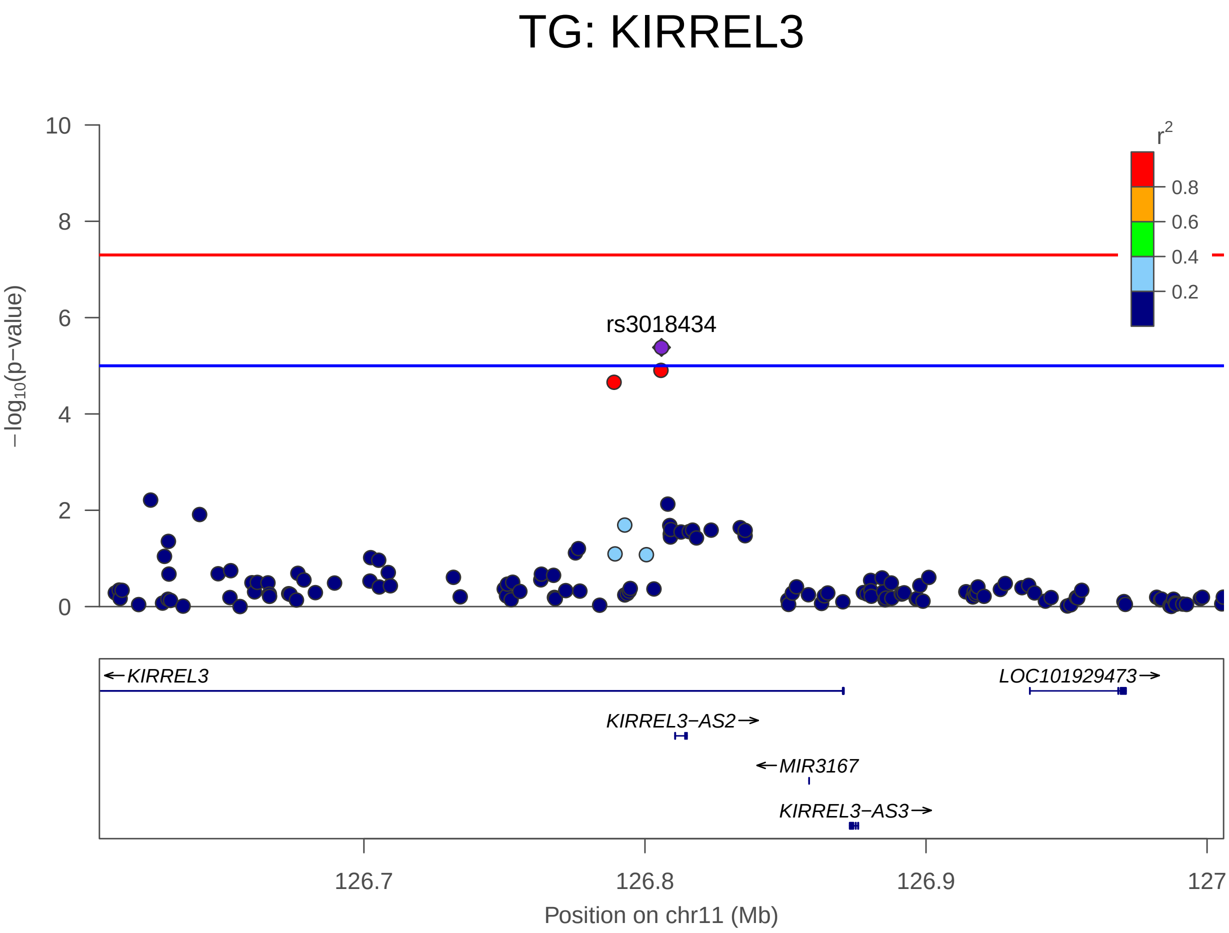


Figure S45. TG on chromosome 11 positions 126605881-127005881

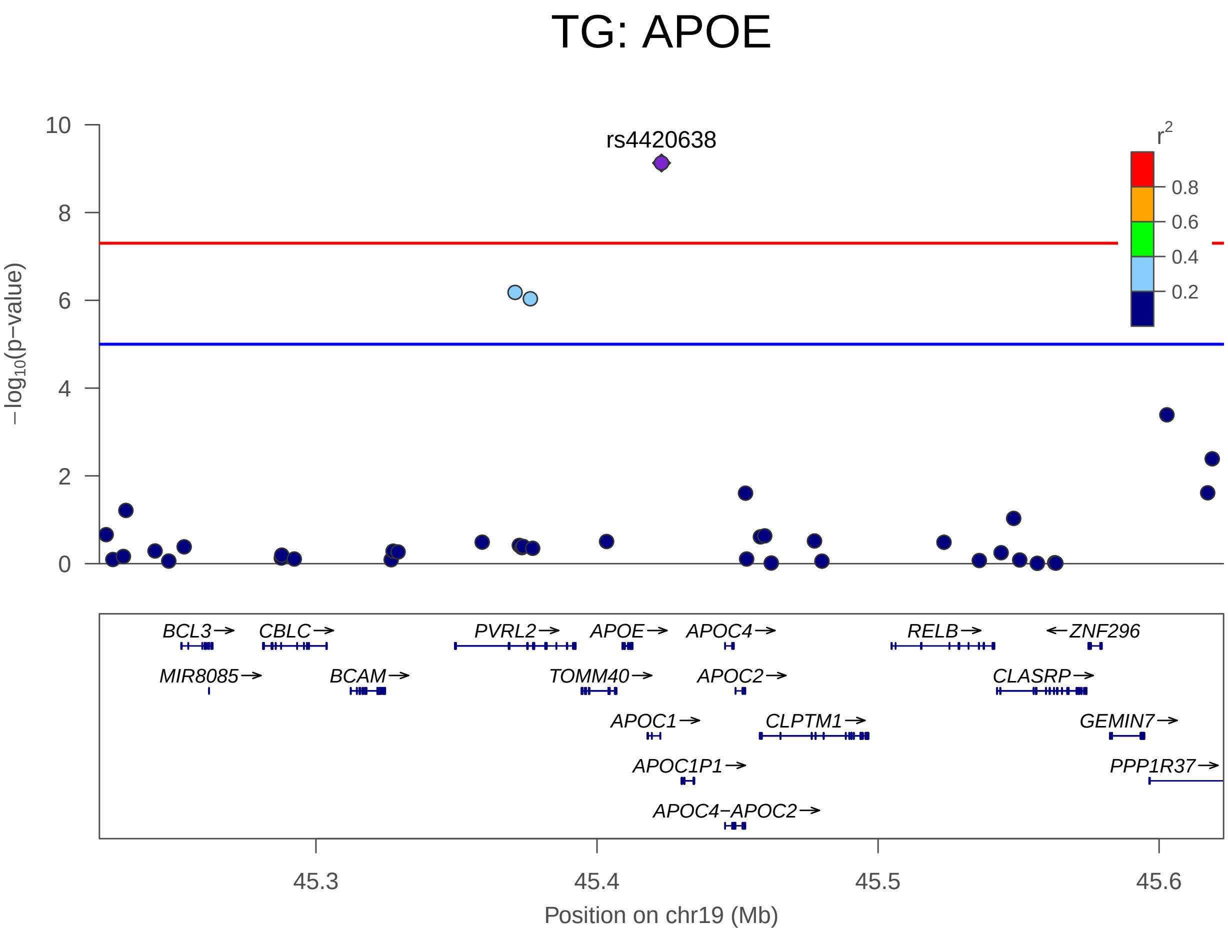


FIGURE S46. TG on chromosome 19 positions 45222946-45622946
